# HiBASIL: Hierarchical Bayesian Source Inference and Localization for Spatial Epidemiology

**DOI:** 10.64898/2026.07.28.741334

**Authors:** Fangfang Guo, Sharmodeep Bhattacharyya, Shirshendu Chatterjee, David Gent, Peter S. Ojiambo

## Abstract

1. Identifying spatial origins of biological invasions, disease outbreaks, or environmental contaminants is critical for timely intervention. However, existing methods struggle to resolve overlapping signals from multiple sources or account for extreme zero/one inflation in bounded data.
2. We developed HiBASIL (**Hi**erarchical **BA**yesian **S**ource **I**nference and **L**ocalization), a Bayesian framework that jointly infers source coordinates and mechanistic dispersal kernels from zero-one-inflated spatial observations. We systematically tested HiBASIL across 2,500 simulations spanning localized to long-distance dispersal regimes, various foci weight mixtures (0.5/0.5 to 0.95/0.05), and varying sample sizes (*N* = 50 to 500) to evaluate geometric sensitivity, spatial robustness, parameter recovery capability, and multi-focal resolution. We applied HiBASIL in tracking cucurbit downy mildew dispersal in field experiments including non-inoculated control, single-focus, and two-foci treatments arranged in a randomized complete block design. To demonstrate broad applicability, we validated HiBASIL with historical London cholera epidemic data to determine whether it could accurately localize the epicenter.
3. HiBASIL achieved 100% accuracy in kernel identification under correct model specification, 100% and 98% (2% predictive equivalence) in under- and over-parameterization scenarios, and maintained appropriate parsimony in null scenarios. Source localization was highly precise, with a median bias of 0.11 m, successfully resolving minority sources contributing only 5% of the observations. Importantly, the framework provides intrinsic self-diagnostics, signaling model complexity mismatch through posterior bimodality (under-parameterization) or parameter collapse (over-parameterization). HiBASIL is applicable in various epidemic scenarios, including diffused long-distance dispersal. Under an isotropic process, HiBASIL achieved sub-meter mean errors on two-source localization in cucurbit downy mildew field tests, effectively isolating transmission signals from landscape noise despite sparse sampling. Despite confounding influence of underlying network processes in the historical cholera data, the posterior localized the epicenter to within ∼ 33*m*, demonstrating portability beyond plant disease epidemiology.
4. In summary, HiBASIL can accurately localize multiple sources across various epidemic scenarios, extremely unbalanced foci mixtures, and sparse sampling conditions. HiBASIL also demonstrates wide applicability across agricultural and human disease epidemic systems. By providing an open-source Python implementation, HiBASIL enables rigorous inverse spatial inference for ecology, epidemiology, and environmental monitoring, transforming how discrete transmission sources are identified in complex landscapes.

## 1 Introduction

The identification of foci location from spatial disease patterns is fundamental to outbreak investigations, biosecurity, and invasion ecology. Whether tracking airborne spores, vector-borne pathogens, or invasive species, the ability to infer the location and intensity of unknown sources from observed gradients is essential for effective intervention and management (Nathan et al., 2012; Riley, 2007). Hierarchical Bayesian frameworks have proven effective in estimating dispersal kernels from ecological data (Clark et al., 1999; Banerjee et al., 2025), however, they typically require *a priori* knowledge of source locations. Although advances in blind source localization for discrete network topologies (Pinto et al., 2012) and heuristic spatial algorithms such as geographic profiling (Le Comber et al., 2011) have addressed specific contexts, a general statistical method for jointly inferring unknown source coordinates and mechanistic dispersal processes in continuous space is still lacking.

Observed spatial gradients rarely arise from a single isolated source but rather, they represent a superposition of overlapping dispersal kernels from multiple proximate foci (Liu, 2021; Rogers et al., 2019). This mixture process can be formalized as a weighted sum of individual kernels (Equation (1)), where the total intensity *κ*(*r*) at location *r* is the aggregate of *m* sources, each contributing a weight *p_i_* and kernel *κ_i_*.

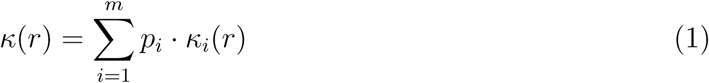

Decoupling these mixed signals (Figure 1A) is an ill-posed inverse problem (Tarantola, 2005). The central challenge here is one of identifiability (Luo et al., 2009). For example, a high-density patch in an observed gradient could arise from either the long-distance tail of a single, highly fecund source or the aggregate center of multiple weaker, proximate sources. If the true process involves multi-source superposition but is analyzed using a single-source model, inference will inevitably produce biased parameter estimates that confound the individual dispersal characteristics of each focus.

**Figure 1:**
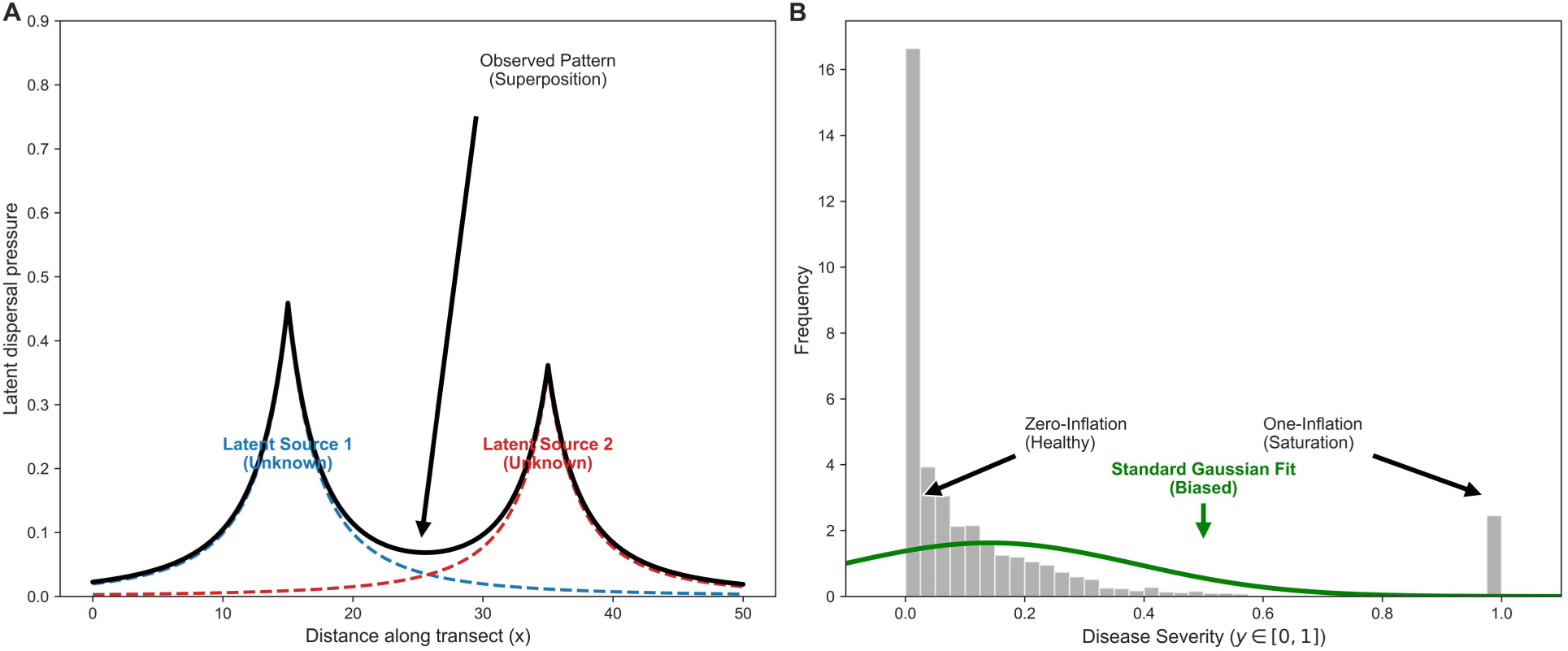
Conceptual framework and structural identifiability of HiBASIL. **A,** Spatial superposition. The observed spatial gradient (solid black line) arises from superposition of multiple cryptic sources with distinct, unknown dispersal kernels (dashed blue and red lines). **B,** Zero-One-Inflated Beta (ZOIB) likelihood. Empirical spatial data frequently exhibit extreme boundary inflation at 0 (structural zeros, e.g., uninfected hosts) and 1 (saturation). Standard continuous models (e.g., Gaussian models, green curve) generate impossible predictions, whereas HiBASIL explicitly decomposes these boundary masses.

This geometric challenge is compounded by a statistical problem, where an empirical intensity of infection is inherently bounded and zero-one-inflated (Figure 1B). Field data are frequently dominated by structural zeros (healthy) and structural ones (complete infection), creating distributions that violate standard statistical assumptions. Traditional approaches fail in distinct ways. Gaussian models produce physically impossible predictions outside [0, 1] (Warton and Hui, 2011). Ad hoc transformations (e.g., logit, arcsine) fail to properly account for boundary masses and obscure biological interpretation (Douma and Weedon, 2019). Even Beta regression, explicitly designed for proportional data, is undefined at boundary values (Ospina and Ferrari, 2012). While zero-one-inflated Beta (ZOIB) models have emerged to address boundary masses (Liu and Kong, 2015; Jensen et al., 2022), no existing framework integrates this bounded probabilistic structure with mechanistic multi-source spatial inference.

Here, we developed **HiBASIL** (**Hi**erarchical **BA**yesian **S**ource **I**nference and **L**ocalization), a hierarchical Bayesian framework that jointly infers source coordinates and mechanistic dispersal kernels from zero-one-inflated spatial observations. HiBASIL couples a ZOIB likelihood with a mixture of latent dispersal kernels, explicitly decomposing boundary masses (structural zeros and ones) from continuous infection intensity, and employs partial- or complete-pooling structures across experimental replicates to stabilize estimates in noisy or sparse datasets (Gelman and Hill, 2007). The framework assumes isotropic and radial spread as a common simplification of real epidemic behavior, though this may not hold under anisotropic dispersal, heterogeneous landscapes, or complex transmission networks. We validate HiBASIL through systematic simulation experiments and apply it to field experiments on cucurbit downy mildew and to the 1854 London cholera outbreak (Snow, John, 1936), to demonstrate its accurate source localization and appropriate parsimony across ecological and epidemiological contexts. HiBASIL is implemented as open-source Python software, providing a transferable tool for source localization across plant pathosystems and human epidemiology when disease spread can be assumed to be isotropic and radial.

## 2 Materials and Methods

### 2.1 Bayesian Inference of Dispersal Kernels Functions

#### Zero-One-Inflated Beta Distribution

Disease severity expressed as a continuous proportion (e.g., percentage of infected leaf area), is bounded on [0, 1] scale but cannot be modeled by the standard Beta distribution, which is defined only for the open interval (0, 1) and does not accommodate structural zeros (completely healthy plants, *y* = 0), or ones (completely diseased, *y* = 1) (Madden and Ojiambo, 2024; Stroup, 2015). We therefore model disease severity using the Zero-One-Inflated Beta (ZOIB) distribution (Equation (2)) (Liu and Kong, 2015), which models severity as a three-component mixture:

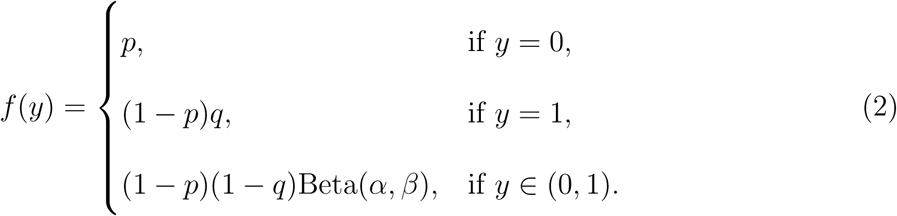

Where *p* denotes the probability of *y* = 0 (structural zero), *q* denotes the conditional probability *Pr*(*y* = 1 | *y ≠* 0) (structural one), and *α*, *β* are the shape parameters of the beta distribution when *y* ∈ (0, 1) (continuous component). Following Ferrari and Cribari-Neto (2004), we reparameterize the beta distribution using the mean 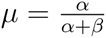 and precision parameter *φ* = *α* + *β*, where *µ* is the expected severity conditional on *y* ∈ (0, 1) and *φ* controls the concentration around the mean.

#### Spatial Dispersal Integration via Identity Links

Let *y_i_* (*y_i_* ∈ [0, 1]) denote the disease severity proportion at location *i* (*i* = 1*, . . ., m*), with ZOIB parameters *p_i_* ∈ [0, 1] (zero-inflation probability), *q* ∈ [0, 1] (one-inflation probability), *µ_i_* ∈ (0, 1) (mean of continuous component), and *φ >* 0 (precision parameter). Spatial dispersal kernels are integrated into *µ_i_* and *p_i_* through identity link functions that preserve parameter bounds without transformation (Liu and Eugenio, 2018).

##### Single Infection Focus

For a single infection focus, the mean parameter *µ_i_* is modeled as:

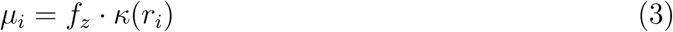

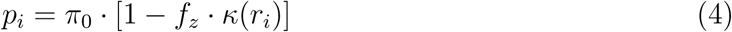

Where *f_z_* ∈ [0, 1] represents the focus intensity, and *κ*(*r_i_*) ∈ (0, 1] denotes the dispersal kernel at distance *r_i_* from the inoculum source, and *π*_0_ ∈ [0, 1] represents the baseline zero-inflation probability. Since *f_z_* ∈ [0, 1] and *κ*(*r_i_*) ∈ (0, 1], both *µ_i_* and *p_i_* remain bounded without transformation. To accommodate the beta distribution’s strict bounds (excluding exactly 0 and 1), boundary values are clipped using a small numerical tolerance *ɛ* = 10*^−^*^12^ during likelihood evaluation. Both one-inflation probability *q* and the precision parameter *φ* are constant across all observations. At each location *i*, the dispersal kernel *κ*(*r_i_*) governs the spatial gradient of ZOIB mixture weights, where locations near the inoculum source have low zero-inflation probability and high mean severity, while distant locations have high zero-inflation probability reflecting inoculum absence.

##### Multiple Foci Extension

For multi-foci, the framework extends through weighted additive contributions:

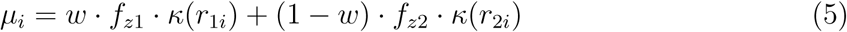

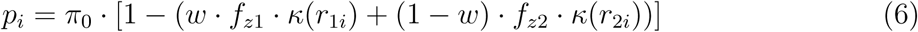

Where *w* ∈ [0, 1] represents the relative contribution weight between foci (shared across replicates), *f_z_*_1_ ∈ [0, 1] and *f_z_*_2_ ∈ [0, 1] denote the intensities of the two foci (replicates-specific), and *r*_1*i*_, *r*_2*i*_ represent distances from the observation point to each focus.

The natural bounds preservation extends to the multiple foci case. Since *w* ∈ [0, 1], *f_z_*_1_*, f_z_*_2_ ∈ [0, 1], and *κ*(*r*_1*i*_)*, κ*(*r*_2*i*_) ∈ (0, 1], the weighted sum *µ_i_* = *w* · *f_z_*_1_ · *κ*(*r*_1*i*_) +(1 − *w*) · *f_z_*_2_ · *κ*(*r*_2*i*_) necessarily remains in [0, 1]. This convex combination preserves all the interpretability and computational advantages of the identity link framework.

#### Dispersal Kernel Types

We implement three biologically motivated spatial dispersal kernels that capture different dispersal mechanisms: i) Exponential kernel representing passive dispersal with constant decay rate; ii) Gaussian kernel modeling diffusion-like processes with accelerating decay; and iii) Power-law kernel capturing long-distance dispersal with polynomial decay (Karisto et al., 2023). These kernels are defined as Equations (7), (8), and (9), respectively:

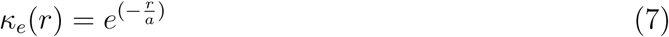

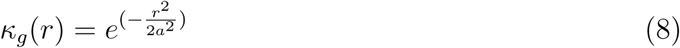

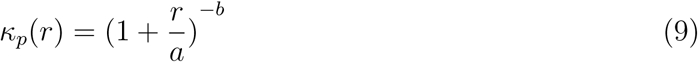

Where *a >* 0 is the scale parameter and *b >* 0 is the exponent parameter (power-law only).

#### Likelihood Function

Given the ZOIB parameterization (Equation 2 and dispersal kernel linkages (equations 3–6), the likelihood for observation *i* is:

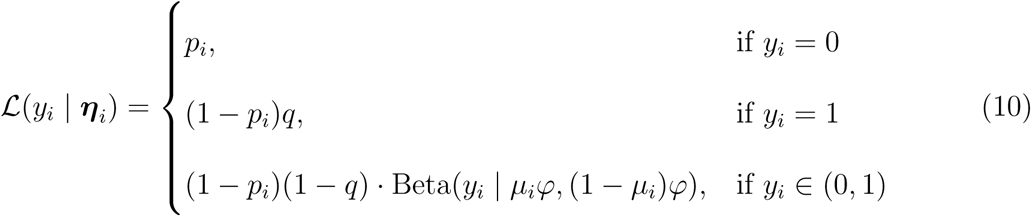

where ***η****_i_* = (*p_i_, q, µ_i_, φ*) represents the location-specific and shared parameters, and Beta(·) denotes the beta probability density function parameterized by the shape parameter *α_i_* = *µ_i_φ* and *β_i_* = (1 − *µ_i_*)*φ*. To preserve model parsimony and ensure parameter identifiability, both the conditional one-inflation probability *q* and precision parameter *φ* are treated as globally constant across. Because the zero-inflation probability *p_i_* varies spatially, the marginal probability of observing complete saturation (Pr(*y_i_* = 1) = (1 − *p_i_*)*q*) inherently retains a spatial gradient dictated by the dispersal kernel.

The full dataset likelihood across *n* observations is computed as:

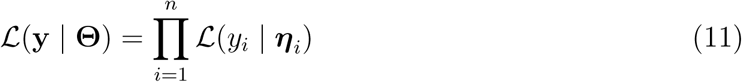

where **Θ** includes dispersal parameters (scale *a* and exponent *b*), source location(s), replicate-specific focus intensities (*f_z_*), and shared parameters (focus weight *w*, *π*_0_, *q*, *φ*).

### 2.2 Bayesian Implementation and Computational Framework

#### Prior Specification

We employed weakly informative priors to allow the data to dominate posterior inference while constraining parameters to biologically plausible ranges (Table 1). Where focus locations were unknown, we assigned normal priors centered on approximate field positions, while where known, their coordinates were fixed.

**Table 1:** Simulated parameters and priors specification for the test of geometric sensitivity, spatial robustness, and multi-focal resolution.

| Parameter | True Value (Simulation) | Prior Distribution (Estimation) |
| --- | --- | --- |
| *Global (ZOIB)* |  |  |
| Zero-inflation ($\pi_0$) | 0.30 | Beta(2, 2) |
| One-inflation ($q$) | 0.01 | Beta(1, 10) |
| Precision ($\varphi$) | 18 | Gamma(2, 0.1) |
| *Dispersal* |  |  |
| Scale ($a$) | 5 | Gamma(2, 0.4) |
| Exponent ($b$) | 2 | Gamma(3.6, 2.0) |
| *Focus 1* |  |  |
| x-coordinate ($f_{x1}$) | 0 | $\mathcal{N}(5.0, 5.0^2)$ or fixed |
| y-coordinate ($f_{y1}$) | 0 | $\mathcal{N}(25.0, 20.0^2)$, $\mathcal{N}(5.0, 20.0^2)$ or fixed |
| Severity ($f_{z1}$) | 0.6 | Beta(1.0, 1.0) or fixed |
| *Focus 2* |  |  |
| x-coordinate ($f_{x2}$) | 0 | $\mathcal{N}(5.0, 5.0^2)$ or fixed |
| y-coordinate ($f_{y2}$) | 50 | $\mathcal{N}(30, 20.0^2)$ or fixed |
| Severity ($f_{z2}$) | 0.80 | Beta(1.0, 1.0) or fixed |
| *Mixture* |  |  |
| Weight ($w$) | 0.60 | Beta(2, 2) |

Bounded proportion parameters, such as focus intensities (*f_z_*) and mixture weight parameter (*w*), were assigned standard or symmetric Beta priors (Beta(1, 1) or Beta(2, 2)) to reflect uniform or neutral prior assumptions without favoring specific foci. Similarly, the baseline zero-inflation *π*_0_ received a symmetric prior Beta(2, 2). Conversely, the one-inflation probability *q* was assigned a strongly right-skewed prior Beta(1, 10), reflecting the biological expectation that complete infection is relatively rare. Finally, scale, exponent, and precision parameters (*a*, *b*, and *φ*) were assigned Gamma priors to strictly enforce positivity while allowing flexibility in variance and the spatial tail shape. Unless otherwise noted for specific validation controls, these priors were used across all analyses.

#### Custom ZOIB Distribution and MCMC Sampling Strategy

We implemented a customized ZOIB distribution within PyMC (version 5.25.1) (Abril-Pla et al., 2023). The custom distribution handles the three-component mixture through a conditional log-likelihood computation. The PyMC implementation employs automatic differentiation through PyTensor (Yoo et al., 2010) to efficiently compute gradients of this piecewise likelihood function, enabling gradient-based sampling algorithms. Posteriors were sampled using the No-U-Turn Sampler (NUTS) (Hoffman and Gelman, 2014), with 4 independent Markov chains, each producing 8,000 post-warmup draws after 4,000 tuning iterations (32,000 total samples). The target acceptance probability was set to 0.85 to balance computational efficiency with thorough exploration of the posterior (Hoffman and Gelman, 2014). Chain initialization employed Automatic Differentiation Variational Inference (ADVI) or jitter with diagonal covariance adaptation, providing improved starting values and reducing warm-up time. All analyses used fixed random seeds to ensure computational reproducibility.

#### Hierarchical (Mixed-Effects) Structure

The ZOIB distribution parameters (*π*_0_, *q*, *φ*) and the mixture weight parameter (*w*) were treated as fixed effects shared across all replicates within each treatment, representing population-level disease characteristics. Dispersal parameters (scale *a* and exponent *b*), focus coordinates (*f_x_* and *f_y_*, when unknown), and focus intensity (*f_z_*) can either partially or completely pooled depending on similarity of disease patterns across the replicates. Generally, focus coordinates and disease intensities are replicate-specific and they were treated as random effects varying among replicates, capturing block-to-block variation in inoculum characteristics. This Hierarchical structure enables simultaneous inference across multiple replicates, borrowing information across replicates to stabilize global parameter estimation while explicitly accounting for replicate-specific biological variation.

### 2.3 Model Evaluation and Diagnostic Criteria

#### Convergence Diagnostics

MCMC convergence was assessed using the rank-normalized potential scale reduction factor *Ȓ* (Vehtari et al., 2021), with *Ȓ* ≤ 1.01 and effective sample size *ESS >* 400 per chain as thresholds for adequate mixing and sampling efficiency.

#### Parameter Recovery

We quantified parameter recovery across simulations using posterior point-estimate bias and 95% credible interval coverage against known true values.

#### Posterior Predictive Checks

The adequacy of the model was evaluated by posterior predictive checks, comparing observed data against posterior predictive draws. Given the three-part nature of ZOIB distribution, we employed a component-wise validation strategy to avoid masking compensating errors. First, we isolated and compared the observed versus predicted proportions of structural zeros (*y* = 0) and ones (*y* = 1). Second, we calculated *R*^2^ and root mean squared error (RMSE) for observations within the (0, 1) interval. Global predictive accuracy was summarized using *R*^2^, RMSE, and mean absolute error (MAE).

#### Model Selection

Leave-One-Out Cross-Validation (LOO-CV) was used to compare models via the ArviZ library (Kumar et al., 2019), which estimates out-of-sample predictive performance without refitting (Vehtari et al., 2017). We computed the expected log-pointwise predictive density (ELPD_loo_) and effective number of parameters (*p_loo_*) using LOO-CV. Model preference was determined by ELPD differences (ΔELPD); differences exceeding twice the standard error were considered evidence of a significant predictive gain.

### 2.4 Simulation Study and Validation Framework

We evaluated the HiBASIL framework through a series of simulation experiments targeting kernel discrimination, source location recovery, parameter recovery, and structural robustness under varied spatial complexities, mixture configurations, and data constraints.

#### Data Simulation

Synthetic disease datasets were generated with known spatial structure using a ZOIB generative process, producing completely healthy (*y* = 0), completely infected (*y* = 1), and partial infected (0 *< y <* 1) observations. For spatial scenarios, mean severity *µ* and the zero-inflation probability *p* were functions of the Euclidean distance from one or more infection foci via dispersal kernels (Equations (3), (4), (5), and (6)). For non-spatial Null scenarios testing inference integrity and specificity, zero-inflation probability *p* and mean severity *µ* were spatially constant, producing a spatially random background without gradient. Regardless of the spatial-dependent or spatial-independent process, the observed severity (*y_i_*) at each point *i* was drawn from the three-component mixture according to the ZOIB distribution (Equation (10)). We utilized these generative processes to generate 100 independent datasets for each testing scenario.

#### Geometric Sensitivity, Spatial Robustness, and Multi-focal Resolution

We first established a baseline for geometric sensitivity by fixing the source coordinates at their true values, isolating the ability of HiBASIL to differentiate kernel shapes via the LOO-CV. Subsequently, we evaluated spatial robustness (the ability to resolve source locations under varying levels of structural complexity) and multi-focal resolution by treating source coordinates as unknown parameters, using the weakly informative priors specified in Table 1. The performance of Bayesian model estimations was evaluated using Mean Bias, 95% Coverage, *Ȓ*, and the bulk-ESS.

#### Parameter Recovery and Model Robustness

We evaluated HiBASIL’s capacity to recover parameters across four epidemiological scenarios (Table S1). The **Base Scenario** represents a moderate dispersal reach with standard decay (*a* = 5*, b* = 2), serving as the reference for initial simulations. The **Narrow and Steep** scenario (*a* = 1*, b* = 3) represents a highly localized spread with a rapid drop-off in infection severity. In contrast, the **Wide and Steep** scenario (*a* = 15*, b* = 3) represents an extended dispersal reach but with a sharply defined spatial boundary. Finally, the **Wide and Shallow** scenario (*a* = 15*, b* = 1.5) represents the long-distance spread characterized by a heavy “fat-tail” distribution.

We conducted 100 simulations per case using the scenario-specific, weakly informative priors detailed in Table S1. More importantly, all estimations were performed under the condition of unknown infection source locations to simulate realistic monitoring challenges. We evaluated the framework’s capacity to distinguish between competing kernel geometries via LOO-CV, predictive accuracy by posterior predictive checks (PPC), and reliability of parameter estimation by evaluating mean bias, 95% coverage, *Ȓ*, and bulk-ESS.

#### Model Inference across Varied Foci Mixture Weights

To evaluate the HiBASIL’s inference robustness under varying degrees of source contribution, we performed 100 simulations for each of several distinct mixture weight configurations. Mixture proportions of (0.5, 0.5), (0.6, 0.4), (0.8, 0.2), and (0.95, 0.05) were evaluated with unknown foci locations, testing spatial and parametric recovery simultaneously. All other parameters and prior specifications were as specified in Table 1, with Beta(2, 2) assigned to mixture weights to remain uninformative. The framework’s capacity to distinguish between competing kernel geometries, predictive accuracy, and accuracy of parameter estimation were evaluated across all mixture weight configurations. Furthermore, Pearson correlation analysis of parameter estimation biases were conducted to explore the underlying structural confounding.

#### Model Misspecification and Structural Robustness

We conducted two types of model misspecification tests to quantify the inferential consequences of structural mismatch. First, under-parameterization applied a single-focus model to two-foci data, assessing bias in the estimation of the focus locations and the potential blurring of dispersal parameters (scale *a* and exponent *b*), as the model attempts to consolidate two distinct spatial signal into a single kernel. Second, over-parameterization applied two-foci model to single-focus data, monitoring whether the redundant focus collapses to a negligible mixture weight (*w*). Both tests used the same datasets generated for the corresponding correctly specified scenarios, allowing direct quantification of inferential cost relative to correctly specified benchmarks. The model’s performances on kernel selection, predictive accuracy, and parameter recovery were evaluated by LOO-CV, posterior predictive checks, and mean bias, 95% coverage, *Ȓ*, and bulk-ESS, respectively. Furthermore, posterior diagnostic was performed by using the find peaks algorithm in the scipy.signal module (Virtanen et al., 2020) to detect peaks from the histogram of posterior distribution of fy.

#### Model Inferential Performance Across Sample Sizes

We evaluated inference performance at sample sizes *N* = 50, 100, 250, and 500 under unknown Single-focus and Two-foci spatial complexity. For each scenario, 100 simulations were generated following the simulation and prior settings in Table 1. Performance was assessed across three criteria: kernel discrimination (Δ*ELPD_loo_* distributions relative to the 2 × *SE* equivalence zone), predictive accuracy (posterior predictive checks (PPC) for disease severity predictions and the calibration of zero- and one-inflation components), and estimation precision (bias, coverage, and convergence (*Ȓ*) across spatial and ZOIB parameters).

### 2.5 Empirical Validation via Field-scale Cucurbit Downy Mildew Experiments

We applied HiBASIL to characterize the dispersal kernels of cucurbit downy mildew across experimental treatments. The treatments were: i) non-treated control, ii) single-focus inoculated once, iii) single-focus inoculated twice, iv) two-foci inoculated once, and v) two-foci inoculated twice. Six disease assessment surveys were conducted over the experimental periods. The third survey was selected for modeling as it captured peak establishment of primary dispersal gradients (Supplementary Note 2 S5). To account for the Randomized Complete Block Design (RCBD) for the experiment, we incorporated block as a random effect, thereby isolating environmental noise from the biological dispersal signal. Disease severity for each treatment was fitted to three candidate dispersal kernels, Power Law, Exponential, and Gaussian kernels. Model selection was performed via LOO-CV. For the non-treated control (CK), we evaluated a null model (no spatial effect) against single-focus model and two-foci models to verify the absence of artificial inoculation source (Table S2).

To assess the model’s spatial sensitivity and the influence of prior knowledge on parameter estimation, we performed the analysis using two distinct prior configurations: informative priors, which fixed foci locations (*f_x_*, *f_y_*) to known experimental coordinates to maximize the precision of the estimated kernel parameters and the precision parameter (*φ*); and weakly informative priors, which assigned broad priors to foci locations to evaluate source recovery from the field survey data.

Dispersal parameter priors were specified hierarchically to account for between-replicate (row) variation within each treatment. When preliminary analysis indicated substantial heterogeneity in gradient shapes across replicates, we employed partial pooling for the scale parameters. Under this approach, row-specific parameters were drawn from population-level distributions with hyperparameters estimated from the data: log(scale) ∼ N (log(scale*_µ_*), scale*_σ_*), implemented via non-centered parameterization for computational efficiency to avoid pathological sampling geometries (Table S3). In contrast, when disease gradients exhibited similar patterns across rows within a treatment, we used complete pooling, wherein all rows shared common scale and exponent parameters. For two-foci configurations, each replicate contained distinct infection sources (Focus 1 and Focus 2); the focus-specific dispersal parameters (scales *a*_1_, *a*_2_; exponents *b*_1_, *b*_2_) were globally pooled across all replicates to ensure statistical stability and capture distinct dispersal patterns. Similarly, other unobserved spatial features, such as unknown source coordinates (*f_x_*, *f_y_* for single focus treatments; *f_x_*_1_, *f_y_*_1_, *f_x_*_2_, *f_y_*_2_ for two-foci treatments), were estimated as shared population-level parameters across all replications within a treatment.

Posterior distributions were sampled using NUTS with 4 chains of 8,000 post-warmup draws following a 4,000-iteration warmup period. Convergence was monitored via the potential scale reduction factor (*Ȓ* ≤ 1.01) and bulk effective sample size (ESS).

### 2.6 Empirical Evaluation: Spatial Modeling of Historical Cholera Outbreak Data

#### Data and Study Context

We validated HiBASIL’s source localization capabilities using mortality records from the 1854 London cholera epidemic (*N* = 1852)(Center for Spatial Data Science, 2021). The established role of the Broad Street pump as the contamination source provides unambiguous ground truth, while waterborne transmission via London’s street network fundamentally violates radial dispersal assumptions, offering a stringent test of robustness under mechanistic misspecification. Death counts were normalized to maximum recorded mortality, converting raw incidence into proportional intensity values on [0, 1] scale.

#### Competing Hypotheses

We evaluated three hierarchical hypotheses under varying prior specifications (Table S4).

*H*_0_ **(Null):** Uniform risk distribution across the study area, representing random background mortality with no localized point-source.
*H*_1_ **(Blind Source Discovery):** Single-focus model treating source location as an unknown latent parameter with weakly informative priors (*σ* = 250*m*) centered on the study area’s geometric centroid to estimate outbreak scenarios where contamination sources are unknown.
*H*_2_ **(Parsimony Stress Test):** Two-foci model with highly informative priors (*σ* = 2*m*) centered on both the Broad Street and Little Marlborough Street pumps, testing whether the framework appropriately rejects unnecessary complexity when strongly guided toward a plausible two-foci alternative. The Little Marlborough Street pump (≈ 211*m* north of Broad Street) was historically attributed to resident preference for Broad Street water rather than pump contamination (Snow, John, 1855).

Exponential, Gaussian, and Power Law kernels were evaluated under *H*_1_ and *H*_2_, with LOO-CV used for model selection.

#### Spatial Visualization

To visualize the spatial distribution of outbreak intensity and quantify model uncertainty, posterior predictive risk maps and coefficient of variation (*σ/µ*) surfaces for the single-focus and two-foci models were generated in ArcGIS Pro 3.6 (Esri, Redlands, CA) using the historical Soho street shapefile (Li, 2017) as a basemap. For each model configuration, a continuous risk surface was produced by kernel density estimation (KDE) of the posterior predictive point layer, weighted by the predicted intensity at each location. The search radius (85.21 m and 86.37 m for single-focus and two-foci, respectively) was determined by the ArcGIS Pro Kernel Density default algorithm, a spatial adaption of Silverman’s rule of thumb (Silverman, 1998). To delineate the core of the outbreak’s risk surface, 50% and 95% highest density regions (HDRs) (Hyndman, 1996) were calculated from the continuous density surface using a custom volume-integration approach implemented via the arcpy and numpy Python Libraries. The continuous raster grid was converted into a two-dimensional numerical array and sorted in descending order of pixel density. We calculated the cumulative sum of this sorted array to find the unique pixel value threshold, *C_α_*, that bounds (1 − *α*) × 100% of the total integrated density mass (Equation (12)):

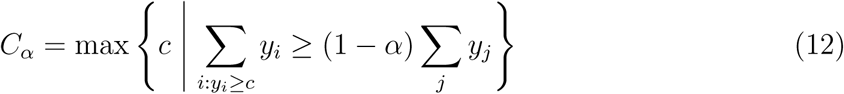

where *y_i_* represents individual pixel values, and *α* = 0.05 and 0.50 for the 95% and 50% contours, respectively. These extracted threshold values were then utilized in the ArcGIS Pro Contour List tool to map the geographical boundaries of the outbreak’s core risk. Parameter uncertainty across space was mapped by calculating the coefficient of variation (*σ/µ*) at each sampling data point using the posterior predictive mean and standard deviation. These point-level uncertainties were subsequently interpolated into a continuous uncertainty surface using inverse distance weighting (IDW).

#### Prior Sensitivity Analysis

We assessed the precision of localization across prior uncertainty levels by varying the standard deviation of single-focus priors centered on the Broad Street pump from *σ* = 5*m* (tight spatial constraint, building level certainty) to *σ* = 250*m* (loose spatial constraint, blind discovery scenario), evaluating all three dispersal kernels at each prior width.

## 3 Results

### 3.1 Performance of HiBASIL on Simulated Data

#### Framework Validation on Null Spatial Conditions

Under null spatial condition, HiBASIL correctly simplified to a non-spatial null state (*M_null_*) in the absence of a spatial signal. Model comparison via LOO-CV indicated that the null hypothesis was either the top-ranked model or predictively equivalent to the best model (|Δ*ELPD_loo_*| *<* 2 × *SE*) in 80% of trials. Furthermore, dispersal parameters remained strictly prior-driven in the absence of a likelihood signal, exhibiting a 100% success rate in maintaining high distributional overlap (*>* 94%). Full details of the control validation procedure and prior-to-posterior shift analysis are provided in **Supplementary Note 1 S4**.

#### Geometric Sensitivity and Multi-focal Resolution

We generated 100 simulations from a Power Law dispersal kernel and estimated each under three competing kernel geometries (Exponential, Gaussian, and Power Law). The LOO-CV identified the Power Law kernel in 100% simulations under both the known single-focus and two-foci scenarios (Figure 2A&B). Furthermore, the predictive density difference (Δ*ELPD_loo_*) for the Power Law kernel consistently exceeded the 2 × *SE* threshold for predictive equivalence across all configurations.

**Figure 2:**
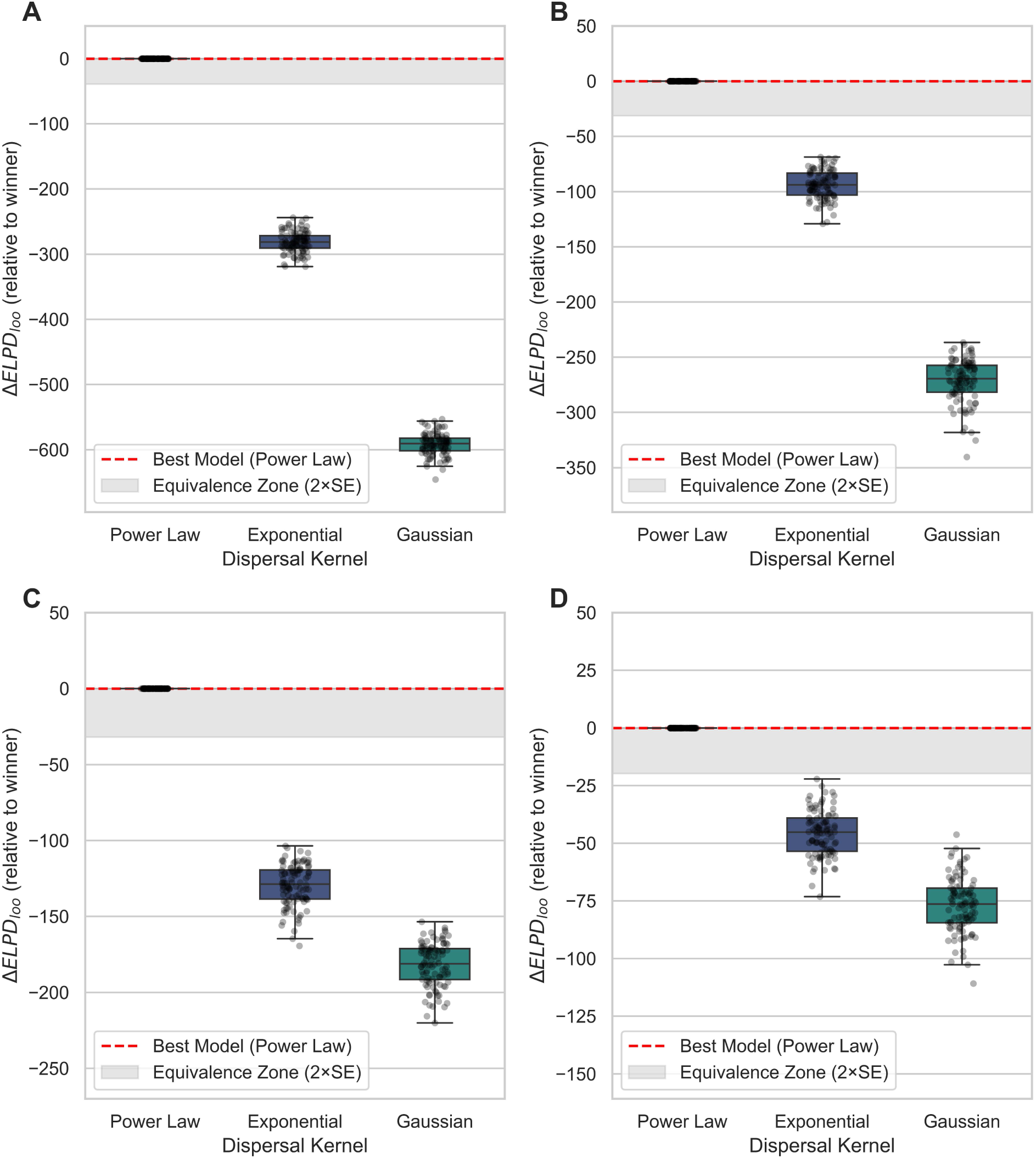
Model selection sensitivity and structural identifiability of dispersal kernels across known and unknown focus location scenarios. Distribution of Δ*ELPD_loo_* relative to the top-ranked model across 100 simulations for (A) known single-focus, (B) known two-foci, (C) unknown single-focus, and (D) unknown two-foci scenarios. The Power Law kernel was correctly identified in 100% of 400 simulations, with ΔELPD_loo_ consistently exceeding the 2 × *SE* threshold for predictive equivalence across all configurations.

Representative model fits for four replicates under known single-focus and two-foci scenarios are shown in Figure 3A–H, with posterior predictive mean accurately recovered the spatial distribution of infection severity and 95% CI effectively capturing the stochastic variation. Posterior predictive check (PPC) showed a high recovery accuracy for both aggregate and component-specific metrics (Figure 4). For the single-focus scenario, mean overall *R*^2^ = 0.47 ± 0.07 and continuous component *R*^2^ = 0.82 ± 0.01; for the two-focus scenario, mean overall *R*^2^ = 0.27 ± 0.06 and continuous component *R*^2^ = 0.61 ± 0.02 (Figure 4). Zero- and one-inflation proportions aligned closely with the 1:1 identity line across both scenarios. Dispersal kernels and ZOIB parameters were recovered with negligible bias and high coverage ≥ 93% (Table 2), and all MCMC chains converged (*Ȓ* = 1.00). Scale parameters in the two-foci scenario exhibited slightly higher bias relative to the single-focus case due to increased model complexity.

**Figure 3:**
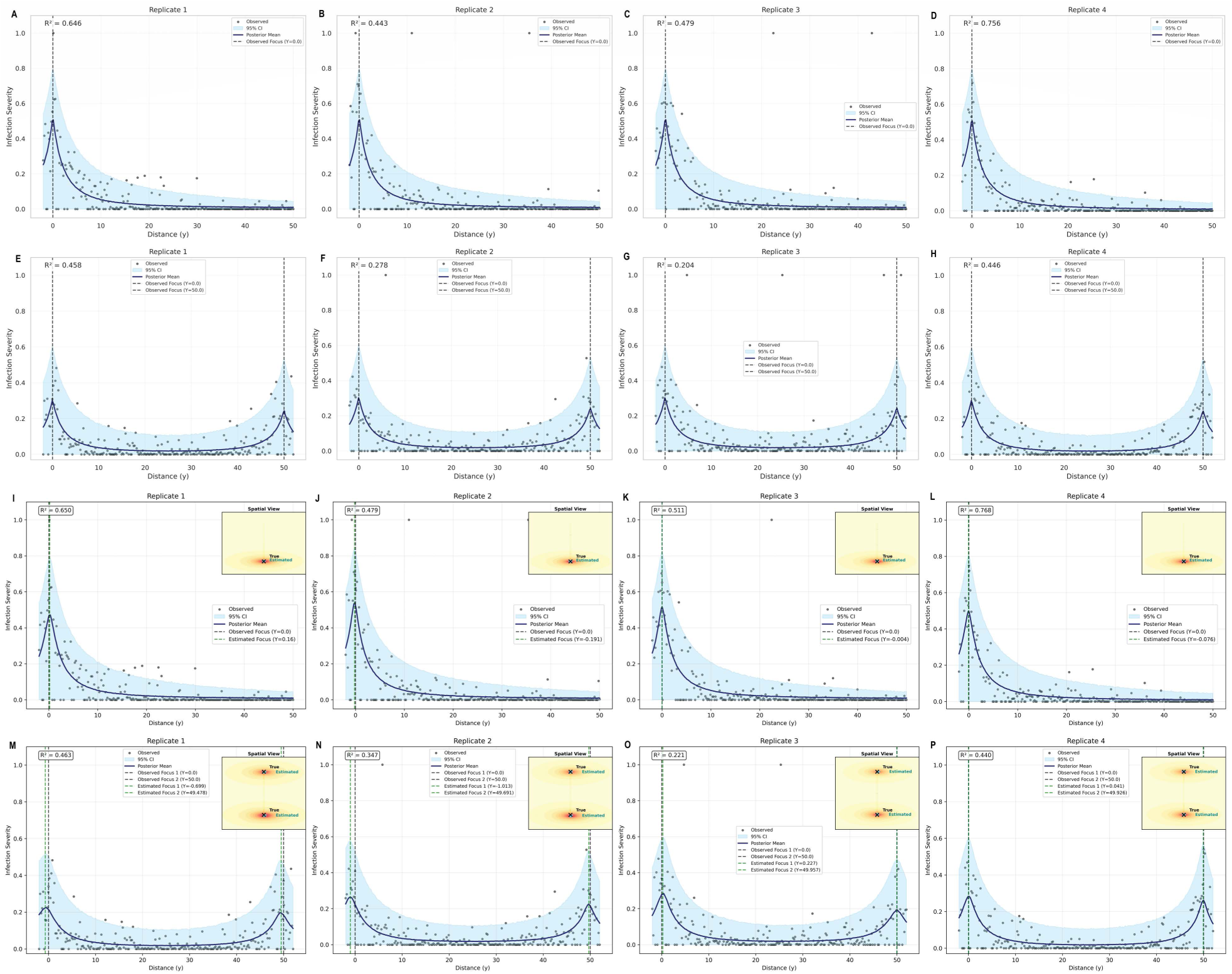
Spatial validation of posterior predictive distribution across hierarchical replicates under known and unknown focus location scenarios. Transect plots (x=0) illustrate the model’s fit for known single-focus (A-D), known two-foci (E-H), unknown single-focus (I-L), and unknown two-foci (M-P) scenarios. Observed infection severity (black points) is compared against the posterior predictive mean (navy line) and the 95% credible interval (shaded sky-blue region). Vertical dashed lines indicate the true foci positions at *y* = 0 and *y* = 50. The 2D insets show joint estimation of unknown infection sources locations (stars) relative to true foci (crosses) under weakly informative priors.

**Figure 4:**
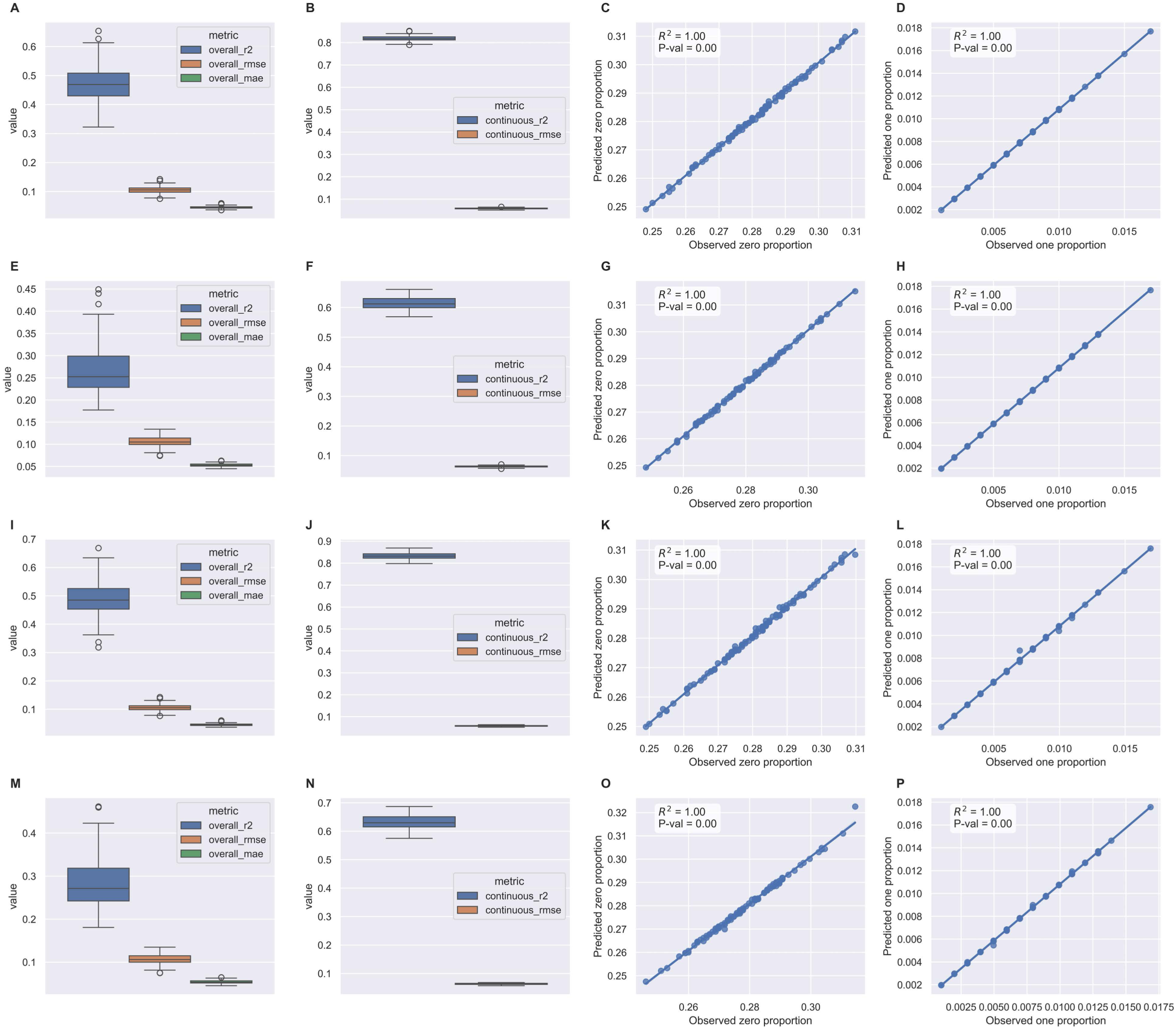
Statistical validation via posterior predictive checks (PPC) for single and two-foci dispersal across known and unknown focus location scenarios. First row (A-D) corresponds to known single-focus simulations; second row (E-H) corresponds to known two-foci scenario; third row (I-L) corresponds to unknown single-focus simulations; fourth row (M-P) corresponds to unknown two-foci scenario. **(A, E, I, M)** Distribution of aggregate goodness-of-fit metrics (*R*^2^, RMSE and MAE) across 100 simulations, indicating high predictive consistency. **(B, F, J, N)** Performance metrics for the continuous (Beta) component of the ZOIB model, assessing the recovery of non-extreme infection severity. **(C, G, K, O)** Correspondence between observed and predicted proportions of zero-severity. **(D, H, L, P)** Correspondence between observed and predicted proportions of one-severity. In panels C, D, G, H, K, L, O, and P the 1:1 identity line represents perfect model calibration.

**Table 2:** Parameter Recovery Performance for Known Single and Two-Foci Dispersal Scenarios. Values represent the mean across 100 simulations ± standard deviation.

| Scenario | Parameter | True | Estimate | Bias | Coverage | $\hat{R}$ | ESS (Bulk) |
| --- | --- | --- | --- | --- | --- | --- | --- |
| *Single-Focus* | **Dispersal Kernel** |  |  |  |  |  |  |
| | $a_1$ (Scale) | 5.00 | $5.02 \pm 0.54$ | +0.02 | 94.00% | 1.00 | 13,388 |
| | $b_1$ (Exponent) | 2.00 | $2.00 \pm 0.11$ | 0.00 | 95.00% | 1.00 | 12,583 |
|  | **ZOIB** |  |  |  |  |  |  |
| | $\pi_0$ (Zero-inflation) | 0.30 | $0.30 \pm 0.01$ | 0.00 | 95.00% | 1.00 | 27,567 |
| | $q$ (One-inflation) | 0.01 | $0.01 \pm 0.00$ | 0.00 | 93.00% | 1.00 | 25,638 |
| | $\phi$ (Precision) | 18.00 | $18.12 \pm 1.31$ | +0.12 | 96.00% | 1.00 | 22,444 |
| *Two-Foci* | **Dispersal Kernel & Mixture** |  |  |  |  |  |  |
| | $a_1$ (Scale) | 5.00 | $5.30 \pm 1.10$ | +0.30 | 100.00% | 1.00 | 10,023 |
| | $b_1$ (Exponent) | 2.00 | $2.06 \pm 0.29$ | +0.06 | 100.0% | 1.00 | 10,003 |
| | $a_2$ (Scale) | 5.00 | $5.33 \pm 1.09$ | +0.33 | 98.00% | 1.00 | 9,704 |
| | $b_2$ (Exponent) | 2.00 | $2.06 \pm 0.25$ | +0.06 | 98.00% | 1.00 | 9,756 |
| | $w$ (Weight) | 0.60 | $0.60 \pm 0.02$ | 0.00 | 96.00% | 1.00 | 14,937 |
|  | **ZOIB** |  |  |  |  |  |  |
| | $\pi_0$ (Zero-inflation) | 0.30 | $0.30 \pm 0.01$ | 0.00 | 96.00% | 1.00 | 26,672 |
| | $q$ (One-inflation) | 0.01 | $0.01 \pm 0.00$ | 0.00 | 95.00% | 1.00 | 26,185 |
| | $\phi$ (Precision) | 18.00 | $17.81 \pm 1.06$ | -0.19 | 93.00% | 1.00 | 28,060 |

Under unknown source locations, LOO-CV identified the Power Law kernel in 100% of simulations for both single-focus and two-foci scenarios (Figure 2 C–D), with ΔELPD_loo_ consistently exceeding the 2 × *SE* equivalence threshold.

The posterior predictive distribution highlights the performance of HiBASIL in jointly estimating foci locations, source intensities, and the dispersal kernels using weakly informative priors (Figure 3I–P), with predictive peaks aligning closely with true foci in both scenarios. Overall *R*^2^ was 0.49 ± 0.06 and 0.29 ± 0.06 for single-focus and two-foci scenarios, respectively. The predictive power was even more pronounced for the continuous component, with *R*^2^ values of 0.83 ± 0.01 and 0.63 ± 0.02, respectively. Zero- and one-inflation proportions remained well-calibrated under unknown source locations, consistent with the known-foci results above.

Parameter recovery remained robust under joint source localization (Table 3), with dispersal parameters absolute biases ranging from 0.01 to 0.94, source location absolute biases varying from 0.02 to 0.22, and mixture weight the bias of −0.08.

**Table 3:** Parameter recovery performance for Power Law dispersal kernel under joint source localization. Results are summarized across 100 simulations.

| Scenario | Parameter Group | True | Estimate | Bias | Coverage | $\hat{R}$ | ESS |
| --- | --- | --- | --- | --- | --- | --- | --- |
| *Single-Focus* | **Dispersal & Location** |  |  |  |  |  |  |
| | $a_1$ (Scale) | 5.00 | $4.06 \pm 0.60$ | -0.94 | 76% | 1.00 | 7,310 |
| | $b_1$ (Exponent) | 2.00 | $1.90 \pm 0.11$ | -0.10 | 85% | 1.00 | 9,466 |
| | $f_{x1}$ (Source X) | 0.00 | $0.11 \pm 0.05$ | +0.11 | 100% | 1.00 | 5,425 |
| | $f_{y1}$ (Source Y) | 0.00 | $-0.02 \pm 0.09$ | -0.02 | 98% | 1.00 | 21,272 |
|  | **ZOIB** |  |  |  |  |  |  |
| | $\pi_0$ (Zero-inflation) | 0.30 | $0.30 \pm 0.01$ | 0.00 | 96% | 1.00 | 23,690 |
| | $q$ (One-inflation) | 0.01 | $0.01 \pm 0.00$ | 0.00 | 93% | 1.00 | 24,566 |
| | $\varphi$ (Precision) | 18.00 | $18.13 \pm 1.32$ | +0.13 | 95% | 1.00 | 21,956 |
| *Two-Foci* | **Dispersal & Location & Mixture** |  |  |  |  |  |  |
| | $a_1$ (Scale 1) | 5.00 | $4.56 \pm 1.18$ | -0.44 | 95% | 1.01 | 15,115 |
| | $b_1$ (Exponent 1) | 2.00 | $2.01 \pm 0.30$ | +0.01 | 98% | 1.01 | 14,997 |
| | $f_{x1}$ (Source 1 X) | 0.00 | $0.18 \pm 0.09$ | +0.18 | 100% | 1.01 | 13,614 |
| | $f_{y1}$ (Source 1 Y) | 0.00 | $-0.08 \pm 0.17$ | -0.08 | 97% | 1.01 | 28,490 |
| | $a_2$ (Scale 2) | 5.00 | $4.71 \pm 1.02$ | -0.29 | 98% | 1.01 | 14,538 |
| | $b_2$ (Exponent 2) | 2.00 | $2.06 \pm 0.26$ | +0.06 | 99% | 1.01 | 14,655 |
| | $f_{x2}$ (Source 2 X) | 0.00 | $0.22 \pm 0.10$ | +0.22 | 100% | 1.01 | 12,832 |
| | $f_{y2}$ (Source 2 Y) | 50.00 | $50.11 \pm 0.24$ | +0.11 | 98% | 1.01 | 25,715 |
| | $w$ (Weight) | 0.60 | $0.52 \pm 0.02$ | -0.08 | 98% | 1.01 | 16,867 |
|  | **ZOIB** |  |  |  |  |  |  |
| | $\pi_0$ (Zero-inflation) | 0.30 | $0.30 \pm 0.01$ | 0.00 | 97% | 1.01 | 40,054 |
| | $q$ (One-inflation) | 0.01 | $0.01 \pm 0.00$ | 0.00 | 94% | 1.01 | 37,972 |
| | $\varphi$ (Precision) | 18.00 | $17.74 \pm 1.07$ | -0.26 | 95% | 1.00 | 32,662 |

#### Parameter Recovery Accuracy and Model Robustness under Multiple Epidemic Scenarios

The Power Law kernel was identified with 100% accuracy across Base, Narrow & Steep, Wide & Steep, and Wide & Shallow scenarios (Figure 5). In the Wide & Shallow scenario (Figure 6M-P), the diffuse infection signals compressed the ΔELPD separation between competing kernels and correspondingly narrowed the equivalence zone itself (Figure 5D); despite this tighter margin, kernel identification remained 100% accurate.

**Figure 5:**
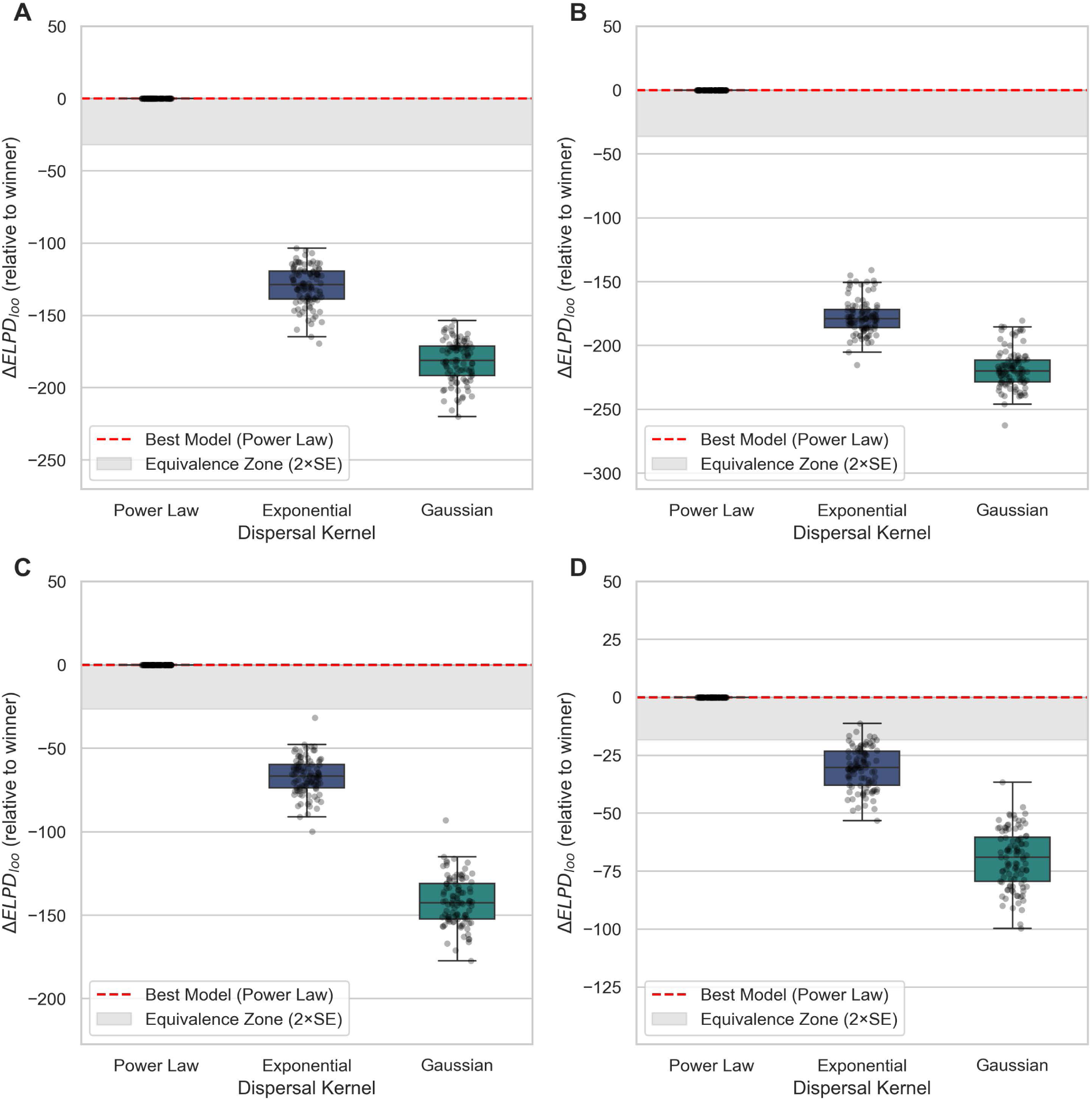
Model selection sensitivity and structural identifiability of dispersal kernels across four epidemiological scenarios. Panels (A) Base, (B) Narrow & Steep, (C) Wide & Steep, and (D) Wide & Shallow correspond to the scenarios defined in Table S1. The distribution of Δ*ELPD_loo_* relative to the top-ranked model demonstrates a 100% success rate in correctly identifying the Power Law kernel across all 400 simulations. In the Wide & Shallow scenario (D), diffuse infection signals compressed the ΔELPD separation between competing kernels, bringing the alternatives closer to the winner; nevertheless, these reduced gaps still exceeded the 2 × *SE* equivalence zone, so kernel identification remained perfect and decisive throughout.

**Figure 6:**
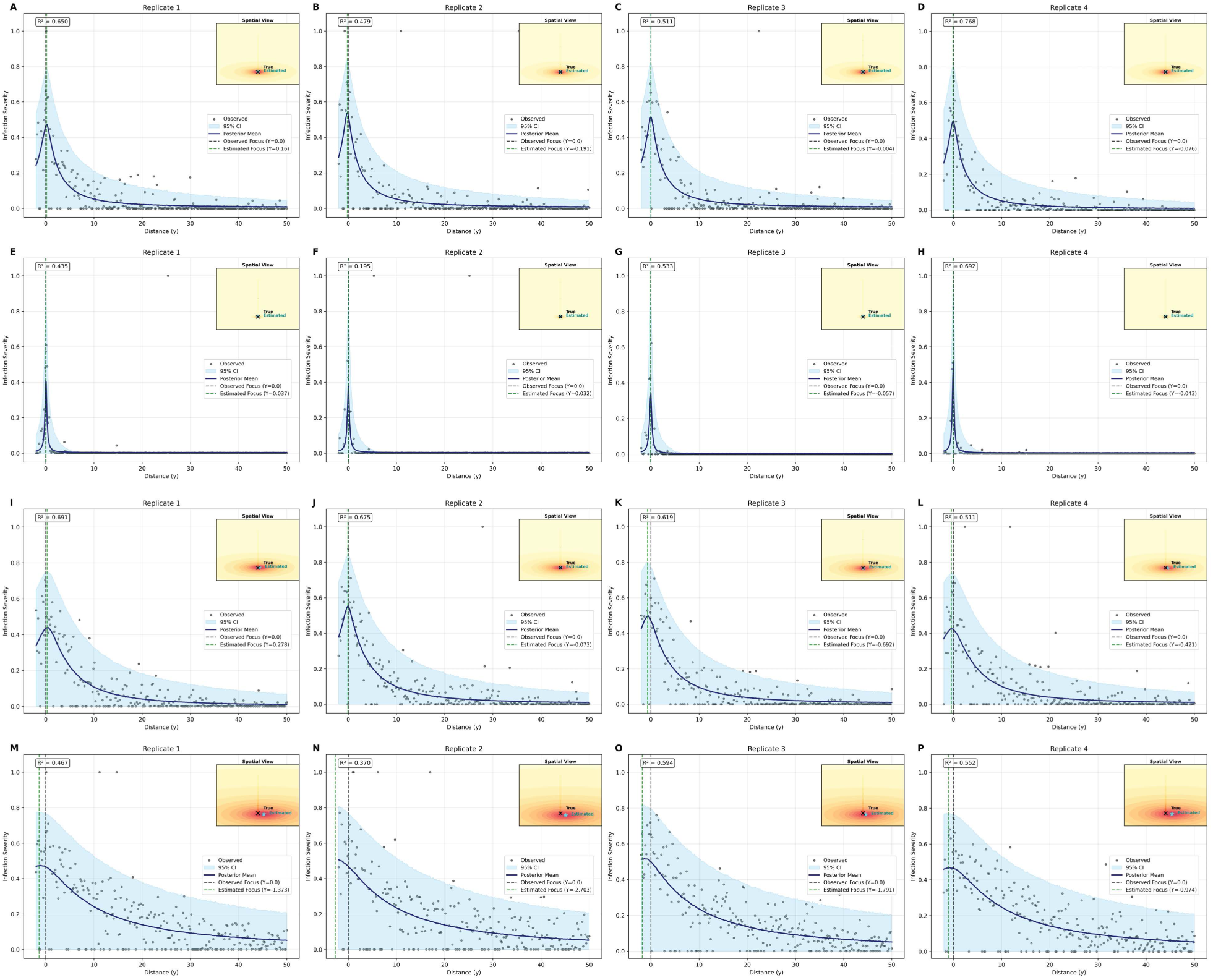
Spatial validation of posterior predictive distribution across four dispersal scenarios. Main panels present 1D spatial transect plots at *x* = 0, comparing observed severity (black points) against the posterior predictive mean (navy line) and 95% credible interval (shaded sky-blue region) for the **Base** (A-D), **Narrow & Steep** (E-H), **Wide & Steep (I-L)**, and **Wide & Shallow** (M-P) scenarios defined in Table S1. The vertical dashed lines indicate the estimated infection focus at *y* = 0. Corresponding 2D insets show joint estimation of unknown infection source locations (stars) relative to true foci (crosses) under weakly informative priors. The Narrow & Steep scenario (E-H) illustrates the highest spatial data sparsity, with the majority of observations clustered near the origin.

Posterior predictive checks (PPC) across 400 simulations (Figure 6) showed that Base, Wide & Steep, and Wide & Shallow scenarios achieved aggregated *R*^2^ of 0.44 to 0.53 (Figure S1 A, I, and M) with near-perfect zero-proportion prediction (*R*^2^ = 1.00) within a 0.2 to 0.3 range (Figure S1C, K, O). The Narrow & Steep scenario (Figure 6E-H) exhibited the highest spatial data sparsity with a significant higher zero proportion (0.60 to 0.66; Figure S1G), which source from the structural zero-inflation (*π*_0_ = 0.3) and kernel-driven decay to near-zero severity. The model therefore over-generated zeros and yielded both weak and biased calibration of the predicted zero proportion (*R*^2^ = 0.69, versus *R*^2^ = 1.0 in al other scenarios).

Parameter recovery was excellent for the Base and Wide scenarios across dispersal, location, and the ZOIB parameters (Table 4). In the Narrow & Steep scenario at the *y* = 50, most sampled points fell outside the active dispersal zone, yielding poor recovery of dispersal parameters, though focus coordinates and one-inflation parameters remained well-recovered across all tests. Under this scale mismatch, the two zero sources become non-identifiable, and *π*_0_ collapsed from its structural target onto the realized zero fraction of the data (Figure S1G). Model performance was contingent upon whether the observation scale matched the pathogen’s biological dispersal reach. Refining the observation window to match the Narrow & Steep pathogen’s dispersal radius (*y* = 8) resolved the estimation bias (Table S5; Figure S2). Within this optimized window, the zero-inflation parameter (*π*_0_) was recovered with 0.30 ± 0.01 and 96.0% coverage (Table S6; Figure S4). Furthermore, the scale (*a*) and exponent (*b*) recovery improved to 77.0% and 88.0% coverage, respectively (*Ȓ* = 1.00; Table S6). For the Wide & Shallow scenario, extending the range to *y* = 75 captured critical “long-tail” information that was truncated at smaller scales, significantly increasing ΔELPD between the Power Law and alternative kernels (Table S5; Figure S2; Figure S3).

**Table 4:**
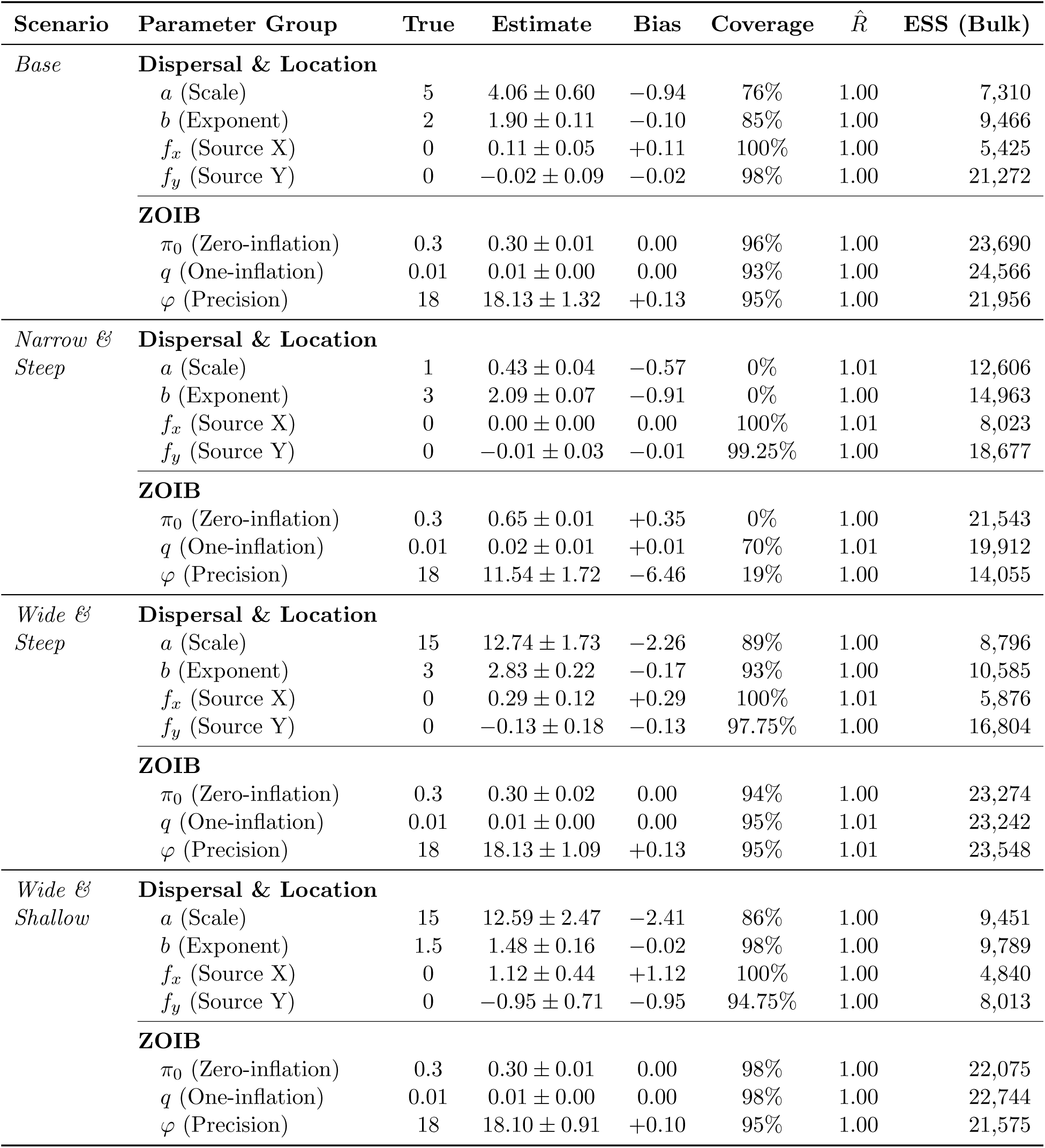
Parameter recovery performance for unknown single-focus dispersal across four epidemiological scenarios. Values represent the mean across 100 simulations ± standard deviation.

| Scenario | Parameter Group | True | Estimate | Bias | Coverage | $\hat{R}$ | ESS (Bulk) |
| --- | --- | --- | --- | --- | --- | --- | --- |
| *Base* | **Dispersal & Location** |  |  |  |  |  |  |
| | $a$ (Scale) | 5 | $4.06 \pm 0.60$ | $-0.94$ | 76% | 1.00 | 7,310 |
| | $b$ (Exponent) | 2 | $1.90 \pm 0.11$ | $-0.10$ | 85% | 1.00 | 9,466 |
| | $f_x$ (Source X) | 0 | $0.11 \pm 0.05$ | $+0.11$ | 100% | 1.00 | 5,425 |
| | $f_y$ (Source Y) | 0 | $-0.02 \pm 0.09$ | $-0.02$ | 98% | 1.00 | 21,272 |
|  | **ZOIB** |  |  |  |  |  |  |
| | $\pi_0$ (Zero-inflation) | 0.3 | $0.30 \pm 0.01$ | $0.00$ | 96% | 1.00 | 23,690 |
| | $q$ (One-inflation) | 0.01 | $0.01 \pm 0.00$ | $0.00$ | 93% | 1.00 | 24,566 |
| | $\varphi$ (Precision) | 18 | $18.13 \pm 1.32$ | $+0.13$ | 95% | 1.00 | 21,956 |
| *Narrow & Steep* | **Dispersal & Location** |  |  |  |  |  |  |
| | $a$ (Scale) | 1 | $0.43 \pm 0.04$ | $-0.57$ | 0% | 1.01 | 12,606 |
| | $b$ (Exponent) | 3 | $2.09 \pm 0.07$ | $-0.91$ | 0% | 1.00 | 14,963 |
| | $f_x$ (Source X) | 0 | $0.00 \pm 0.00$ | $0.00$ | 100% | 1.01 | 8,023 |
| | $f_y$ (Source Y) | 0 | $-0.01 \pm 0.03$ | $-0.01$ | 99.25% | 1.00 | 18,677 |
|  | **ZOIB** |  |  |  |  |  |  |
| | $\pi_0$ (Zero-inflation) | 0.3 | $0.65 \pm 0.01$ | $+0.35$ | 0% | 1.00 | 21,543 |
| | $q$ (One-inflation) | 0.01 | $0.02 \pm 0.01$ | $+0.01$ | 70% | 1.01 | 19,912 |
| | $\varphi$ (Precision) | 18 | $11.54 \pm 1.72$ | $-6.46$ | 19% | 1.00 | 14,055 |
| *Wide & Steep* | **Dispersal & Location** |  |  |  |  |  |  |
| | $a$ (Scale) | 15 | $12.74 \pm 1.73$ | $-2.26$ | 89% | 1.00 | 8,796 |
| | $b$ (Exponent) | 3 | $2.83 \pm 0.22$ | $-0.17$ | 93% | 1.00 | 10,585 |
| | $f_x$ (Source X) | 0 | $0.29 \pm 0.12$ | $+0.29$ | 100% | 1.01 | 5,876 |
| | $f_y$ (Source Y) | 0 | $-0.13 \pm 0.18$ | $-0.13$ | 97.75% | 1.00 | 16,804 |
|  | **ZOIB** |  |  |  |  |  |  |
| | $\pi_0$ (Zero-inflation) | 0.3 | $0.30 \pm 0.02$ | $0.00$ | 94% | 1.00 | 23,274 |
| | $q$ (One-inflation) | 0.01 | $0.01 \pm 0.00$ | $0.00$ | 95% | 1.01 | 23,242 |
| | $\varphi$ (Precision) | 18 | $18.13 \pm 1.09$ | $+0.13$ | 95% | 1.01 | 23,548 |
| *Wide & Shallow* | **Dispersal & Location** |  |  |  |  |  |  |
| | $a$ (Scale) | 15 | $12.59 \pm 2.47$ | $-2.41$ | 86% | 1.00 | 9,451 |
| | $b$ (Exponent) | 1.5 | $1.48 \pm 0.16$ | $-0.02$ | 98% | 1.00 | 9,789 |
| | $f_x$ (Source X) | 0 | $1.12 \pm 0.44$ | $+1.12$ | 100% | 1.00 | 4,840 |
| | $f_y$ (Source Y) | 0 | $-0.95 \pm 0.71$ | $-0.95$ | 94.75% | 1.00 | 8,013 |
|  | **ZOIB** |  |  |  |  |  |  |
| | $\pi_0$ (Zero-inflation) | 0.3 | $0.30 \pm 0.01$ | $0.00$ | 98% | 1.00 | 22,075 |
| | $q$ (One-inflation) | 0.01 | $0.01 \pm 0.00$ | $0.00$ | 98% | 1.00 | 22,744 |
| | $\varphi$ (Precision) | 18 | $18.10 \pm 0.91$ | $+0.10$ | 95% | 1.00 | 21,575 |

#### Inference Robustness under Varied Foci Mixture Weights

HiBASIL identified the Power Law kernel with a 100% accuracy across all mixture configurations: balanced (0.5, 0.5), nearly balanced (0.6, 0.4), skewed (0.8, 0.2), and extremely skewed (0.95, 0.05)(Figure 7). At the most extreme configuration (0.95, 0.05), the ΔELPD distribution exhibits partial overlap with the aggregated 2 × *SE* equivalence zone (Figure 7D), suggesting the secondary focus approaches the lower limit of predictive detectability at a 5% contribution. The Beta(2, 2) weight prior additionally exerted a shrinkage effect toward the prior mean at this extreme.

**Figure 7:**
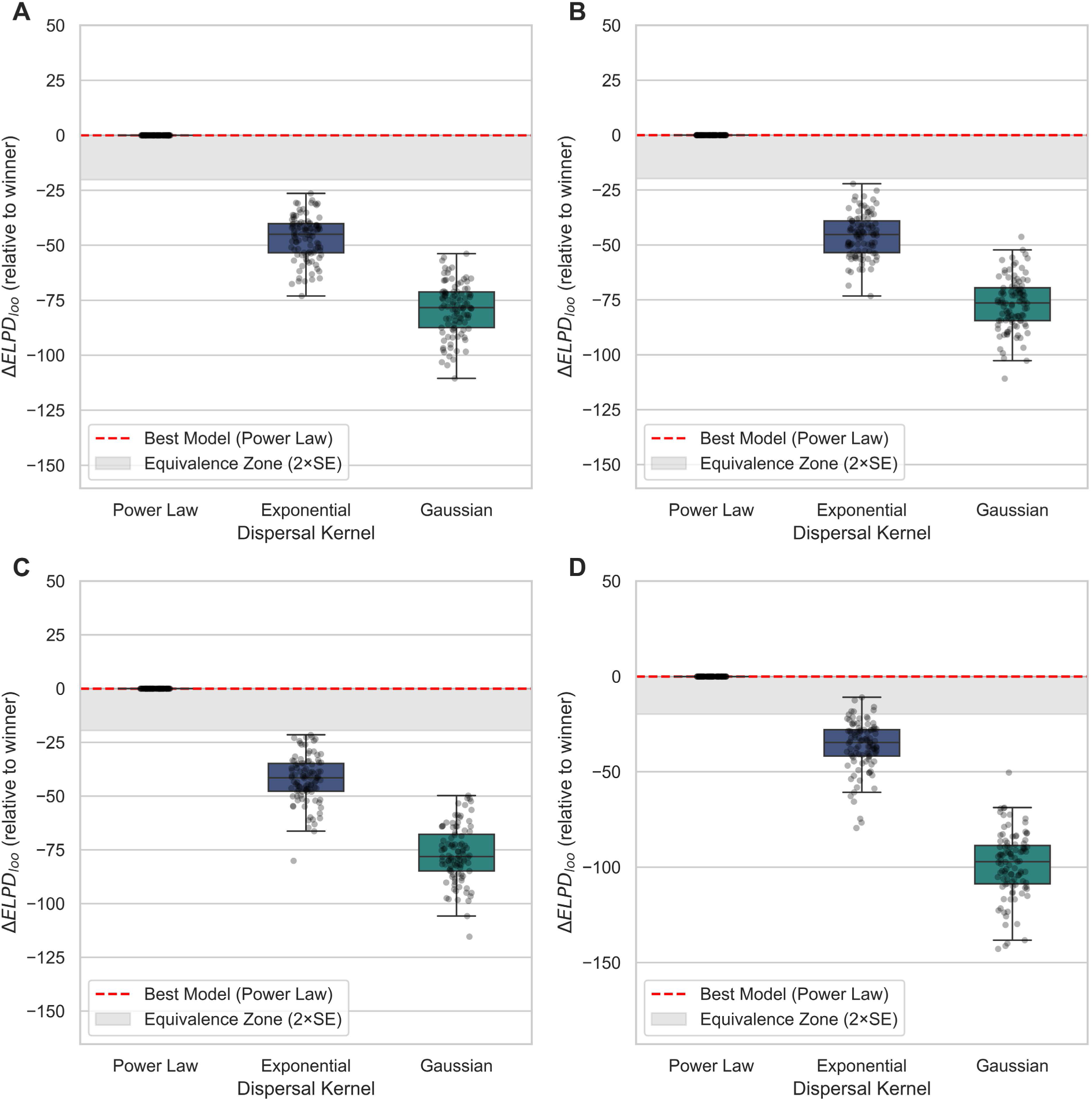
Model selection sensitivity and structural identifiability of dispersal kernels across varying foci mixture weights. Panels display the distribution of Δ*ELPD_loo_* relative to the top-ranked model for mixture configurations of (A) (0.5, 0.5), (B) (0.6, 0.4), (C) (0.8, 0.2), and (D) (0.95, 0.05). Across all 400 simulations, the Bayesian framework correctly identified the generative Power Law kernel with a 100% success rate. In the marginal configuration (D; 0.95/0.05), the Δ*ELPD* distribution exhibits partial overlap with the aggregated 2 × *SE* equivalence zone (gray shaded region). This overlap indicates that at a 5% contribution threshold, the secondary focus’s spatial signal approaches the limit of predictive detectability.

Posterior predictive checks (PPC) indicated a high predictive performance across all mixture scenarios (Figure 8). Aggregated results across 100 simulations per scenario yielded global *R*^2^ values between 0.29±0.06 to 0.46±0.07 (Figure S5 A, E, I, M), while the component-specific *R*^2^ for the continuous Beta component varied from 0.63±0.02 to 0.81±0.02 (Figure S5 B, F, J, N). Zero-(Figure S5 C, G, K, O) and one-inflation (Figure S5 D, H, L, P) proportions were near-perfectly calibrated (*R*^2^ = 1.00) across all mixture configurations.

**Figure 8:**
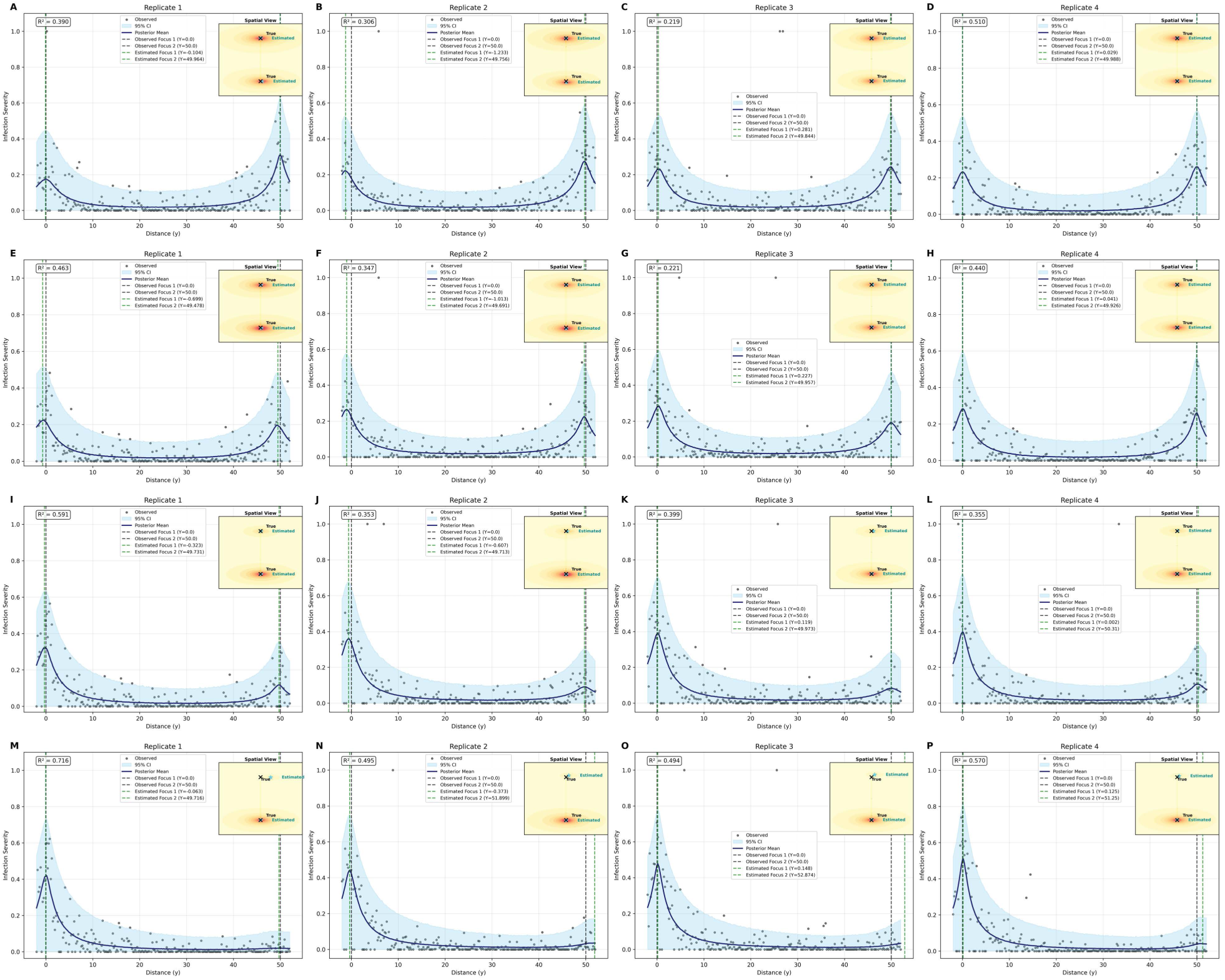
Spatial validation of posterior predictive distribution across diverse foci mixture weights. The main panels present 1D spatial transects at *x* = 0, comparing observed severity (black points) against the posterior predictive mean (navy line) and 95% credible interval (shaded sky-blue region). Configurations range from balanced mixtures (A-D) (0.5, 0.5) and (E-H) (0.6, 0.4) to highly skewed and marginal scenarios (I-L) (0.8, 0.2) and (M-P) (0.95, 0.05). The vertical dashed lines denote the estimated source locations at *y* = 0 and *y* = 50. The 2D insets show joint estimation of unknown infection source locations (stars) relative to true foci (crosses) under weakly informative priors.

The recovery of parameters remained robust across mixture configurations for dispersal, foci locations, and ZOIB parameters (Table 5), but mixture weights bias increased with skewness. The primary foci intensity parameter, *f_z_*_1_, consistently exhibited a high bias (approximately 0.24 to 0.28) with negligible coverage of the true value (0% - 18.50%).

**Table 5:**
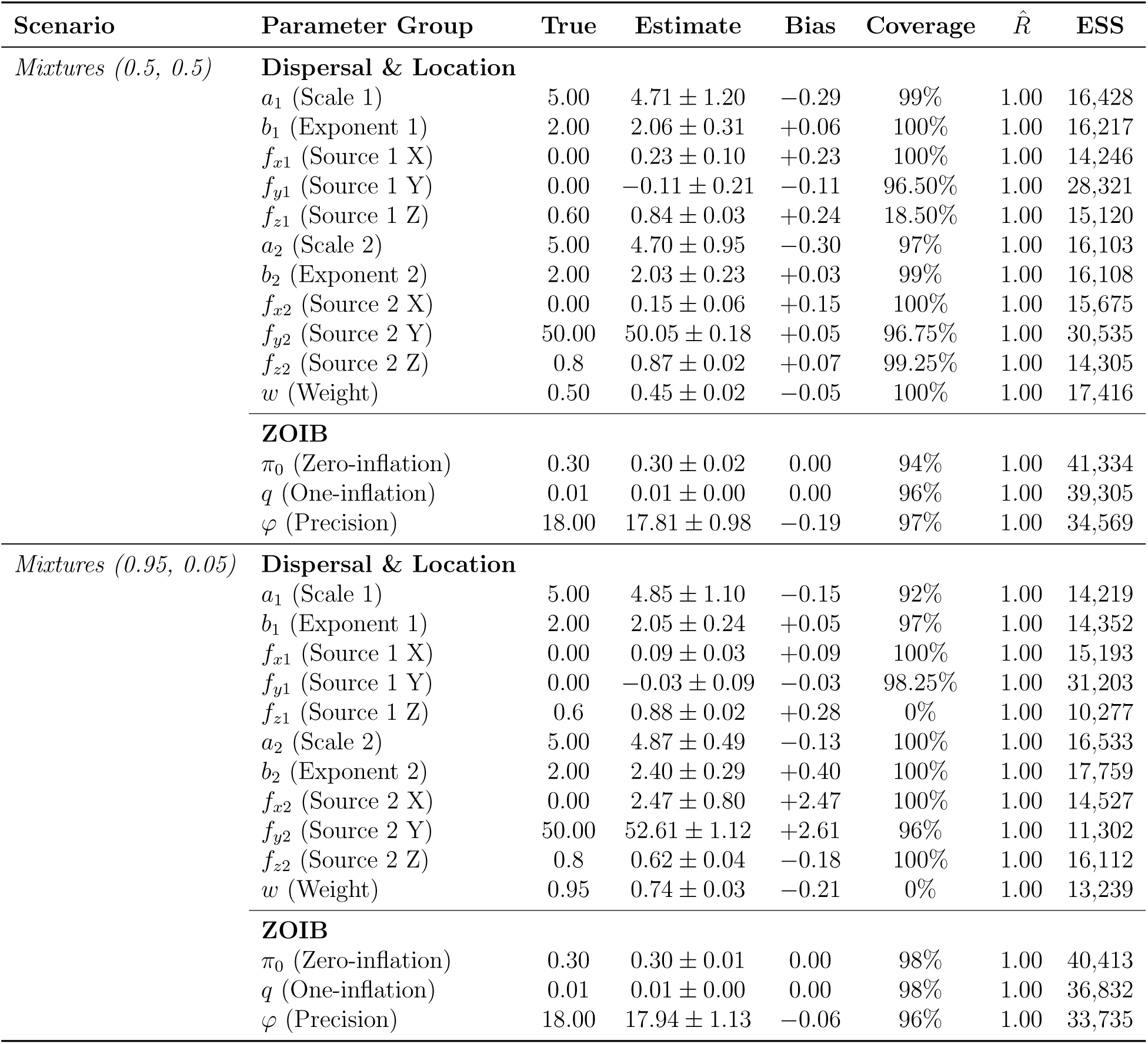
Parameter recovery performance for Power Law dispersal kernel across all foci mixture scenarios. Results are summarized across 100 simulations. Here we only show the balanced and extremely skewed conditions and for full details, please see Table S7.

Pearson correlation analysis of parameter estimation biases revealed the underlying structural confounding responsible for these shifts (Figure S6). The correlation between primary scale (*a*1) and exponent (*b*1) remained stable (*r >* 0.9), indicating persistent structural confounding inherent to the kernel geometry. In contrast, uncertainty in the secondary location (*f_x_*_2_ and *f_y_*_2_) and confounding between primary intensity (*f_z_*_1_) and weight increased substantially with mixture imbalance. Furthermore, the primary and secondary foci intensities (*f_z_*_1_ and *f_z_*_2_) exhibited persistent high correlation. These patterns suggest an identifiability trade-off wherein precise partitioning of source strength between disease intensity and the mixture weight may require more informative priors or temporal data to fully decouple.

#### Structural Robustness and Kernel Identifiability

We assessed model performance under under-parameterization (fitting a Single-focus model to Two-foci data) and over-parameterization (fitting a Two-foci model to Single-focus data).

##### Kernel Identifiability

The Power Law kernel was identified as the top-ranked model in 100% simulations across both misspecification scenarios (Figure 9). Specifically, in the under-parametrization scenario, while the Power Law kernel was universally preferred, Exponential and Gaussian kernels were statistically indistinguishable from the top-ranked model (within 2 × *SE* of Δ*ELPD*) only in 4% and 8% of simulations, respectively (Figure 9A). In the over-parameterization scenario, the Power Law remained the best-performing model in 98% of cases and remained predictive equivalent to the winner for all 100 simulations. While the Exponential model was statistically indistinguishable from the winner in 24% of simulations, the Δ*ELPD* distributions remained largely distinct (Figure 9B).

**Figure 9:**
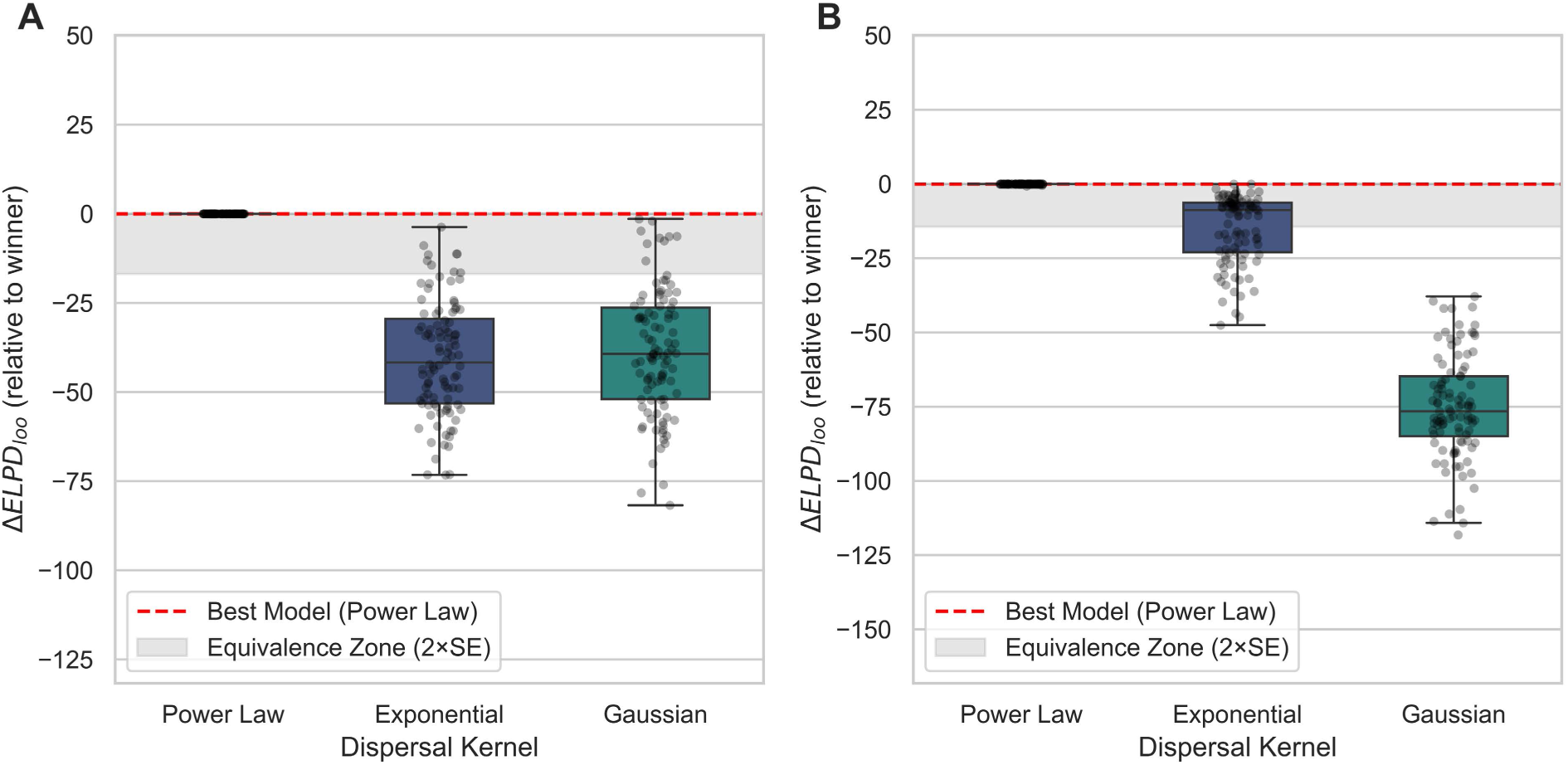
Model selection sensitivity and structural identifiability of dispersal kernels under model misspecification. Distributions of Δ*ELPD_loo_* are shown relative to the top-ranked model for scenarios involving (A) a Single-focus model applied to Two-foci data, and (B) a Two-foci model applied to Single-focus data. The Power Law dispersal kernel was correctly identified in 100% of 200 simulations. In the scenario A, the Power Law kernel was identified as the best-performing model in all 100 times, though the Exponential and Gaussian kernels were predictive equivalent to the top-ranked model for 4 and 8 times, respectively. So both the Δ*ELPD* distribution of Exponential and Gaussian exhibits partial overlap with the aggregated 2 × *SE* equivalence zone (gray shaded region). In scenario B, the Power Law was the top-ranked model for 98 times and remained predictive equivalent to the winner for all 100 times. While the Exponential was predictive equivalent to the top-ranked model for 24 times, its Δ*ELPD* distribution remains largely distinct from the equivalence zone.

##### Predictive Stability and Posterior Diagnostics

Posterior predictive checks (PPC) revealed distinct behaviors for each scenario (Figure 10; Figure S7). For the under-parameterized scenario, the 95% CI captured sample variation but the posterior predictive mean was less precise than the correctly specified Two-foci model, with significantly lower global and continuous Beta *R*^2^ values (Figure 10A&B; Figure S7A&B; Figure 4I&J). For the over-parameterized scenario, posterior predictions were nearly identical to the correctly specified Single-focus model with no significant degradation (Figure 10E&F; Figure S7E&F; Figure 4M&N). Zero- and one-inflation proportions remained in near-perfect alignment across both scenarios (*R*^2^ = 1.00; Figure S7C, D, G, H).

**Figure 10:**
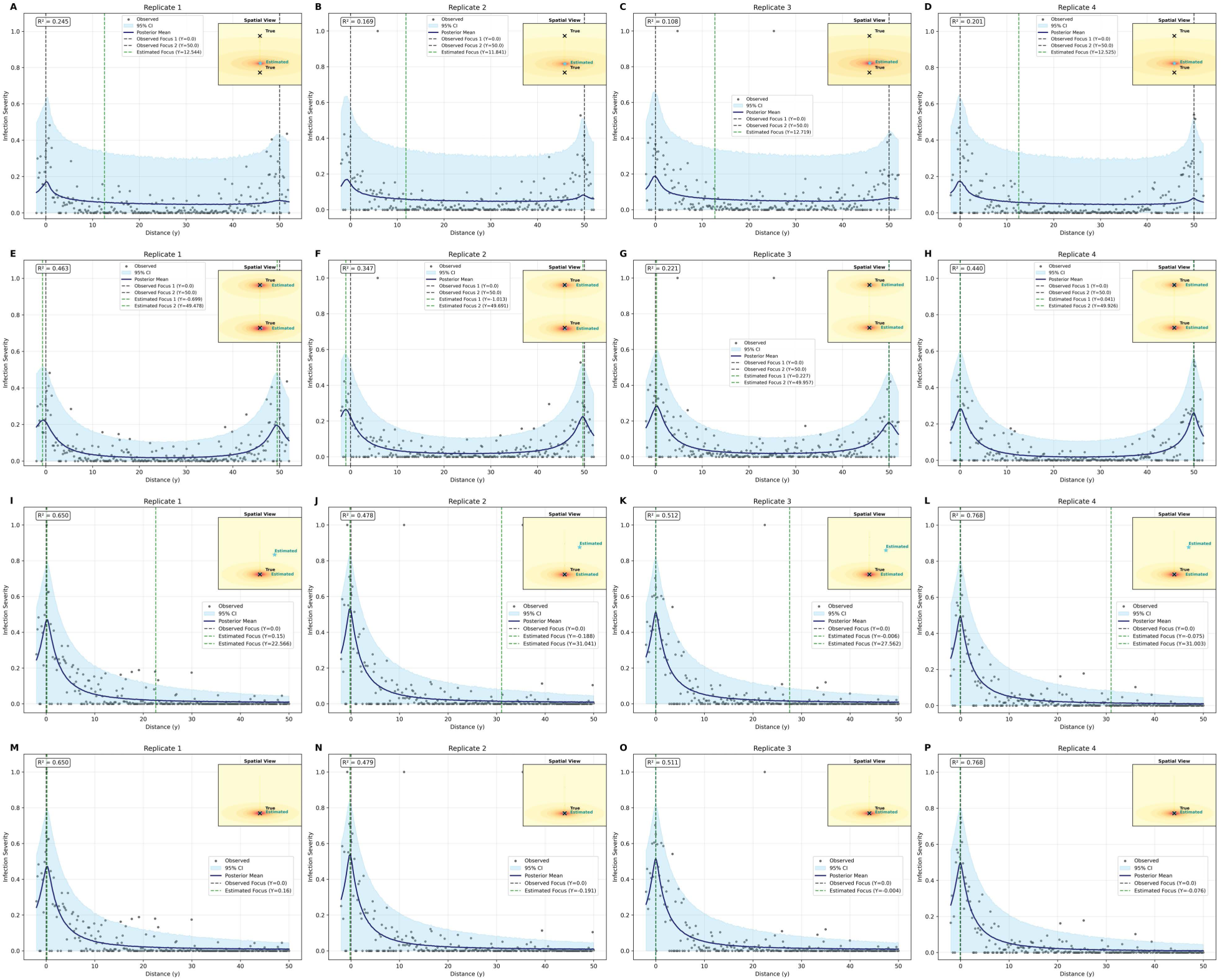
Spatial validation of posterior predictive distribution under correctly specified and misspecified model scenarios. Main panels show 1D spatial transects at *x* = 0, comparing observed severity (black points) against the posterior predictive mean (navy line) and 95% credible interval (shaded sky-blue region). (A-D) Single-focus model applied to Two-foci data, (E-H) Two-foci model applied to Two-foci data, (I-L) Two-foci model applied to Single-focus data, and (M-P) Single-focus model applied to Single-focus data. Vertical dashed lines indicate estimated source locations at *y* = 0 and *y* = 50. 2D insets show joint estimation of unknown infection source locations (stars) relative to the true foci (crosses) under weakly informative priors. Under the misspecification of a Single-focus model applied to Two-foci data (A-D), the posterior distribution exhibits characteristic bimodal peaks.

Misspecification was signaled through distinct parameter behaviors. Under under-parameterization, dispersal scale (*a*1), exponent (*b*1), focus *y* coordinate, focus intensity *f_z_*, and ZOIB precision *φ* exhibited large error or poor recovery with *Ȓ* > 1.01 (Table 6), with this instability manifesting as bimodality in the posterior of source location *f_y_* (Figure 10A-D). Peak detection on the posterior of *f_y_* identified bimodal distributions in 53.25% of replicates (213 out of 400 rows; each simulation has four rows as replicates), of which 96.7% were well-separated (mean distance =50.38), with peaks accurately localizing near the ground true values at *y* = −0.17 and *y* = 50.20 (Supplementary File dataset S2). This indicates that the single-focus posterior recovers the hidden two-source spatial structure.

**Table 6:**
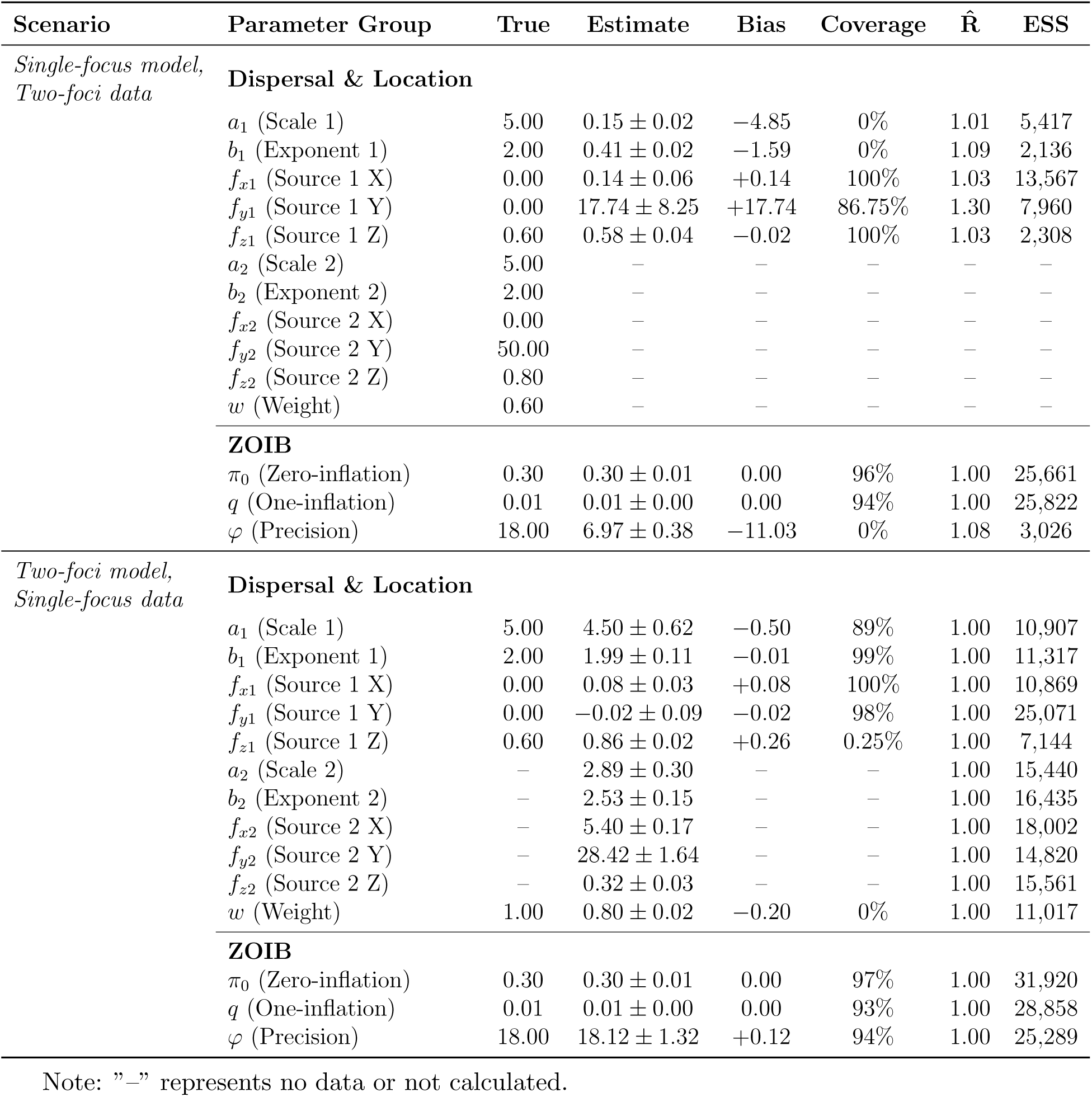
Performance of parameters estimation under model misspecification. Values represent the mean across 100 simulations ± standard deviation.

| Scenario | Parameter Group | True | Estimate | Bias | Coverage | $\hat{R}$ | ESS |
| --- | --- | --- | --- | --- | --- | --- | --- |
| *Single-focus model,*<br>*Two-foci data* | **Dispersal & Location** |  |  |  |  |  |  |
| | $a_1$ (Scale 1) | 5.00 | $0.15 \pm 0.02$ | -4.85 | 0% | 1.01 | 5,417 |
| | $b_1$ (Exponent 1) | 2.00 | $0.41 \pm 0.02$ | -1.59 | 0% | 1.09 | 2,136 |
| | $f_{x1}$ (Source 1 X) | 0.00 | $0.14 \pm 0.06$ | +0.14 | 100% | 1.03 | 13,567 |
| | $f_{y1}$ (Source 1 Y) | 0.00 | $17.74 \pm 8.25$ | +17.74 | 86.75% | 1.30 | 7,960 |
| | $f_{z1}$ (Source 1 Z) | 0.60 | $0.58 \pm 0.04$ | -0.02 | 100% | 1.03 | 2,308 |
| | $a_2$ (Scale 2) | 5.00 | — | — | — | — | — |
| | $b_2$ (Exponent 2) | 2.00 | — | — | — | — | — |
| | $f_{x2}$ (Source 2 X) | 0.00 | — | — | — | — | — |
| | $f_{y2}$ (Source 2 Y) | 50.00 | — | — | — | — | — |
| | $f_{z2}$ (Source 2 Z) | 0.80 | — | — | — | — | — |
| | $w$ (Weight) | 0.60 | — | — | — | — | — |
|  | **ZOIB** |  |  |  |  |  |  |
| | $\pi_0$ (Zero-inflation) | 0.30 | $0.30 \pm 0.01$ | 0.00 | 96% | 1.00 | 25,661 |
| | $q$ (One-inflation) | 0.01 | $0.01 \pm 0.00$ | 0.00 | 94% | 1.00 | 25,822 |
| | $\varphi$ (Precision) | 18.00 | $6.97 \pm 0.38$ | -11.03 | 0% | 1.08 | 3,026 |
| *Two-foci model,*<br>*Single-focus data* | **Dispersal & Location** |  |  |  |  |  |  |
| | $a_1$ (Scale 1) | 5.00 | $4.50 \pm 0.62$ | -0.50 | 89% | 1.00 | 10,907 |
| | $b_1$ (Exponent 1) | 2.00 | $1.99 \pm 0.11$ | -0.01 | 99% | 1.00 | 11,317 |
| | $f_{x1}$ (Source 1 X) | 0.00 | $0.08 \pm 0.03$ | +0.08 | 100% | 1.00 | 10,869 |
| | $f_{y1}$ (Source 1 Y) | 0.00 | $-0.02 \pm 0.09$ | -0.02 | 98% | 1.00 | 25,071 |
| | $f_{z1}$ (Source 1 Z) | 0.60 | $0.86 \pm 0.02$ | +0.26 | 0.25% | 1.00 | 7,144 |
| | $a_2$ (Scale 2) | — | $2.89 \pm 0.30$ | — | — | 1.00 | 15,440 |
| | $b_2$ (Exponent 2) | — | $2.53 \pm 0.15$ | — | — | 1.00 | 16,435 |
| | $f_{x2}$ (Source 2 X) | — | $5.40 \pm 0.17$ | — | — | 1.00 | 18,002 |
| | $f_{y2}$ (Source 2 Y) | — | $28.42 \pm 1.64$ | — | — | 1.00 | 14,820 |
| | $f_{z2}$ (Source 2 Z) | — | $0.32 \pm 0.03$ | — | — | 1.00 | 15,561 |
| | $w$ (Weight) | 1.00 | $0.80 \pm 0.02$ | -0.20 | 0% | 1.00 | 11,017 |
|  | **ZOIB** |  |  |  |  |  |  |
| | $\pi_0$ (Zero-inflation) | 0.30 | $0.30 \pm 0.01$ | 0.00 | 97% | 1.00 | 31,920 |
| | $q$ (One-inflation) | 0.01 | $0.01 \pm 0.00$ | 0.00 | 93% | 1.00 | 28,858 |
| | $\varphi$ (Precision) | 18.00 | $18.12 \pm 1.32$ | +0.12 | 94% | 1.00 | 25,289 |

Note: "—" represents no data or not calculated.

For the over-parameterized scenario, 96% of replicates (384/400) showed unimodal posteriors with mean peak at *y* = −0.002 (true *y* = 0); the remaining 4% (16/400) replicates were bimodal but poorly separated (mean distance was 0.13) (Supplementary File dataset S3). While parameters for the primary focus were estimated accurately, the framework effectively nullified the redundant second focus by assigning it a negligible mixture weight (1 − *w* = 0.2) and low intensity (*f_z_*_2_ = 0.32 ± 0.03) (Table 6). Consequently, disease severity contributed by the superfluous source was negligible (≈ 0.06 at the source).

#### Impact of Sample Size on Inferential Model Performance

The Power Law kernel was identified as the best model in 100% of 800 simulations across all sample sizes and spatial configurations (Figure 11). Identifiability was subtly constrained under low data density (*N* = 50): in the single-focus scenario, the Exponential kernel was predictively equivalent to the Power Law for one time (Figure 11A); in the two-foci scenario, Exponential and Gaussian kernels were predictively equivalent for 19 and 8 times, respectively (Figure 11E). At *N* = 100 under two-foci scenario, the Exponential kernel achieved predictively equivalent for 4 times (Figure 11F). In all remaining scenarios, the ΔELPD distributions were well-separated across all configurations (Figure 11B, C, D, G, H).

**Figure 11:**
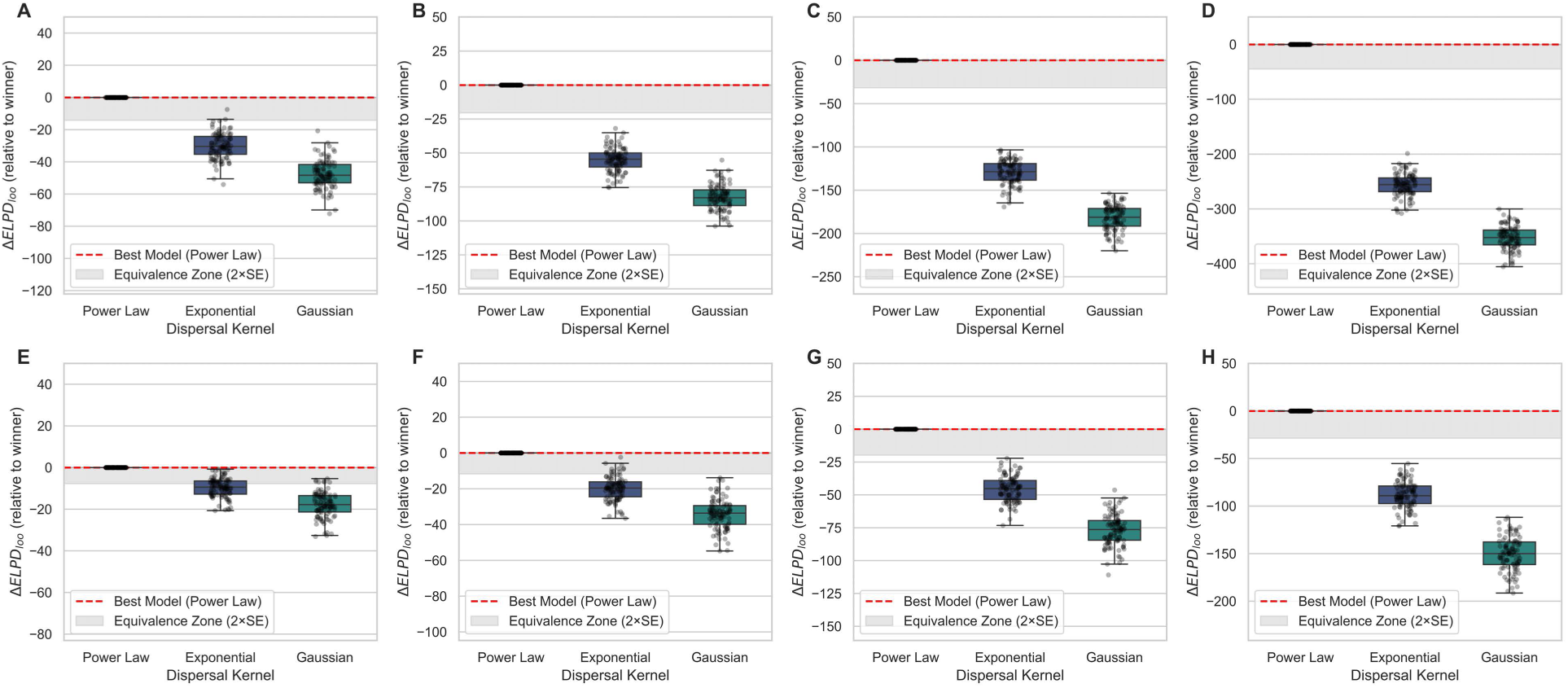
Model selection sensitivity and structural identifiability of dispersal kernels across gradients of sample sizes and spatial complexity. Distributions of the expected log pointwise predictive density (Δ*ELPD_loo_*) are presented relative to the best-performing model for eight distinct simulation scenarios. Panels (A-D) represent single-focus scenarios with samples sizes of *N* = 50, 100, 250, and 500, respectively; Panels (E-H) represent the corresponding two-foci scenarios. The Power Law kernel was correctly identified in 100% of 800 simulations. In the *N* = 50 single-focus scenario (A), the Power Law model was identified as the best-performing model in all simulations, although the Exponential kernel was found to be predictively equivalent to the Power law model in one case. Consequently, the Δ*ELPD* distribution for the Exponential kernel exhibited a marginal overlap with the aggregated 2 × *SE* equivalence zone (gray shaded region). Similarly, in the *N* = 50 two-foci scenario (E), the Power Law remained the best model 100% of the time, the Exponential and Gaussian kernels were predictively equivalent in 19 and 8 cases, respectively, though their overall distributions remained largely distinct from the equivalence zone. At *N* = 100 in the two-foci scenario (F), the Exponential kernel achieved predictively equivalent in 4 cases. In all remaining scenarios, the Δ*ELPD* distributions were well separated, with no overlap between competing kernels and the equivalence zone.

Posterior predictive checks (PPC) further validated model performance (Figure S8; Figure S9). Zero- and one-inflation proportions calibration remained stable across all the scenarios (*R*^2^ = 1.0; Figure S10). Global *R*^2^ showed a slight inverse relationship with sample size, decreasing from 0.547 ± 0.136 at *N* = 50 to 0.481 ± 0.048 at *N* = 500 in single-focus scenarios, while global RMSE improving from 0.110 ± 0.025 at *N* = 50 to 0.1055 ± 0.008 at *N* = 500 (Figure S10). Similar trends were observed across two-foci scenarios (Figure S10). This pattern reflects increasing capture of biological variability with denser sampling, larger samples better represent local stochastic variation, increasing total variance while simultaneously improving absolute prediction error. *R*^2^ and *RMSE* therefore convey complementary information and should be interpreted jointly.

Parameter estimation improved with sample size across both spatial configurations (Table 7). In the single-focus scenarios, the absolute mean bias for source location (*f_x_*_1_, *f_y_*_1_) and ZOIB parameters (*π*_0_, and *q*) decreased as sample size increased (Table 7). The benefit of increased data density was more evident in the two-foci scenarios. Source localization accuracy improved markedly, with the bias for *f_y_*_1_ decreasing from −0.23 at *N* = 50 to −0.04 at *N* = 500 (Table 7). Additionally, the precision parameter (*φ*) converged toward the true value (18.00), moving from 16.82 ± 2.63 at *N* = 50 to 17.95 ± 0.74 at *N* = 500 (Table 7). Both zero-inflation (*π*_0_) and one-inflation (*q*) parameters were recovered with near-zero bias and high coverage across the entire sample size gradient (Table 7).

**Table 7:**
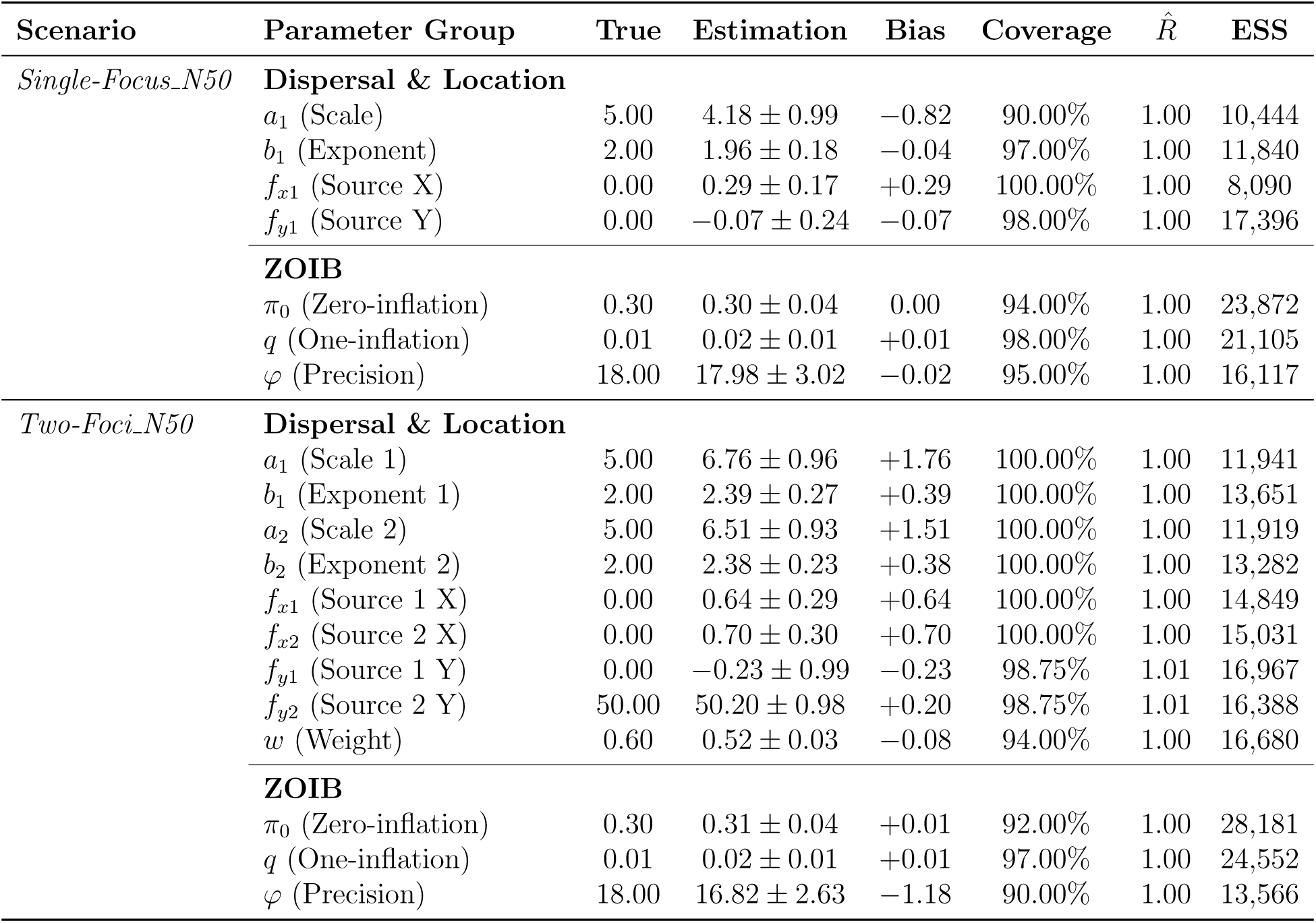
Performance of parameter estimation across gradients of spatial complexity and sample sizes (*N* = 50, 100, 250, 500). Values represent mean ± SD across 100 simulations. Here, we only show the scenarios of sample size *N* = 50, the full details please see Table S8.

### 3.2 Empirical Validation: Tracking Cucurbit Downy Mildew Dispersal in Managed Field Experiments

We validated HiBASIL under empirical conditions using field-scale experiments on cucurbit downy mildew arranged in a Randomized Complete Block Design across five treatment tiers, ranging from untreated controls to complex two-foci configurations. The ZOIB distribution effectively characterized the field disease severity data (*P >* 0.15 in 80% of treatments), providing a robust likelihood for Bayesian inference (Supplementary Note 2 S5).

We evaluated framework performance across three tiers: specificity in untreated controls, baseline accuracy in scenarios with known focus coordinates (Supplementary Note 3 S6), and joint inference of both dispersal parameters and unknown source locations. The following sections focus on the third tier.

#### Non-Treatment Controls: Assessment of Specificity

LOO-CV correctly identified the Null model as the most parsimonious fit in non-treatment control plots. Although complex spatial models exhibited slightly higher raw *R*^2^ values due to noise-fitting, their resulting spatial gradients were entirely flat and virtually indistinguishable from the Null baseline (Figure 12A–E; Figure S16; Supplementary Note 3 S6).

**Figure 12:**
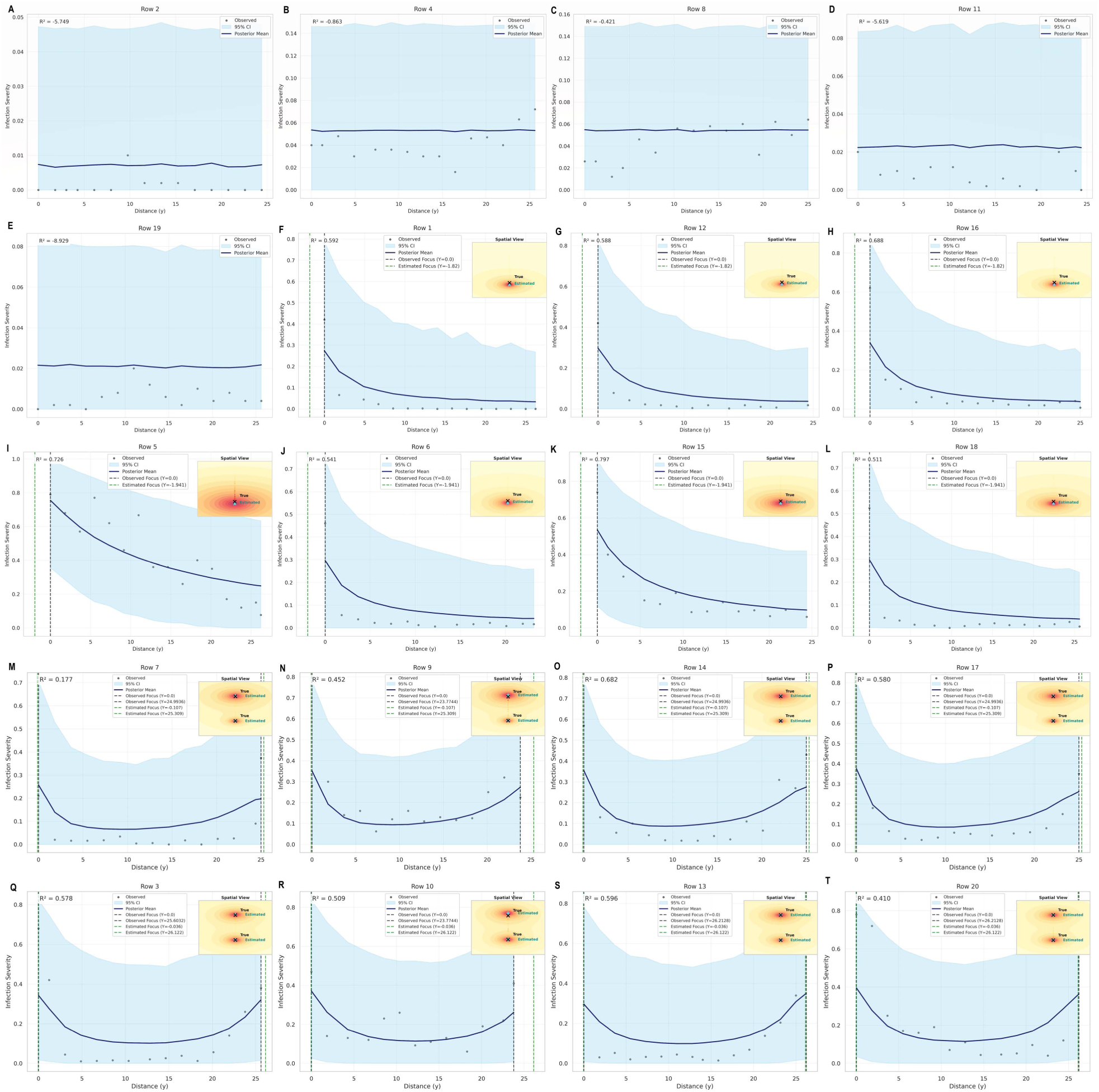
Spatial validation of posterior predictive distribution for treatments control, single-focus, and two-foci data. Model performance is evaluated at unknown focus location scenario. Panels represent non-treated controls (A-E), single-focus inoculated once (F-H), single-focus inoculated twice (I-L), two-foci inoculated once (M-P), and two-foci inoculated twice (Q-T) replicates. Primary panels display 1D spatial transects at *x* = 0, comparing observed severity (black points) against the posterior predictive mean (navy line) and 95% credible interval (shaded sky-blue region). Vertical green dashed lines mark the estimated locations of infection sources along the transect. Corresponding 2D insets visualize the spatial offset between the true observed source (cross) and inferred source location (star).

#### Single-Focus scenarios: Gradient Characterization and Estimates

##### Model Selection and Predictive Accuracy

The Power Law kernel consistently exhibited the best performance under unknown focus location across both inoculated-once and inoculated-twice treatments, with the highest ELPD_loo_ and lowest effective number of parameters (*p_loo_*) (Table S9). Furthermore, the Power Law kernel offered a substantial improvement in precision, reducing the overall RMSE and MAE compared to Exponential and Gaussian kernels. Overall *R*^2^ was 0.65 and 0.84 for inoculated-once and inoculated-twice treatments, respectively. Similar results were observed for the *R*^2^ of the continuous Beta component (Table S9; Figure S19I – P). Fixed source coordinates models yielded slightly higher predictive accuracy, supporting the selected kernel structure (Supplementary Note 3 S6).

##### Spatial Inference and Localization

Visual inspection of the spatial posterior distributions confirms the model’s robust ability to localize the unknown infection source (Figure 12F – L). Although jointly estimating the focus location increased model complexity, our strategy, which employ a global shared focus location with partial or complete pooling of dispersal parameters, achieved highly precise spatial inference. The *x*-coordinate (*f_x_*_1_) was estimated with negligible bias (0.01 to 0.02) and posterior standard deviation of 1.46 to 1.60 (prior: 3.0; Table 8). The y-coordinate (*f_y_*_1_) exhibited only slight bias (-1.82 to -1.94 m) with posterior standard deviation of 1.53-1.81 (prior: 10; Table 8).

**Table 8:**
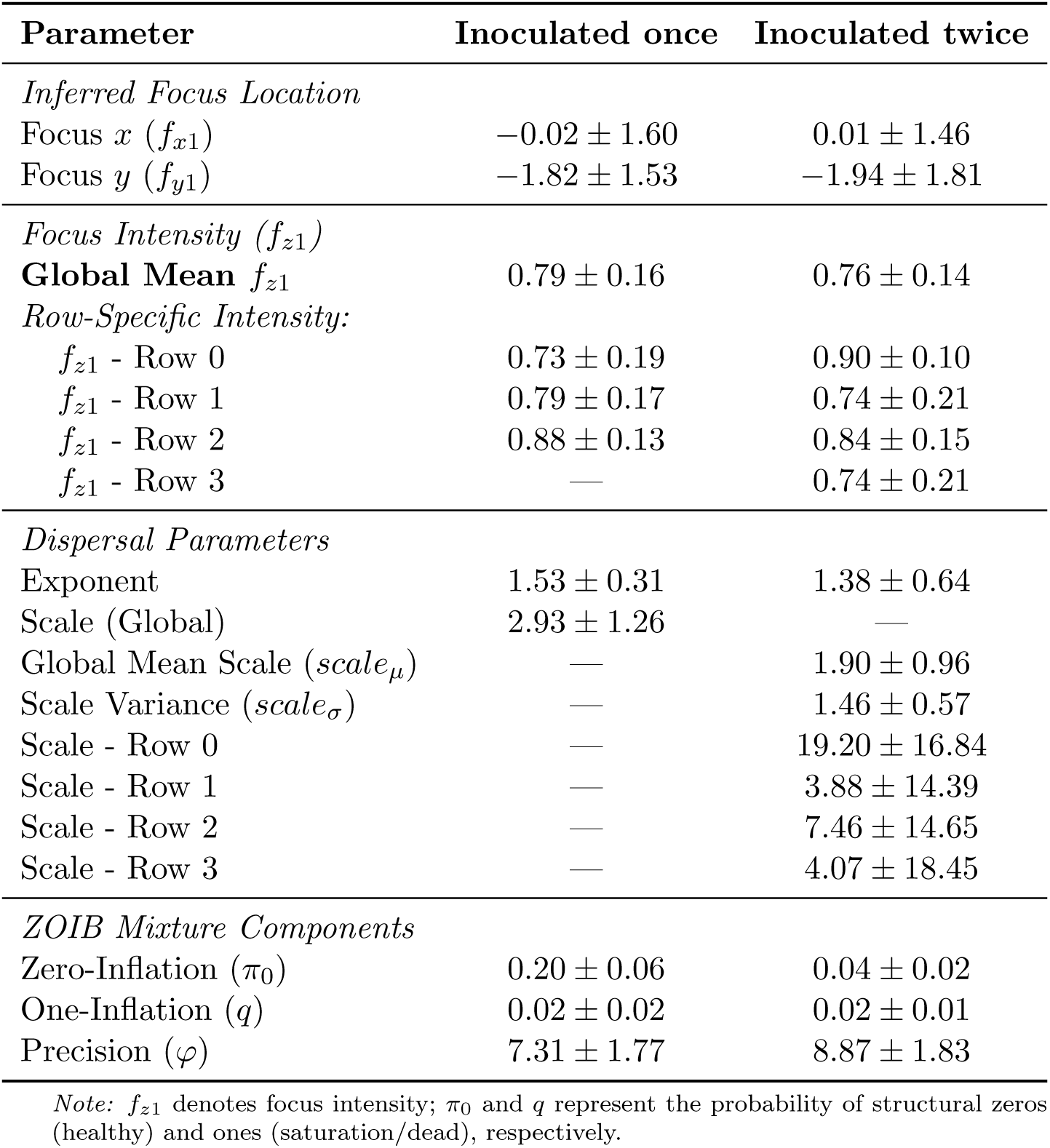
Parameter Estimation Results of Single-Focus for Cucurbit Downy Mildew Field Experiment Under Scenario of Unknown Focus Location (Joint Estimation).

| Parameter | Inoculated once | Inoculated twice |
| --- | --- | --- |
| *Inferred Focus Location* |  |  |
| Focus $x$ ($f_{x1}$) | $-0.02 \pm 1.60$ | $0.01 \pm 1.46$ |
| Focus $y$ ($f_{y1}$) | $-1.82 \pm 1.53$ | $-1.94 \pm 1.81$ |
| *Focus Intensity ($f_{z1}$)* |  |  |
| **Global Mean** $f_{z1}$ | $0.79 \pm 0.16$ | $0.76 \pm 0.14$ |
| *Row-Specific Intensity:* |  |  |
| $f_{z1}$ - Row 0 | $0.73 \pm 0.19$ | $0.90 \pm 0.10$ |
| $f_{z1}$ - Row 1 | $0.79 \pm 0.17$ | $0.74 \pm 0.21$ |
| $f_{z1}$ - Row 2 | $0.88 \pm 0.13$ | $0.84 \pm 0.15$ |
| $f_{z1}$ - Row 3 | — | $0.74 \pm 0.21$ |
| *Dispersal Parameters* |  |  |
| Exponent | $1.53 \pm 0.31$ | $1.38 \pm 0.64$ |
| Scale (Global) | $2.93 \pm 1.26$ | — |
| Global Mean Scale ($scale_\mu$) | — | $1.90 \pm 0.96$ |
| Scale Variance ($scale_\sigma$) | — | $1.46 \pm 0.57$ |
| Scale - Row 0 | — | $19.20 \pm 16.84$ |
| Scale - Row 1 | — | $3.88 \pm 14.39$ |
| Scale - Row 2 | — | $7.46 \pm 14.65$ |
| Scale - Row 3 | — | $4.07 \pm 18.45$ |
| *ZOIB Mixture Components* |  |  |
| Zero-Inflation ($\pi_0$) | $0.20 \pm 0.06$ | $0.04 \pm 0.02$ |
| One-Inflation ($q$) | $0.02 \pm 0.02$ | $0.02 \pm 0.01$ |
| Precision ($\varphi$) | $7.31 \pm 1.77$ | $8.87 \pm 1.83$ |

*Note:* $f_{z1}$ denotes focus intensity; $\pi_0$ and $q$ represent the probability of structural zeros (healthy) and ones (saturation/dead), respectively.

##### Biological Interpretation of Parameter Estimates

The estimated probability of structural zeros (healthy plants, *π*_0_) dropped from 0.20 in the inoculated-once group to 0.04 in the inoculated-twice group (Table 8), indicating substantially greater infection establishment under double inoculation. The inoculated-twice treatment was associated with a shallower dispersal gradient (i.e., further relative spread), indicated by a lower exponent (decay) estimate of 1.38 compared to 1.53 in the inoculated-once treatment (Table 8; Figure 12F – L). Furthermore, while the inoculated-once treatment yielded a global dispersal scale of 2.93, the inoculated-twice treatment exhibited high variance among row-specific scales (ranging from 3.88 to 19.20), reflecting the complex micro-environmental heterogeneity captured by the hierarchical structure (Table 8).

#### Two-Foci: Resolution of Overlapping Kernels

##### Model Selection and Spatial Inference

The Power Law kernel was the top-ranked model across all two-foci treatments (inoculated-once and inoculated-twice), with the highest Expected Log Predictive Density (ELPD_loo_) and minimum overall RMSE (Table S10).

Using a globally shared focus location with complete pooling of dispersal parameters, Hi-BASIL achieved a strong posterior contraction for all inferred source locations (Figure 12M– T; Figure S20I–P). Estimated *x*-coordinates (*f_x_*_1_ and *f_x_*_2_) showed negligible absolute bias (0.00 to 0.01) and a significant reduction in standard deviation from a prior of 3.0 to a posterior of 0.56 to 1.61 (Table 9). Estimated *y*-coordinates were recovered with highly accuracy. Estimated *f_y_*_1_ showed negligible absolute bias (0.04 to 0.11) and a reduction in uncertainty, with standard deviations narrowing from a prior of 10.0 to a posterior of 0.74 to 0.95 (Table 9). Estimated *f_y_*_2_ was very close to the true values (25.31 to [25, 23.8, 25, 25] for the inoculated-once treatment; 26.12 to [25.6, 23.8, 26.2, 26.2] for the inoculated-twice treatment), and with a reduction in standard deviation from a prior of 10.0 to a posterior of 1.09 to 1.69 (Table 9).

**Table 9:** Parameter Estimation Results of Two-Foci Models for Cucurbit Downy Mildew Field Experiment Under Scenario of Unknown Focus Location (joint estimation).

| Parameter | Inoculated once | Inoculated twice |
| --- | --- | --- |
| *Inferred Focus Locations* |  |  |
| Focus 1 $x$ ($f_{x1}$) | $0.00 \pm 0.56$ | $0.00 \pm 0.76$ |
| Focus 1 $y$ ($f_{y1}$) | $-0.11 \pm 0.74$ | $-0.04 \pm 0.91$ |
| Focus 2 $x$ ($f_{x2}$) | $0.01 \pm 1.61$ | $-0.00 \pm 0.95$ |
| Focus 2 $y$ ($f_{y2}$) | $25.31 \pm 1.69$ | $26.12 \pm 1.09$ |
| Weight ($w$) | $0.52 \pm 0.09$ | $0.50 \pm 0.08$ |
| *Focus Intensity ($f_z$)* |  |  |
| **Global Mean** $f_{z1}$ | $0.75 \pm 0.14$ | $0.78 \pm 0.13$ |
| **Global Mean** $f_{z2}$ | $0.75 \pm 0.14$ | $0.81 \pm 0.12$ |
| *Row-Specific $f_{z1}$:* |  |  |
| $f_{z1}$ - Row 0 | $0.62 \pm 0.22$ | $0.80 \pm 0.17$ |
| $f_{z1}$ - Row 1 | $0.83 \pm 0.17$ | $0.87 \pm 0.14$ |
| $f_{z1}$ - Row 2 | $0.84 \pm 0.16$ | $0.69 \pm 0.20$ |
| $f_{z1}$ - Row 3 | $0.90 \pm 0.12$ | $0.93 \pm 0.10$ |
| *Row-Specific $f_{z2}$:* |  |  |
| $f_{z2}$ - Row 0 | $0.61 \pm 0.18$ | $0.82 \pm 0.16$ |
| $f_{z2}$ - Row 1 | $0.91 \pm 0.11$ | $0.90 \pm 0.12$ |
| $f_{z2}$ - Row 2 | $0.85 \pm 0.14$ | $0.87 \pm 0.13$ |
| $f_{z2}$ - Row 3 | $0.80 \pm 0.16$ | $0.89 \pm 0.12$ |
| *Dispersal Parameters* |  |  |
| Exponent 1 | $3.37 \pm 1.26$ | $2.35 \pm 1.09$ |
| Exponent 2 | $1.54 \pm 0.57$ | $1.88 \pm 0.85$ |
| Scale 1 | $4.87 \pm 1.47$ | $5.40 \pm 1.59$ |
| Scale 2 | $5.51 \pm 1.66$ | $5.55 \pm 1.67$ |
| *ZOIB Mixture Components* |  |  |
| Zero-Inflation ($\pi_0$) | $0.05 \pm 0.03$ | $0.02 \pm 0.02$ |
| One-Inflation ($q$) | $0.02 \pm 0.02$ | $0.01 \pm 0.01$ |
| Precision ($\varphi$) | $8.11 \pm 1.60$ | $5.87 \pm 1.11$ |

##### Biological Interpretation: Intensity and Symmetry

Parameter estimates revealed distinct biological differences between treatments (Table 9). The inoculated-twice treatment resulted in a higher estimated global mean source intensity (*f_z_*_1_ = 0.78 and *f_z_*_2_ = 0.81) compared to the inoculated-once (*f_z_*_1_ = 0.75 and *f_z_*_2_ = 0.75). Furthermore, the increased inoculum load significantly reduced the proportion of structural zeros (healthy plants, *π*_0_), which decreased from 0.05 in the inoculated-once group to 0.02 in the inoculated-twice treatment. This indicates an almost complete infection of all plants within the plot.

The mixture weight parameter (*w*) and spatial kernel parameters revealed distinct dispersal dynamics between treatments. In the inoculated-once treatment, we observed marked asymmetry, with Focus 1 exhibiting a short-range, concentrated dispersal profile (exponent *b*1 = 3.37), while Focus 2 exhibiting a more diffuse dispersal profile (exponent *b*2 = 1.54) (Table 9). This heterogeneity was reflected in the mixing weight (*w* ≈ 0.52), which suggests a slight dominance of the first focus in local infection contribution. In contrast, the inoculated-twice treatment exhibited more symmetric dispersal profiles between the two foci (exponents of 2.35 and 1.88, respectively) and a perfectly balanced mixing weight (*w* ≈ 0.50) (Table 9).

### 3.3 Cross-Domain Portability and Multistage Validation: The Historical Cholera Outbreak

We applied HiBASIL to the 1854 London cholera epidemic data across four validation stages: null hypothesis rejection, blind source localization, parsimonious testing, and diagnostic assessment under mechanistic misspecification.

#### Null Hypothesis Rejection

The null model (*H*_0_) exhibited the lowest predictive density (ELPD) across all candidates and was decisively rejected in favor of spatially explicit models (Table S11), confirming the presence of a localized infection driver rather than random background mortality.

#### Blind Source Discovery (***H*_1_**)

Despite weakly informative priors, all the single-focus dispersal models successfully localized the high-mortality risk area within 42 m of the Broad Street pump (Table S11). While the Gaussian minimized information loss, the Power Law model was selected for its superior localization accuracy and statistical equivalent to the best performing model (Δ*ELPD <* 2 × *SE*; Table S11). The single-focus Power Law model exhibited a localization bias of 32.65 m relative to the true pump location (Table 10; Figure 13A&C). This represents precisely localization, particularly given that the model successfully overcame the network-based waterborne transmission, socioeconomic clustering, and building density introducing deviations from radial distance decay.

**Figure 13:**
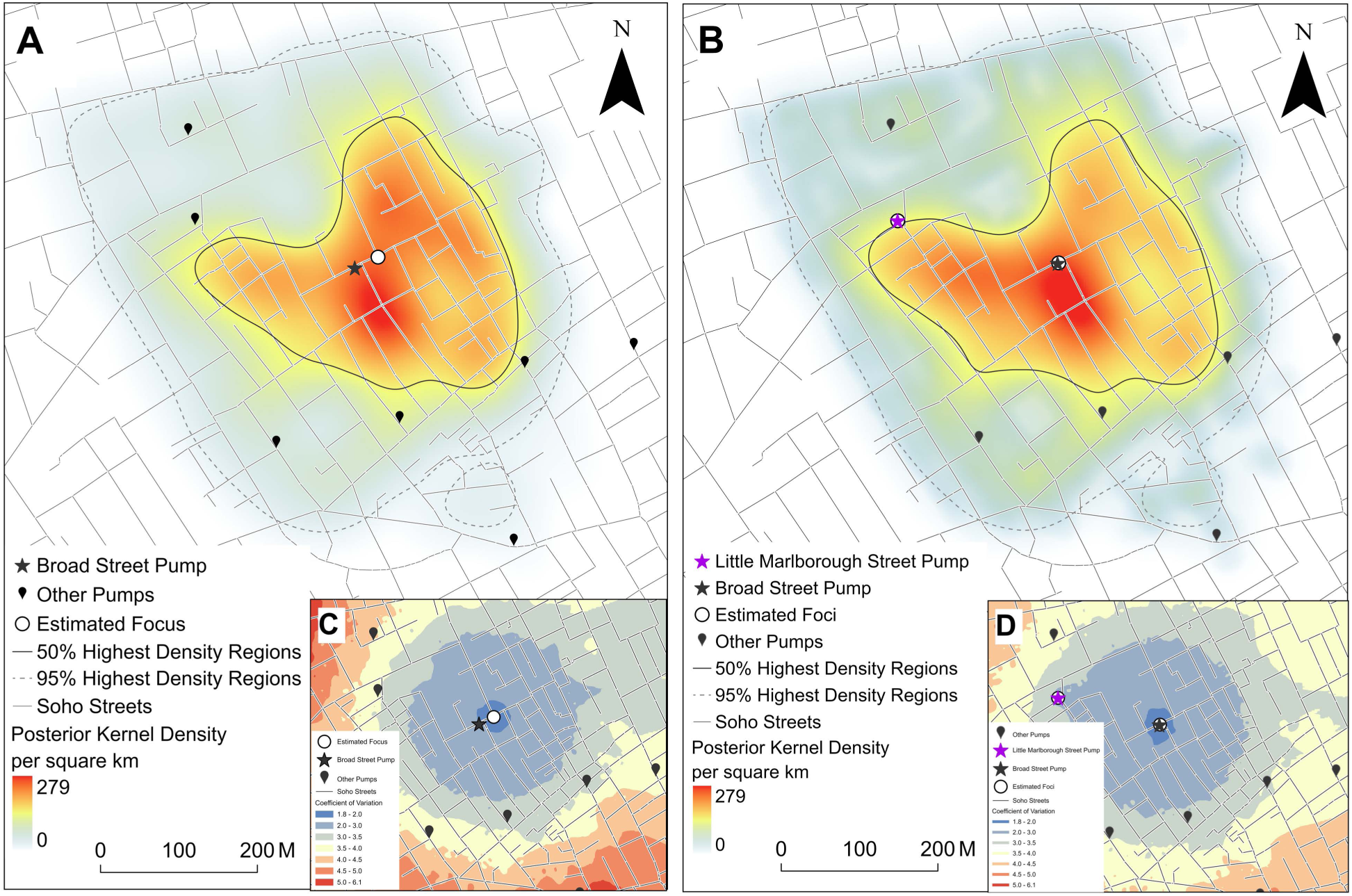
Spatial visualization of posterior predictive distribution and diagnostic uncertainty quantification for the 1854 cholera outbreak. Panels compare the single-focus blind discovery (A, C) with the two-foci parsimony stress test (B, D). **(A, B)** Posterior mean kernel density estimates of predicted mortality intensity (transparent = low, yellow = moderate, red = high) with 50% and 95% highest density regions (HDRs). Black star marks the Broad Street pump, the purple star marks the Little Marlborough street pump; open circles mark the estimated source locations, and teardrops markers indicate other historical pump locations. Despite visual similarity, leave-one-out cross-validation strongly favored the single-source model (Table S11). **(C, D)** Coefficient of Variation (*σ/µ*) surfaces quantifying the spatial reliability of predictions, interpolated by inverse distance weighting. Low CV (blue) over the outbreak core indicates high reliability, while warmer colors toward the periphery indicate lower reliability.

**Table 10:**
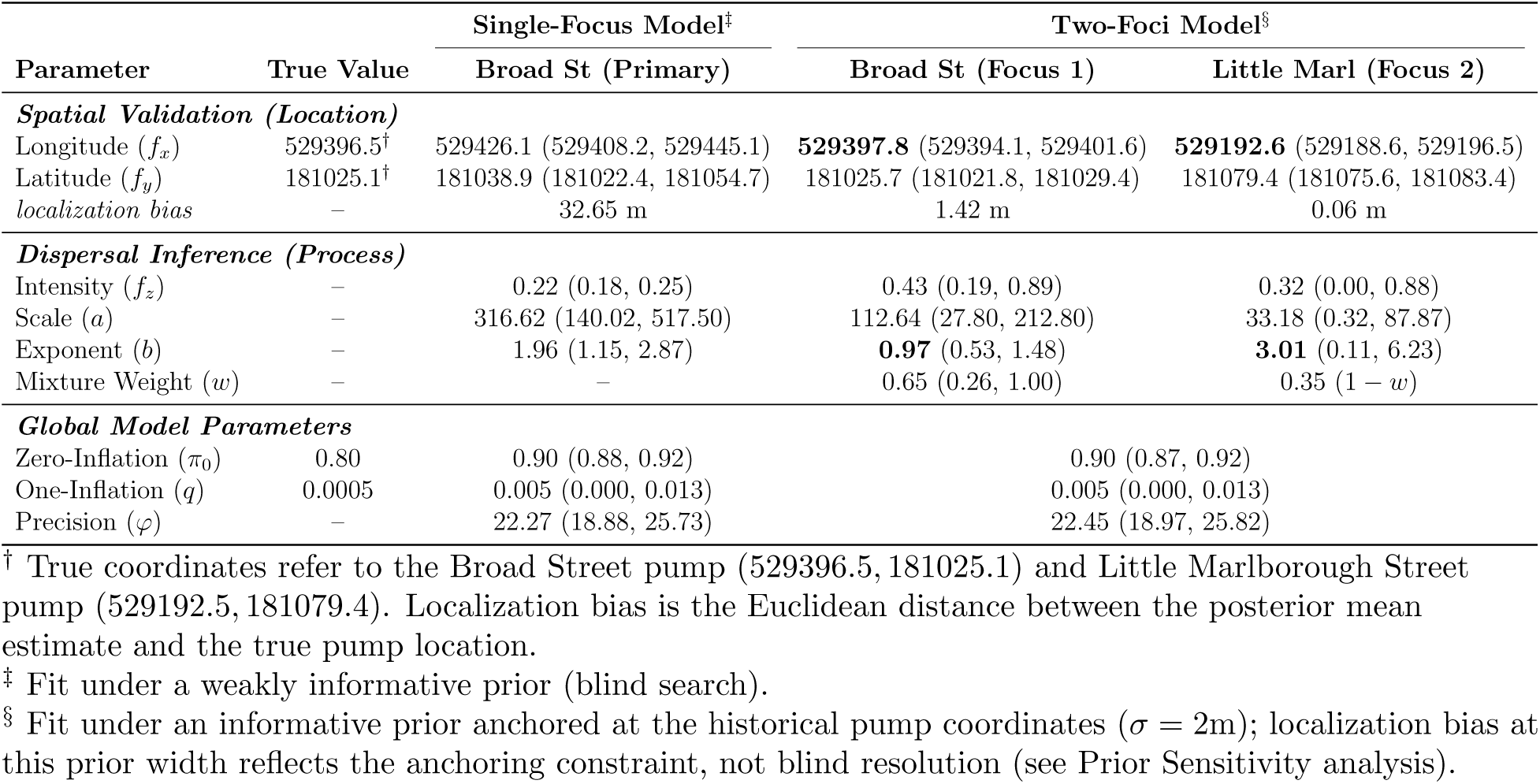
Comparative posterior parameter estimates for single- and two-foci Power Law models. Spatial coordinates (*f_x_, f_y_*) are validated against true pump locations. Under the tight prior anchoring the Two-Foci model (*σ* = 2*m*), posterior mean positions converge near both pump coordinates; this reflects the anchoring constraint rather than independent recovery, not improved localization precision. Despite this favorable anchoring, the dispersal shape and mixture parameters estimated by the Two-Foci model support the rejection of *H*_2_ by LOO-CV (Table S11). Abbreviation: CI, 95% Credible Interval.

| Parameter | True Value | Single-Focus Model<sup>‡</sup> | Two-Foci Model<sup>§</sup> |  |
| --- | --- | --- | --- | --- |
|  |  | Broad St (Primary) | Broad St (Focus 1) | Little Marl (Focus 2) |
| ***Spatial Validation (Location)*** |  |  |  |  |
| Longitude ($f_x$) | 529396.5<sup>†</sup> | 529426.1 (529408.2, 529445.1) | **529397.8** (529394.1, 529401.6) | **529192.6** (529188.6, 529196.5) |
| Latitude ($f_y$) | 181025.1<sup>†</sup> | 181038.9 (181022.4, 181054.7) | 181025.7 (181021.8, 181029.4) | 181079.4 (181075.6, 181083.4) |
| *localization bias* | – | 32.65 m | 1.42 m | 0.06 m |
| ***Dispersal Inference (Process)*** |  |  |  |  |
| Intensity ($f_z$) | – | 0.22 (0.18, 0.25) | 0.43 (0.19, 0.89) | 0.32 (0.00, 0.88) |
| Scale ($a$) | – | 316.62 (140.02, 517.50) | 112.64 (27.80, 212.80) | 33.18 (0.32, 87.87) |
| Exponent ($b$) | – | 1.96 (1.15, 2.87) | **0.97** (0.53, 1.48) | **3.01** (0.11, 6.23) |
| Mixture Weight ($w$) | – | – | 0.65 (0.26, 1.00) | 0.35 ($1 - w$) |
| ***Global Model Parameters*** |  |  |  |  |
| Zero-Inflation ($\pi_0$) | 0.80 | 0.90 (0.88, 0.92) | 0.90 (0.87, 0.92) | |
| One-Inflation ($q$) | 0.0005 | 0.005 (0.000, 0.013) | 0.005 (0.000, 0.013) | |
| Precision ($\varphi$) | – | 22.27 (18.88, 25.73) | 22.45 (18.97, 25.82) | |

<sup>†</sup> True coordinates refer to the Broad Street pump (529396.5, 181025.1) and Little Marlborough Street pump (529192.5, 181079.4). Localization bias is the Euclidean distance between the posterior mean estimate and the true pump location.
<sup>‡</sup> Fit under a weakly informative prior (blind search).
<sup>§</sup> Fit under an informative prior anchored at the historical pump coordinates ( $\sigma = 2m$ ); localization bias at this prior width reflects the anchoring constraint, not blind resolution (see Prior Sensitivity analysis).

#### Parsimony Validation (***H*_2_**): the Failure of the Two-foci Hypothesis

Despite priors deliberately constructed to favor the two-foci alternative with highly informative priors (*σ* = 2*m*) centered on the Broad Street and Little Marlborough Street pumps (Figure 13B&D), LOO-CV rejected the mixture model in favor of the parsimonious single-focus explanation (*H*_1_) (Table S11). This constitutes a stress test in the strictest sense, even give the sources’ exact coordinates as a strong prior, the data provided insufficient evidence to retain it. Within the two-foci model group, the Power Law model was selected for subsequent analyses (Table S11).

Parameter diagnostics revealed why the secondary focus was appropriately rejected. The Little Marlborough Street pump focus exhibited: (1) extreme decay exponent (*b*2 ≈ 3.01, 95% CI [0.11, 6.23]) indicating singularity-like spatial drop-off inconsistent with waterborne dispersal, compared to the diffuse gradient from the Broad Street focus (*b*1 ≈ 0.97, 95% CI [0.53, 1.48]); (2) a mixture weight extends to 1.0 (95% CI [0.26, 1.00]), placing complete collapse to a single-source within the credible range; and (3) intensity 95% CI [0.0, 0.88] encompassing zero contribution. Under tight prior, the posterior mean positions of both foci converged near their respective anchor coordinates (positional error 1.42 m and 0.06 m for the Broad Street and Little Marlborough foci, respectively; Table 10). Given the *σ* = 2 m prior width, this convergence reflects the strength of the anchoring constraint rather than an independent recovery of either source’s location, and does not indicate improved localization precision relative to the blind *H*_1_ estimate (32.65 m). These results indicate the secondary focus carries no genuine mortality signal, consistent with Snow’s attribution of nearby mortality to resident preference for the Broad Street pump rather than secondary pump contamination (Snow, John, 1855).

#### Prior Sensitivity

The Power Law kernel was statistically predictive equivalent to the top-ranked Gaussian (|ΔELPD| *<* 2×*SE*) across all prior widths and achieved consistently lower localization error than both alternative kernels at every evaluated width (Table S12), with an improvement of 8.95 m over Gaussian in the blind search scenario (Gaussian 41.69 m and Power Law 32.74 m at *σ* = 250*m*). The Power Law kernel was therefore selected for subsequent analysis.

The sensitivity response of the Power Law dispersal model exhibited an inverse L-shaped trajectory characterized by two opposing regimes (Figure S11). In the data-dominant regime (*σ* ≥ 50*m*), representing a blind search, localization error reached a plateau at approximate 32.7 m. Increasing prior width from 50 m to 250 m resulted in a negligible variation (less than 1.0 m), confirming an intrinsic resolution limit where the model’s convergence is driven by the mortality signal rather than initialization bias. Conversely, in the prior-driven regime (*σ <* 50*m*), forensic precision below the block level (*<* 10*m*) was only achievable when the prior constraint exceeded the intrinsic resolution of the mortality records. We observed a rapid reduction in error as the prior tightened, reaching a global minimum of 7.5 m at *σ* = 5*m*. This sharp transition demonstrates that while the cholera mortality data is robust enough to identify the correct neighborhood, pinpointing the exact pump requires integrating external information to break the intrinsic stochastic noise.

#### Diagnostic Behavior Under Misspecification

HiBASIL achieved precise source localization (32.65 m error) despite low explanatory power (*R*^2^ ≈ 0.09; Table S13), indicating that in this case, source identification did not require comprehensive variance explanation, even though the model’s fundamental mechanistic assumptions are violated.

Spatial uncertainty analysis (Figure 13C & D) revealed heterogeneous predictive confidence. HiBASIL remained highly stable near the estimated source (low coefficient of variation), where the mortality signal was strongest. Conversely, relative uncertainty increased at the spatial periphery (high coefficient of variation). This pattern reflects robust model behavior, where confidence concentrates in high-density zones, while the instability inherent in predicting rare events at the data-sparse outbreak edge is correctly reflected as elevated variability.

The low *R*^2^ reflects a mechanistic mismatch rather than model failure. Waterborne transmission via street networks, socioeconomic clustering, and building density patterns introduce systematic variance that simple radial dispersal models cannot capture; because the radial form is misspecified for network-structured transmission, a high *R*^2^ from this model would more likely reflect overfitting to these spatial confounders than a correct account of the mechanism. Two lines of evidence support appropriate model behavior despite this mismatch. First, accurate zero-inflation recovery: the model correctly predicted 82.0% of locations, closely matching the 80.1% observed (Table S13), indicating it recovers the outbreak’s spatial boundary despite structural misspecification. Second, kernel equivalence: the Power Law, Exponential, and Gaussian kernels remained statistical indistinguishable in predictive performance across every elevated prior width (|ΔELPD| *<* 2×SE; Table S12), indicating the data do not strongly favor one radial decay profile over another. This appropriately reflects mechanistic uncertainty: when the true transmission geometry (network) violate the model’s radial assumption, the precise shape of the decay tail become secondary to the dominant structural error.

## 4 Discussion

HiBASIL integrates mechanistic dispersal kernels with a ZOIB likelihood within a hierarchical Bayesian framework, enabling simultaneous inference of source locations, dispersal mechanisms, and uncertainty from bounded spatial observations. Validation of HiBASIL using data from simulations, field experiments on cucurbit downy mildew, and reconstruction of the 1854 cholera outbreak, demonstrate robust performance across various ecological and epidemiological contexts.

The central challenge in multi-source localization is disentangling overlapping dispersal gradients. Heuristic clustering (Verity et al., 2014) cannot distinguish true sources from secondary spread, kernel density estimation (KDE) (Wand, 1994) over-smooths multi-source patterns, and spatial GLMs treat spatial autocorrelation as a phenomenological texture rather than a mechanistic trajectory (Dormann et al., 2007). Preliminary results from a systematic comparison of HiBASIL against GLM and KDE approaches indicate that standard methods substantially under-perform HiBASIL in multi-source localization (Guo, unpublished data). Recent optimization-based approaches such as Zhou et al. (2026) have demonstrated effective multi-source localization in discrete network setting but require candidate source sets to be specified *a priori* and are designed for network topology rather than continuous spatial fields. HiBASIL addresses these gaps by operating in continuous space and evaluating competing source configurations via LOO-CV with two constraints. First, mechanistic dispersal kernels impose biological geometry that improves identifiability, achieving 100% correct model selection in 2,300 correctly specified simulations. Second, the ZOIB likelihood preserves the tripartite data structure (structural zeros, continuous intensity, and structural ones) preventing boundary blurring common to standard transformations (Tang et al., 2023). Together, these enable the detection of minority sources contributing as little as 5% of the total signal while maintaining robust parameter recovery across sample sizes.

The consistent selection of the Power Law kernel across single- and two-foci field scenarios aligns with published dispersal studies for cucurbit downy mildew. Ojiambo et al. (2017) analyzed cucurbit downy mildew epidemics across multiple years in the eastern US, finding Power Law spread parameter *b* ranging from 1.51 to 4.16 across epidemic years, closely matching our empirical estimates (*b* = 1.38 − 1.53 for single-focus and *b* = 1.54 − 3.37 for two-foci treatments under unknown conditions). This is further consistent with empirical observations of accelerating epidemic waves driven by long-distance dispersal, where Power Law exponents consistently fall within a narrow range of 1.74 - 2.36 regardless of spatial scale (Mundt et al., 2009; Ojiambo et al., 2017). The selection of Power Law over Exponential and Gaussian kernels suggests that standard diffusion-based models may underrepresent the tail dispersal probability in the cucurbit downy mildew pathosystem.

The 32.65 m blind localization error in the application of HiBASIL to the historical cholera data can be contextualized against geographic profiling (Le Comber et al., 2011), which also successfully identified the Broad Street pump as the most likely source (top 0.2% of the geoprofile) but required 13 pre-identified candidate neighborhood pumps locations as input. HiBASIL achieves comparable source identification without requiring prior candidate lists, estimating source coordinates continuously from spatial intensity data, a blind localization that remains accurate despite a radial dispersal assumption that mechanistically misspecifies the network-based transmission underlying the cholera outbreak. That accurate localization persists under low explanatory power (*R*^2^ = 0.09) reframes the model’s role from complete system description to targeted signal isolation with transparent uncertainty. This capacity is particularly relevant for biosecurity and invasion biology (Chapple et al., 2022), where resolving unknown minority sources is essential for early intervention. More broadly, the framework’s signaling of mechanistic ambiguity via kernel equivalence and elevated uncertainty under misspecification supports reliable decision-making across a range of inverse problems, from tracking pollution in heterogeneous flow fields to reconstructing invasion pathways in fragmented landscapes.

Performance depends critically on alignment between the dispersal scale and the observation window (Nathan et al., 2012): narrow steep gradients (local epidemic) require suitable spatial coverage to avoid data sparsity that dilutes the mechanistic signal; while wide shallow gradients require an extended sampling area to capture tail behavior necessary for kernel discrimination and prevent truncation bias. When these conditions are not met, HiBASIL provides transparent warnings rather than producing misleading estimates. Under null scenarios lacking genuine spatial structure, dispersal-related posteriors remain informational equivalent to priors, demonstrating that the model defaults to regularization rather than inferring spurious sources. When reality complexity and model specification deviates substantially, the framework demonstrates excellent self-diagnostic capacity: under-parameterization (hidden sources) manifests through posterior bimodality (Betancourt, 2018) and convergence failure (*Ȓ* > 1.01), while over-parameterization triggers parameter collapse. Sample size analysis established viable inference at *N* = 50 with 100% kernel identification maintained, though predictive equivalence emerged in 19% of sparse sampling two-foci scenarios being resolved completely by *N* = 100. Hierarchical pooling across replicates in field experiments obtained accurate multi-focus localization and also confirmed operational robustness under sparse sampling.

Several structural assumptions define the current scope of HiBASIL. First, the isotropic dispersal assumption degrades its performance in strongly anisotropic systems driven by prevailing winds, topographic channeling, or host resistance distributions (Soubeyrand et al., 2008); future integration of environmental covariates (Gorton and Shaw, 2023) would address this. Second, the standard kernel library (Exponential, Gaussian, Power Law) cannot represent stratified dispersal (e.g., combining local splash and long-distance transport), network-based transmission, or vector-mediated spread, which require alternative functional forms or hierarchical extensions. Third, extending the framework to spatial-temporal dynamics or integrating genetic data (Polonsky et al., 2019) could resolve ambiguities in overlapping signatures that spatial data alone cannot distinguish. Finally, operational utility in blind scenarios would be enhanced by formal source-count selection; candidate approaches include sparsity-inducing shrinkage priors (e.g., the Horseshoe prior) (Piironen and Vehtari, 2017), which prune redundant source parameters automatically, or Bayesian stacking (Yao et al., 2018) to weight predictions from competing multi-source models.

HiBASIL provides a principled framework for jointly estimating multi-source locations and mechanistic dispersal kernels from bounded spatial observations, demonstrating that appropriate handling of zero-one-inflation and hierarchical structure enables precise inference even from sparse, noisy real-world data. As global change accelerates complex, multi-focal introductions across ecological and epidemiological systems, the capacity to rapidly reconstruct transmission origins from limited surveillance will become increasingly valuable. The framework’s domain-agnostic design makes it applicable across plant disease management, human epidemiology, environmental monitoring, and invasion biology.

## Supporting information

supplementary manuscript

## Acknowledgment

We gratefully acknowledge Jophr Galian, Isaac Kikway, and Roden Lizardo for their help in the field experiment. We also thank John Garner (Williamsdale Biofuels Field Lab and Horticultural Crops Research Station, NC State University) and Stewart Biegler (North Carolina Department of Agriculture & Consumer Services) for their support with the field experiment. This research was supported by the Ecology and Evolution of Infectious Diseases (EEID) program jointly funded by NSF and USDA NIFA, United States. David Gent was also supported by the U.S. Department of Agriculture CRIS project 2072-21000-061-000-D.

## Data and Code Availability

The codes, epidemiological datasets, results, as well as the software tutorial are available in the GitHub repository HiBASIL.

