## supplementary manuscript for "HiBASIL: Hierarchical Bayesian Source Inference and Localization for Spatial Epidemiology"

### Supporting information

Table S1: **Parameter specifications and scenario-specific priors for dispersal kernel geometry.**

| Scenario | Parameter | True Value | Prior Distribution | Scenario Description |
|---|---|---|---|---|
| Base | Scale ($a$)<br>Exponent ($b$) | 5<br>2 | $\text{Gamma}(2.0, 0.4)$<br>$\text{Gamma}(4.0, 2.0)$ | Moderate reach,<br>standard decay |
| Narrow & Steep | Scale ($a$)<br>Exponent ($b$) | 1<br>3 | $\text{Gamma}(1.5, 1.5)$<br>$\text{Gamma}(9.0, 3.0)$ | Localized spread,<br>rapid drop-off |
| Wide & Steep | Scale ($a$)<br>Exponent ($b$) | 15<br>3 | $\text{Gamma}(5.0, 0.33)$<br>$\text{Gamma}(9.0, 3.0)$ | Extended reach,<br>sharp boundary |
| Wide & Shallow | Scale ($a$)<br>Exponent ($b$) | 15<br>1.5 | $\text{Gamma}(5.0, 0.33)$<br>$\text{Gamma}(3.0, 2.0)$ | Long-distance spread,<br>fat-tailed |

Table S2: Prior specifications for field experiment control data analysis across the Null, Single-focus, and Two-foci models

| Model | Parameter | Prior Distribution |
|---|---|---|
| Null Model | Severity ($f_{z1}^{\text{base}}$) | $\text{Beta}(1, 11)$ |
| | $\sigma_{\text{row}}$ | $\text{HalfNormal}(\sigma = 0.2)$ |
| | One-inflation ($q$) | $\text{Beta}(1, 50)$ |
| | Zero-inflation ($\pi_0$) | $\text{Beta}(1, 1, \text{shape} = 5)$ |
| | Precision ($\varphi$) | $\text{Beta}(2, 0.5)$ |
| Single-focus Model | x-coordinate ($f_{x1}$) | $\text{Normal}(\mu = 0, \sigma = 5.0, \text{shape} = 5)$ |
| | y-coordinate ($f_{y1}$) | $\text{Normal}(\mu = 13, \sigma = 10.0, \text{shape} = 5)$ |
| | Severity ($f_{z1}^{\text{base}}$) | $\text{Beta}(1, 11)$ |
| | $\sigma_{\text{row}}$ | $\text{HalfNormal}(\sigma = 0.2)$ |
| | Scale ($a_1$) | $\text{Gamma}(3, 0.5)$ |
| | Exponent ($\text{b}_1$) | $\text{Gamma}(4, 2.0)$ |
| | One-inflation ($q$) | $\text{Beta}(1, 50)$ |
| | Zero-inflation ($\pi_0$) | $\text{Beta}(1, 1, \text{shape} = 5)$ |
| | Precision ($\varphi$) | $\text{Beta}(2, 0.5)$ |
| Two-foci Model | x-coordinate ($f_{x1}$) | $\text{Normal}(\mu = 0, \sigma = 5.0, \text{shape} = 5)$ |
| | y-coordinate ($f_{y1}$) | $\text{Normal}(\mu = 8, \sigma = 10.0, \text{shape} = 5)$ |
| | x-coordinate ($f_{x2}$) | $\text{Normal}(\mu = 0, \sigma = 5.0, \text{shape} = 5)$ |
| | y-coordinate ($f_{y2}$) | $\text{Normal}(\mu = 18, \sigma = 10.0, \text{shape} = 5)$ |
| | Severity 1 ($f_{z1}^{\text{base}}$) | $\text{Beta}(1, 11)$ |
| | Severity 2 ($f_{z2}^{\text{base}}$) | $\text{Beta}(1, 11)$ |
| | $\sigma_{\text{row}}$ | $\text{HalfNormal}(\sigma = 0.2)$ |
| | Scale 1 ($a_1$) | $\text{Gamma}(3, 0.5)$ |
| | Exponent 1 ($b_1$) | $\text{Gamma}(4, 2.0)$ |
| | Scale 2 ($a_2$) | $\text{Gamma}(3, 0.5)$ |
| | Exponent 2 ($b_2$) | $\text{Gamma}(4, 2.0)$ |
| | Weight ($w$) | $\text{Beta}(2, 2)$ |
| | One-inflation ($q$) | $\text{Beta}(1, 50)$ |
| | Zero-inflation ($\pi_0$) | $\text{Beta}(1, 1, \text{shape} = 5)$ |
| | Precision ($\varphi$) | $\text{Beta}(2, 0.5)$ |

Table S3: Model priors and parameter specification across treatments. Source locations were evaluated as either **Known** (fixed to physical measurements) or **Unknown** (estimated with weakly informative priors). Parameters are classified by scope: **Global** (shared across replicates) or **Row-Specific** (hierarchical).

| Treatment | Parameter | Scope | Prior / Definition |
|---|---|---|---|
| Single-focus (Inoculated once) | $f_x, f_y$ | Global | **Known:** Fixed at 0 \| **Unknown:** $\text{Normal}(0,3)$ / $\text{Normal}(0,10)$ |
| | $\text{scale}_1$ | Global | $\text{Gamma}(2,2)$ |
| | $\text{exponent}_1$ | Global | $\text{Gamma}(15,2)$ |
| | $f_{z1\_base}$ | Global | $\text{Beta}(1,1)$ |
| | $\sigma_{row}$ | Hyperparam | $\text{HalfNormal}(0.2)$ |
| | $f_{z1\_row}$ | Row-Specific | $\text{Normal}(0,\sigma_{row})$ |
| | ZOIB params | Global | $q \sim \text{Beta}(1,10), \pi_0 \sim \text{Beta}(2,5), \varphi \sim \text{Gamma}(4,2)$ |
| Single-focus (Inoculated twice) | $f_x, f_y$ | Global | **Known:** Fixed at 0 \| **Unknown:** $\text{Normal}(0,3)$ / $\text{Normal}(0,10)$ |
| | $\text{scale}_\mu, \sigma$ | Hyperparams | $\text{Gamma}(2,2)$, $\text{HalfNormal}(1.2)$ |
| | $\text{scale}_{raw}$ | Row-Specific | $\text{Normal}(0,1)$ |
| | $\text{scale}_{row}$ | Deterministic | $\text{scale}_\mu \cdot \exp(\text{scale}_\sigma \cdot \text{scale}_{raw})$ |
| | exponent | Global | $\text{Gamma}(10,2)$ |
| | $f_{z1\_base}$ | Global | $\text{Beta}(2,2)$ |
| | $\sigma_{row}$ | Hyperparam | $\text{HalfNormal}(0.3)$ |
| | $f_{z1\_row}$ | Row-Specific | $\text{Normal}(0,\sigma_{row})$ |
| | ZOIB params | Global | $q, \pi_0 \sim \text{Beta}(1,10), \varphi \sim \text{Gamma}(4,2)$ |
| Two-foci (Inoculated once / twice) | $f_{x1}, f_{y1}$ | Global | **Known:** Fixed at 0 \| **Unknown:** $\text{Normal}(0,3)$ / $\text{Normal}(0,10)$ |
| | $f_{x2}$ | Global | **Known:** Fixed at 0 \| **Unknown:** $\text{Normal}(0,3)$ |
| | $f_{y2}$ | Global | **Known:** Inoc 1: $[25, 23.8, 25, 25]$<br>Inoc 2: $[25.6, 23.8, 26.2, 26.2]$<br>**Unknown:** $\text{Normal}(25,10)$ |
| | $\text{scale}_{1,2}$ | Global | $\text{Gamma}(6,2)$ |
| | $\text{exponent}_{1,2}$ | Global | $\text{Gamma}(12,2)$ |
| | $f_{z1\_base}$ | Global | $\text{Beta}(2,2)$ |
| | $f_{z2\_base}$ | Global | $\text{Beta}(2,2)$ |
| | $\sigma_{row}$ | Hyperparam | $\text{HalfNormal}(0.25)$ |
| | $f_{z1\_row}$ | Row-Specific | $\text{Normal}(0,\sigma_{row})$ |
| | $f_{z2\_row}$ | Row-Specific | $\text{Normal}(0,\sigma_{row})$ |
| | Weight ($w$) | Global | $\text{Beta}(2,2)$ |
| | ZOIB params | Global | $q, \pi_0 \sim \text{Beta}(1,10), \varphi \sim \text{Gamma}(4,2)$ |

Note: All models reached convergence.
[a] $\pi_0$ is the base proportion of 0.
[b] $q$ is the proportion of 1 given data greater than 0.
[c] $\varphi$ is the precision of ZOIB distribution.

Table S4: **Prior distributions specification for spatial models of the 1854 Soho cholera epidemic data.** The table details the prior settings for the Null, Single-Focus, and Two-Foci models. Notably, the Two-Foci specification employs highly informative priors for location parameters ($f_x, f_y$) centered on specific pump coordinates ($\sigma = 2m$), whereas the Single-Focus models use weakly informative spatial priors ($\sigma = 250m$ to allow for broad search). *Notation:* $N(\mu, \sigma)$, Normal distribution; Gamma$(\alpha, \beta)$, Gamma distribution; Beta$(\alpha, \beta)$, Beta distribution.

| Scenario | Parameter | Symbol | Prior Distribution |
|---|---|---|---|
| **Null Model** | Intensity (Background) | $f_z$ | $\text{Beta}(1, 1)$ |
| | Zero-inflation | $\pi_0$ | $\text{Beta}(2, 2)$ |
| | One-inflation | $q$ | $\text{Beta}(1, 10)$ |
| | Precision | $\varphi$ | $\text{Gamma}(2, 0.1)$ |
| **Single-Focus** | Longitude | $f_x$ | $\text{Normal}(529346, 250)$ |
| | Latitude | $f_y$ | $\text{Normal}(180983, 250)$ |
| | Intensity (Focus) | $f_z$ | $\text{Beta}(1, 1)$ |
| | Scale | $a$ | $\text{Gamma}(3, 0.015)$ |
| | Exponent | $b$ | $\text{Gamma}(10, 5)$ |
| | Zero-inflation | $\pi_0$ | $\text{Beta}(2, 2)$ |
| | One-inflation | $q$ | $\text{Beta}(1, 10)$ |
| | Precision | $\varphi$ | $\text{Gamma}(2, 0.1)$ |
| **Two-Foci** | Longitude (Focus 1) | $f_{x1}$ | $\text{Normal}(529396.5, 2)$ |
| | Latitude (Focus 1) | $f_{y1}$ | $\text{Normal}(181025.1, 2)$ |
| | Longitude (Focus 2) | $f_{x2}$ | $\text{Normal}(529192.5, 2)$ |
| | Latitude (Focus 2) | $f_{y2}$ | $\text{Normal}(181079.4, 2)$ |
| | Intensity (Focus 1, 2) | $f_{z1}, f_{z2}$ | $\text{Beta}(1, 1)$ |
| | Scale (Focus 1, 2) | $a_1, a_2$ | $\text{Gamma}(2, 0.04)$ |
| | Exponent (Focus 1, 2) | $b_1, b_2$ | $\text{Gamma}(2, 1)$ |
| | Mixture Weight | $w$ | $\text{Beta}(1, 1)$ |
| | Zero-inflation | $\pi_0$ | $\text{Beta}(2, 2)$ |
| | One-inflation | $q$ | $\text{Beta}(1, 10)$ |
| | Precision | $\varphi$ | $\text{Gamma}(2, 0.1)$ |

Table S5: Supplementary assessment of the impact of spatial domain scaling on parameter recovery. The comparison between the primary grid ($y = 50$) and scaled reference grids ($y = 8$ and $y = 75$) demonstrates the model's sensitivity to the alignment between biological dispersal reach and sampling resolution.

| **Scenario** | **Dispersal Characteristic** | **Primary Range** | **Reference Range** | **Inference Objective** |
|---|---|---|---|---|
| *Narrow & Steep* | Localized Spread ($a = 1, b = 3$) | $y = 50$ | $y = 8$ | Validates that parameter recovery (0% coverage at $y = 50$) is restored when the sampling resolution captures the localized spatial gradient. |
| *Wide & Shallow* | Long-Distance Fat-Tail ($a = 15, b = 1.5$) | $y = 50$ | $y = 75$ | Serves as a reference to confirm that capturing the distal dispersal tail stabilizes the estimation of the fat-tail exponent ($b$). |
| *Base Scenario* | Moderate Reach ($a = 5, b = 2$) | $y = 50$ | — | Establishes the optimal ratio of dispersal reach to domain size where recovery is most stable. |

Table S6: Parameter recovery results for scale-optimized scenarios: Narrow & Steep ($y = 8$) and Wide & Shallow ($y = 75$). Results are aggregated over 100 independent simulations using the Power Law kernel.

| **Scenario** | **Parameter Group** | **True** | **Estimate** | **Bias** | **Coverage** | $\hat{R}$ | **ESS (Bulk)** |
|---|---|---|---|---|---|---|---|
| *Narrow & Steep* ($y = 8$) | **Dispersal & Location** | | | | | | |
| | $a$ (Scale) | 1 | $0.84 \pm 0.10$ | $-0.16$ | 77.00% | 1.00 | 7,467 |
| | $b$ (Exponent) | 3 | $2.87 \pm 0.13$ | $-0.13$ | 88.00% | 1.00 | 9,513 |
| | $f_x$ (Source X) | 0 | $0.00 \pm 0.00$ | 0.00 | 100.00% | 1.00 | 6,047 |
| | $f_y$ (Source Y) | 0 | $0.00 \pm 0.01$ | 0.00 | 95.00% | 1.00 | 24,285 |
| | **ZOIB** | | | | | | |
| | $\pi_0$ (Zero-inflation) | 0.3 | $0.30 \pm 0.01$ | 0.00 | 96.00% | 1.00 | 25,193 |
| | $q$ (One-inflation) | 0.01 | $0.01 \pm 0.00$ | 0.00 | 97.00% | 1.00 | 24,980 |
| | $\varphi$ (Precision) | 18 | $18.31 \pm 1.46$ | $+0.31$ | 94.00% | 1.00 | 22,667 |
| *Wide & Shallow* ($y = 75$) | **Dispersal & Location** | | | | | | |
| | $a$ (Scale) | 15 | $12.51 \pm 2.10$ | $-2.49$ | 89.00% | 1.00 | 9,243 |
| | $b$ (Exponent) | 1.5 | $1.49 \pm 0.13$ | $-0.01$ | 98.00% | 1.00 | 9,480 |
| | $f_x$ (Source X) | 0 | $1.35 \pm 0.42$ | $+1.35$ | 100.00% | 1.00 | 5,706 |
| | $f_y$ (Source Y) | 0 | $-1.25 \pm 0.82$ | $-1.25$ | 93.50% | 1.00 | 7,793 |
| | **ZOIB** | | | | | | |
| | $\pi_0$ (Zero-inflation) | 0.3 | $0.30 \pm 0.02$ | 0.00 | 97.00% | 1.00 | 21,272 |
| | $q$ (One-inflation) | 0.01 | $0.01 \pm 0.00$ | 0.00 | 96.00% | 1.00 | 22,047 |
| | $\varphi$ (Precision) | 18 | $18.06 \pm 0.97$ | $+0.06$ | 93.00% | 1.00 | 22,161 |

Table S7: Parameter recovery performance for Power Law dispersal kernel across all mixture scenarios. Results are summarized across 100 independent simulation runs.

| Scenario | Parameter Group | True | Estimate | Bias | Coverage | $\hat{R}$ | ESS |
|---|---|---|---|---|---|---|---|
| *Mixtures (0.5, 0.5)* | **Dispersal & Location** | | | | | | |
| | $a_1$ (Scale 1) | 5.00 | $4.71 \pm 1.20$ | $-0.29$ | 99% | 1.00 | 16,428 |
| | $b_1$ (Exponent 1) | 2.00 | $2.06 \pm 0.31$ | $+0.06$ | 100% | 1.00 | 16,217 |
| | $f_{x1}$ (Source 1 X) | 0.00 | $0.23 \pm 0.10$ | $+0.23$ | 100% | 1.00 | 14,246 |
| | $f_{y1}$ (Source 1 Y) | 0.00 | $-0.11 \pm 0.21$ | $-0.11$ | 96.50% | 1.00 | 28,321 |
| | $f_{z1}$ (Source 1 Z) | 0.60 | $0.84 \pm 0.03$ | $+0.24$ | 18.50% | 1.00 | 15,120 |
| | $a_2$ (Scale 2) | 5.00 | $4.70 \pm 0.95$ | $-0.30$ | 97% | 1.00 | 16,103 |
| | $b_2$ (Exponent 2) | 2.00 | $2.03 \pm 0.23$ | $+0.03$ | 99% | 1.00 | 16,108 |
| | $f_{x2}$ (Source 2 X) | 0.00 | $0.15 \pm 0.06$ | $+0.15$ | 100% | 1.00 | 15,675 |
| | $f_{y2}$ (Source 2 Y) | 50.00 | $50.05 \pm 0.18$ | $+0.05$ | 96.75% | 1.00 | 30,535 |
| | $f_{z2}$ (Source 2 Z) | 0.8 | $0.87 \pm 0.02$ | $+0.07$ | 99.25% | 1.00 | 14,305 |
| | $w$ (Weight) | 0.50 | $0.45 \pm 0.02$ | $-0.05$ | 100% | 1.00 | 17,416 |
| | **ZOIB** | | | | | | |
| | $\pi_0$ (Zero-inflation) | 0.30 | $0.30 \pm 0.02$ | 0.00 | 94% | 1.00 | 41,334 |
| | $q$ (One-inflation) | 0.01 | $0.01 \pm 0.00$ | 0.00 | 96% | 1.00 | 39,305 |
| | $\varphi$ (Precision) | 18.00 | $17.81 \pm 0.98$ | $-0.19$ | 97% | 1.00 | 34,569 |
| *Mixtures (0.6, 0.4)* | **Dispersal & Location** | | | | | | |
| | $a_1$ (Scale 1) | 5.00 | $4.56 \pm 1.18$ | $-0.44$ | 95% | 1.01 | 15,115 |
| | $b_1$ (Exponent 1) | 2.00 | $2.01 \pm 0.30$ | $+0.01$ | 98% | 1.01 | 14,997 |
| | $f_{x1}$ (Source 1 X) | 0.00 | $0.18 \pm 0.09$ | $+0.18$ | 100% | 1.01 | 13,614 |
| | $f_{y1}$ (Source 1 Y) | 0.00 | $-0.08 \pm 0.17$ | $-0.08$ | 96.75% | 1.01 | 28,490 |
| | $f_{z1}$ (Source 1 Z) | 0.60 | $0.86 \pm 0.02$ | $+0.26$ | 6.50% | 1.00 | 13,583 |
| | $a_2$ (Scale 2) | 5.00 | $4.71 \pm 1.02$ | $-0.29$ | 98% | 1.01 | 14,538 |
| | $b_2$ (Exponent 2) | 2.00 | $2.06 \pm 0.26$ | $+0.06$ | 99% | 1.01 | 14,655 |
| | $f_{x2}$ (Source 2 X) | 0.00 | $0.22 \pm 0.10$ | $+0.22$ | 100% | 1.01 | 12,832 |
| | $f_{y2}$ (Source 2 Y) | 50.00 | $50.11 \pm 0.24$ | $+0.11$ | 97.75% | 1.01 | 25,715 |
| | $f_{z2}$ (Source 2 Z) | 0.8 | $0.84 \pm 0.02$ | $+0.04$ | 100% | 1.00 | 13,856 |
| | $w$ (Weight) | 0.60 | $0.52 \pm 0.02$ | $-0.08$ | 98% | 1.01 | 16,867 |
| | **ZOIB** | | | | | | |
| | $\pi_0$ (Zero-inflation) | 0.30 | $0.30 \pm 0.01$ | 0.00 | 97% | 1.01 | 40,054 |
| | $q$ (One-inflation) | 0.01 | $0.01 \pm 0.00$ | 0.00 | 94% | 1.01 | 37,972 |

Continued on next page...

**Table S7 – continued from previous page**

| Scenario | Parameter Group | True | Estimate | Bias | Coverage | $\hat{R}$ | ESS |
|---|---|---|---|---|---|---|---|
| | $\varphi$ (Precision) | 18.00 | $17.74 \pm 1.07$ | $-0.26$ | 95% | 1.00 | 32,662 |
| *Mixtures (0.8, 0.2)* | **Dispersal & Location** | | | | | | |
| | $a_1$ (Scale 1) | 5.00 | $4.50 \pm 1.16$ | $-0.50$ | 88% | 1.00 | 15,624 |
| | $b_1$ (Exponent 1) | 2.00 | $1.98 \pm 0.27$ | $-0.02$ | 95% | 1.00 | 16,069 |
| | $f_{x1}$ (Source 1 X) | 0.00 | $0.12 \pm 0.05$ | $+0.12$ | 100% | 1.00 | 14,168 |
| | $f_{y1}$ (Source 1 Y) | 0.00 | $-0.06 \pm 0.12$ | $-0.06$ | 98% | 1.00 | 30,881 |
| | $f_{z1}$ (Source 1 Z) | 0.6 | $0.87 \pm 0.02$ | $+0.27$ | 0.50% | 1.00 | 11,614 |
| | $a_2$ (Scale 2) | 5.00 | $4.96 \pm 1.00$ | $-0.04$ | 100% | 1.00 | 14,883 |
| | $b_2$ (Exponent 2) | 2.00 | $2.24 \pm 0.27$ | $+0.24$ | 100% | 1.00 | 14,593 |
| | $f_{x2}$ (Source 2 X) | 0.00 | $0.61 \pm 0.28$ | $+0.61$ | 100% | 1.00 | 10,148 |
| | $f_{y2}$ (Source 2 Y) | 50.00 | $50.46 \pm 0.58$ | $+0.46$ | 94.75% | 1.00 | 16,942 |
| | $f_{z2}$ (Source 2 Z) | 0.8 | $0.75 \pm 0.03$ | $-0.05$ | 100% | 1.00 | 15,405 |
| | $w$ (Weight) | 0.80 | $0.65 \pm 0.02$ | $-0.15$ | 34% | 1.00 | 14,807 |
| | **ZOIB** | | | | | | |
| | $\pi_0$ (Zero-inflation) | 0.30 | $0.30 \pm 0.01$ | 0.00 | 96% | 1.00 | 39,423 |
| | $q$ (One-inflation) | 0.01 | $0.01 \pm 0.00$ | 0.00 | 98% | 1.00 | 38,198 |
| | $\varphi$ (Precision) | 18.00 | $17.99 \pm 1.20$ | $-0.01$ | 94% | 1.00 | 33,070 |
| *Mixtures (0.95, 0.05)* | **Dispersal & Location** | | | | | | |
| | $a_1$ (Scale 1) | 5.00 | $4.85 \pm 1.10$ | $-0.15$ | 92% | 1.00 | 14,219 |
| | $b_1$ (Exponent 1) | 2.00 | $2.05 \pm 0.24$ | $+0.05$ | 97% | 1.00 | 14,352 |
| | $f_{x1}$ (Source 1 X) | 0.00 | $0.09 \pm 0.03$ | $+0.09$ | 100% | 1.00 | 15,193 |
| | $f_{y1}$ (Source 1 Y) | 0.00 | $-0.03 \pm 0.09$ | $-0.03$ | 98.25% | 1.00 | 31,203 |
| | $f_{z1}$ (Source 1 Z) | 0.6 | $0.88 \pm 0.02$ | $+0.28$ | 0% | 1.00 | 10,277 |
| | $a_2$ (Scale 2) | 5.00 | $4.87 \pm 0.49$ | $-0.13$ | 100% | 1.00 | 16,533 |
| | $b_2$ (Exponent 2) | 2.00 | $2.40 \pm 0.29$ | $+0.40$ | 100% | 1.00 | 17,759 |
| | $f_{x2}$ (Source 2 X) | 0.00 | $2.47 \pm 0.80$ | $+2.47$ | 100% | 1.00 | 14,527 |
| | $f_{y2}$ (Source 2 Y) | 50.00 | $52.61 \pm 1.12$ | $+2.61$ | 96% | 1.00 | 11,302 |
| | $f_{z2}$ (Source 2 Z) | 0.8 | $0.62 \pm 0.04$ | $-0.18$ | 100% | 1.00 | 16,112 |
| | $w$ (Weight) | 0.95 | $0.74 \pm 0.03$ | $-0.21$ | 0% | 1.00 | 13,239 |
| | **ZOIB** | | | | | | |
| | $\pi_0$ (Zero-inflation) | 0.30 | $0.30 \pm 0.01$ | 0.00 | 98% | 1.00 | 40,413 |
| | $q$ (One-inflation) | 0.01 | $0.01 \pm 0.00$ | 0.00 | 98% | 1.00 | 36,832 |

Continued on next page...

**Table S7 – continued from previous page**

| Scenario | Parameter Group | True | Estimate | Bias | Coverage | $\hat{R}$ | ESS |
|---|---|---|---|---|---|---|---|
| | $\varphi$ (Precision) | 18.00 | $17.94 \pm 1.13$ | $-0.06$ | 96% | 1.00 | 33,735 |

Table S8: Parameter recovery performance across gradients of spatial complexity and sample sizes ($N = 50, 100, 250, 500$). Values represent mean ± SD across 100 independent simulations.

| **Scenario** | **Parameter Group** | **True** | **Estimation** | **Bias** | **Coverage** | $\hat{R}$ | **ESS** |
|---|---|---|---|---|---|---|---|
| *Single-Focus_N50* | **Dispersal & Location** | | | | | | |
| | $a_1$ (Scale) | 5.00 | $4.18 \pm 0.99$ | $-0.82$ | 90.00% | 1.00 | 10,444 |
| | $b_1$ (Exponent) | 2.00 | $1.96 \pm 0.18$ | $-0.04$ | 97.00% | 1.00 | 11,840 |
| | $f_{x1}$ (Source X) | 0.00 | $0.29 \pm 0.17$ | $+0.29$ | 100.00% | 1.00 | 8,090 |
| | $f_{y1}$ (Source Y) | 0.00 | $-0.07 \pm 0.24$ | $-0.07$ | 98.00% | 1.00 | 17,396 |
| | **ZOIB** | | | | | | |
| | $\pi_0$ (Zero-inflation) | 0.30 | $0.30 \pm 0.04$ | 0.00 | 94.00% | 1.00 | 23,872 |
| | $q$ (One-inflation) | 0.01 | $0.02 \pm 0.01$ | $+0.01$ | 98.00% | 1.00 | 21,105 |
| | $\varphi$ (Precision) | 18.00 | $17.98 \pm 3.02$ | $-0.02$ | 95.00% | 1.00 | 16,117 |
| *Single-Focus_N100* | **Dispersal & Location** | | | | | | |
| | $a_1$ (Scale) | 5.00 | $3.90 \pm 0.72$ | $-1.10$ | 77.00% | 1.00 | 9,116 |
| | $b_1$ (Exponent) | 2.00 | $1.90 \pm 0.15$ | $-0.10$ | 90.00% | 1.00 | 11,005 |
| | $f_{x1}$ (Source X) | 0.00 | $0.20 \pm 0.08$ | $+0.20$ | 100.00% | 1.00 | 6,188 |
| | $f_{y1}$ (Source Y) | 0.00 | $-0.04 \pm 0.18$ | $-0.04$ | 96.00% | 1.00 | 19,956 |
| | **ZOIB** | | | | | | |
| | $\pi_0$ (Zero-inflation) | 0.30 | $0.30 \pm 0.03$ | 0.00 | 93.00% | 1.00 | 23,268 |
| | $q$ (One-inflation) | 0.01 | $0.01 \pm 0.01$ | 0.00 | 97.00% | 1.00 | 23,592 |
| | $\varphi$ (Precision) | 18.00 | $18.02 \pm 1.96$ | $+0.02$ | 93.00% | 1.00 | 19,637 |
| *Single-Focus_N250* | **Dispersal & Location** | | | | | | |
| | $a_1$ (Scale) | 5.00 | $4.06 \pm 0.60$ | $-0.94$ | 76.00% | 1.00 | 7,310 |
| | $b_1$ (Exponent) | 2.00 | $1.90 \pm 0.11$ | $-0.10$ | 85.00% | 1.00 | 9,466 |
| | $f_{x1}$ (Source X) | 0.00 | $0.11 \pm 0.05$ | $+0.11$ | 100.00% | 1.00 | 5,425 |
| | $f_{y1}$ (Source Y) | 0.00 | $-0.02 \pm 0.09$ | $-0.02$ | 98.00% | 1.00 | 21,272 |
| | **ZOIB** | | | | | | |
| | $\pi_0$ (Zero-inflation) | 0.30 | $0.30 \pm 0.01$ | 0.00 | 96.00% | 1.00 | 23,690 |
| | $q$ (One-inflation) | 0.01 | $0.01 \pm 0.00$ | 0.00 | 93.00% | 1.00 | 24,566 |
| | $\varphi$ (Precision) | 18.00 | $18.13 \pm 1.32$ | $+0.13$ | 95.00% | 1.00 | 21,956 |
| *Single-Focus_N500* | **Dispersal & Location** | | | | | | |
| | $a_1$ (Scale) | 5.00 | $4.25 \pm 0.48$ | $-0.75$ | 72.00% | 1.00 | 7,274 |

Continued on next page...

Table S8 – continued from previous page

| Scenario | Parameter Group | True | Estimation | Bias | Coverage | $\hat{R}$ | ESS |
|---|---|---|---|---|---|---|---|
| | $b_1$ (Exponent) | 2.00 | $1.92 \pm 0.08$ | $-0.08$ | 88.00% | 1.00 | 8,888 |
| | $f_{x1}$ (Source X) | 0.00 | $0.06 \pm 0.05$ | $+0.06$ | 100.00% | 1.00 | 6,274 |
| | $f_{y1}$ (Source Y) | 0.00 | $-0.01 \pm 0.07$ | $-0.01$ | 96.25% | 1.00 | 21,702 |
| | **ZOIB** | | | | | | |
| | $\pi_0$ (Zero-inflation) | 0.30 | $0.30 \pm 0.01$ | 0.00 | 97.00% | 1.00 | 24,123 |
| | $q$ (One-inflation) | 0.01 | $0.01 \pm 0.00$ | 0.00 | 99.00% | 1.00 | 23,724 |
| | $\varphi$ (Precision) | 18.00 | $18.14 \pm 0.87$ | $+0.14$ | 97.00% | 1.00 | 22,245 |
| *Two-Foci_N50* | **Dispersal & Location** | | | | | | |
| | $a_1$ (Scale 1) | 5.00 | $6.76 \pm 0.96$ | $+1.76$ | 100.00% | 1.00 | 11,941 |
| | $b_1$ (Exponent 1) | 2.00 | $2.39 \pm 0.27$ | $+0.39$ | 100.00% | 1.00 | 13,651 |
| | $a_2$ (Scale 2) | 5.00 | $6.51 \pm 0.93$ | $+1.51$ | 100.00% | 1.00 | 11,919 |
| | $b_2$ (Exponent 2) | 2.00 | $2.38 \pm 0.23$ | $+0.38$ | 100.00% | 1.00 | 13,282 |
| | $f_{x1}$ (Source 1 X) | 0.00 | $0.64 \pm 0.29$ | $+0.64$ | 100.00% | 1.00 | 14,849 |
| | $f_{x2}$ (Source 2 X) | 0.00 | $0.70 \pm 0.30$ | $+0.70$ | 100.00% | 1.00 | 15,031 |
| | $f_{y1}$ (Source 1 Y) | 0.00 | $-0.23 \pm 0.99$ | $-0.23$ | 98.75% | 1.01 | 16,967 |
| | $f_{y2}$ (Source 2 Y) | 50.00 | $50.20 \pm 0.98$ | $+0.20$ | 98.75% | 1.01 | 16,388 |
| | $w$ (Weight) | 0.60 | $0.52 \pm 0.03$ | $-0.08$ | 94.00% | 1.00 | 16,680 |
| | **ZOIB** | | | | | | |
| | $\pi_0$ (Zero-inflation) | 0.30 | $0.31 \pm 0.04$ | $+0.01$ | 92.00% | 1.00 | 28,181 |
| | $q$ (One-inflation) | 0.01 | $0.02 \pm 0.01$ | $+0.01$ | 97.00% | 1.00 | 24,552 |
| | $\varphi$ (Precision) | 18.00 | $16.82 \pm 2.63$ | $-1.18$ | 90.00% | 1.00 | 13,566 |
| *Two-Foci_N100* | **Dispersal & Location** | | | | | | |
| | $a_1$ (Scale 1) | 5.00 | $5.49 \pm 1.16$ | $+0.49$ | 100.00% | 1.00 | 11,592 |
| | $b_1$ (Exponent 1) | 2.00 | $2.22 \pm 0.28$ | $+0.22$ | 100.00% | 1.00 | 12,225 |
| | $a_2$ (Scale 2) | 5.00 | $5.73 \pm 1.06$ | $+0.73$ | 100.00% | 1.00 | 11,502 |
| | $b_2$ (Exponent 2) | 2.00 | $2.24 \pm 0.25$ | $+0.24$ | 100.00% | 1.00 | 12,167 |
| | $f_{x1}$ (Source 1 X) | 0.00 | $0.34 \pm 0.13$ | $+0.34$ | 100.00% | 1.00 | 13,224 |
| | $f_{x2}$ (Source 2 X) | 0.00 | $0.45 \pm 0.18$ | $+0.45$ | 100.00% | 1.00 | 12,701 |
| | $f_{y1}$ (Source 1 Y) | 0.00 | $-0.17 \pm 0.33$ | $-0.17$ | 97.00% | 1.00 | 18,570 |
| | $f_{y2}$ (Source 2 Y) | 50.00 | $50.29 \pm 0.33$ | $+0.29$ | 98.50% | 1.00 | 17,743 |
| | $w$ (Weight) | 0.60 | $0.52 \pm 0.02$ | $-0.08$ | 98.00% | 1.00 | 15,397 |
| | **ZOIB** | | | | | | |

Continued on next page...

**Table S8 – continued from previous page**

| Scenario | Parameter Group | True | Estimation | Bias | Coverage | $\hat{R}$ | ESS |
|---|---|---|---|---|---|---|---|
| | $\pi_0$ (Zero-inflation) | 0.30 | $0.30 \pm 0.02$ | 0.00 | 99.00% | 1.00 | 27,301 |
| | $q$ (One-inflation) | 0.01 | $0.01 \pm 0.01$ | 0.00 | 93.00% | 1.00 | 23,738 |
| | $\varphi$ (Precision) | 18.00 | $17.42 \pm 1.60$ | $-0.58$ | 93.00% | 1.00 | 17,935 |
| *Two-Foci_N250* | **Dispersal & Location** | | | | | | |
| | $a_1$ (Scale 1) | 5.00 | $4.56 \pm 1.18$ | $-0.44$ | 95.00% | 1.01 | 15,115 |
| | $b_1$ (Exponent 1) | 2.00 | $2.01 \pm 0.30$ | $+0.01$ | 98.00% | 1.01 | 14,997 |
| | $a_2$ (Scale 2) | 5.00 | $4.71 \pm 1.02$ | $-0.29$ | 98.00% | 1.01 | 14,538 |
| | $b_2$ (Exponent 2) | 2.00 | $2.06 \pm 0.26$ | $+0.06$ | 99.00% | 1.01 | 14,655 |
| | $f_{x1}$ (Source 1 X) | 0.00 | $0.18 \pm 0.09$ | $+0.18$ | 100.00% | 1.01 | 13,614 |
| | $f_{x2}$ (Source 2 X) | 0.00 | $0.22 \pm 0.10$ | $+0.22$ | 100.00% | 1.01 | 12,832 |
| | $f_{y1}$ (Source 1 Y) | 0.00 | $-0.08 \pm 0.17$ | $-0.08$ | 96.75% | 1.01 | 28,490 |
| | $f_{y2}$ (Source 2 Y) | 50.00 | $50.11 \pm 0.24$ | $+0.11$ | 97.75% | 1.01 | 25,715 |
| | $w$ (Weight) | 0.60 | $0.52 \pm 0.02$ | $-0.08$ | 98.00% | 1.01 | 16,867 |
| | **ZOIB** | | | | | | |
| | $\pi_0$ (Zero-inflation) | 0.30 | $0.30 \pm 0.01$ | 0.00 | 97.00% | 1.01 | 40,054 |
| | $q$ (One-inflation) | 0.01 | $0.01 \pm 0.00$ | 0.00 | 94.00% | 1.01 | 37,972 |
| | $\varphi$ (Precision) | 18.00 | $17.74 \pm 1.07$ | $-0.26$ | 95.00% | 1.00 | 32,662 |
| *Two-Foci_N500* | **Dispersal & Location** | | | | | | |
| | $a_1$ (Scale 1) | 5.00 | $4.33 \pm 0.99$ | $-0.67$ | 93.00% | 1.00 | 12,280 |
| | $b_1$ (Exponent 1) | 2.00 | $1.96 \pm 0.25$ | $-0.04$ | 97.00% | 1.00 | 12,374 |
| | $a_2$ (Scale 2) | 5.00 | $4.28 \pm 1.00$ | $-0.72$ | 91.00% | 1.00 | 12,010 |
| | $b_2$ (Exponent 2) | 2.00 | $1.96 \pm 0.26$ | $-0.04$ | 97.00% | 1.00 | 12,204 |
| | $f_{x1}$ (Source 1 X) | 0.00 | $0.11 \pm 0.04$ | $+0.11$ | 100.00% | 1.00 | 11,556 |
| | $f_{x2}$ (Source 2 X) | 0.00 | $0.13 \pm 0.06$ | $+0.13$ | 100.00% | 1.00 | 10,976 |
| | $f_{y1}$ (Source 1 Y) | 0.00 | $-0.04 \pm 0.13$ | $-0.04$ | 96.50% | 1.00 | 29,287 |
| | $f_{y2}$ (Source 2 Y) | 50.00 | $50.04 \pm 0.14$ | $+0.04$ | 96.25% | 1.00 | 27,820 |
| | $w$ (Weight) | 0.60 | $0.52 \pm 0.02$ | $-0.08$ | 99.00% | 1.00 | 12,111 |
| | **ZOIB** | | | | | | |
| | $\pi_0$ (Zero-inflation) | 0.30 | $0.30 \pm 0.01$ | 0.00 | 98.00% | 1.00 | 41,873 |
| | $q$ (One-inflation) | 0.01 | $0.01 \pm 0.00$ | 0.00 | 97.00% | 1.00 | 37,943 |
| | $\varphi$ (Precision) | 18.00 | $17.95 \pm 0.74$ | $-0.05$ | 96.00% | 1.00 | 35,644 |

Table S9: Predictive Performance and LOO-CV Results of Single-Focus for Cucurbit Downy Mildew Field Experiment under Scenarios of **Unknown Focus Location** (Joint Estimation).

| | Single-focus (inoculated once) | | | Single-focus (inoculated twice) | | |
|---|---|---|---|---|---|---|
| **Metric** | Exponential | Gaussian | Power Law | Exponential | Gaussian | Power Law |
| overall $R^2$ | 0.481 | 0.012 | **0.649** | 0.766 | 0.739 | **0.835** |
| overall RMSE | 0.089 | 0.123 | **0.073** | 0.112 | 0.118 | **0.094** |
| overall MAE | 0.068 | 0.090 | **0.057** | 0.088 | 0.093 | **0.073** |
| obs_zeros | 0.163 | 0.163 | 0.163 | 0.017 | 0.017 | 0.017 |
| pred_zeros | 0.182 | 0.180 | 0.180 | 0.028 | 0.028 | 0.028 |
| obs_ones | 0.000 | 0.000 | 0.000 | 0.000 | 0.000 | 0.000 |
| pred_ones | 0.017 | 0.017 | 0.017 | 0.014 | 0.014 | 0.014 |
| continuous $R^2$ | 0.471 | -0.006 | **0.652** | 0.767 | 0.740 | **0.835** |
| continuous RMSE | 0.096 | 0.133 | **0.078** | 0.112 | 0.118 | **0.094** |
| n_obs | 43 | 43 | 43 | 59 | 59 | 59 |
| elpd_loo | 42.516 | 35.918 | **49.572** | 66.861 | 65.410 | **74.353** |
| p_loo | 3.269 | 4.327 | **2.904** | 6.071 | 5.841 | **4.983** |
| rank_loo | 2 | 3 | **1** | 2 | 3 | **1** |
| se_d_loo | 1.167 | 2.647 | 0.000 | 2.029 | 3.554 | 0.000 |
| warning | False | False | False | False | False | False |
| pareto_k_bad | 0 | 0 | 0 | 0 | 0 | 0 |

*Note:* Bold values indicate the best performance in each experimental condition.

Table S10: Predictive Performance and LOO-CV Results of Two-foci Models for Cucurbit Downy Mildew Field Experimental Data under Scenario of **Unknown Focus Location**.

| | Two-foci (inoculated once) | | | Two-foci (inoculated twice) | | |
|---|---|---|---|---|---|---|
| **Metric** | Exponential | Gaussian | Power Law | Exponential | Gaussian | Power Law |
| overall $R^2$ | 0.526 | 0.439 | **0.596** | 0.507 | 0.430 | **0.522** |
| overall RMSE | 0.105 | 0.114 | **0.097** | 0.142 | 0.153 | **0.140** |
| overall MAE | 0.078 | 0.087 | **0.074** | 0.102 | 0.116 | **0.100** |
| obs_zeros | 0.034 | 0.034 | 0.034 | 0.000 | 0.000 | 0.000 |
| pred_zeros | 0.043 | 0.042 | 0.042 | 0.016 | 0.014 | 0.014 |
| obs_ones | 0.000 | 0.000 | 0.000 | 0.000 | 0.000 | 0.000 |
| pred_ones | 0.014 | 0.014 | 0.014 | 0.014 | 0.014 | 0.014 |
| continuous $R^2$ | 0.525 | 0.443 | **0.596** | 0.507 | 0.430 | **0.522** |
| continuous RMSE | 0.105 | 0.114 | **0.097** | 0.142 | 0.153 | **0.140** |
| n_obs | 58 | 58 | 58 | 59 | 59 | 59 |
| elpd_loo | 55.297 | 49.991 | **57.628** | 50.830 | 46.442 | **51.301** |
| p_loo | **6.333** | 7.314 | 6.679 | **6.595** | 7.537 | 6.760 |
| rank_loo | 2 | 3 | **1** | 2 | 3 | **1** |
| se_d_loo | 0.874 | 1.153 | 0.000 | 1.222 | 1.973 | 0.000 |
| warning | False | False | False | False | False | False |
| pareto_k_bad | 0 | 0 | 0 | 0 | 0 | 0 |

*Note:* Bold values indicate the best performance in each category.

Table S11: **Bayesian model selection summary for spatial analyses of the 1854 Soho cholera epidemic data.** Models are compared globally (overall) and within their structural class (within-group) using leave-one-out cross-validation (LOO-CV). *Abbreviations:* ELPD, Expected Log Pointwise Predictive Density; $p_{loo}$, effective number of parameters; **SE**, standard error of the difference in ELPD; Pareto-$k$ bad, number of observations with Pareto-$k > 0.7$.

| | **Model Fit Statistics** | | **Overall** | | **Within-Group Comparison** | | | **Forensic** | **Diagnostics** | |
|---|---|---|---|---|---|---|---|---|---|---|
| **Model Specification** | **ELPD** | $p_{loo}$ | **Rank** | **SE** | **Rank** | **SE** | **Equivalent** ‡ | **Localization Error (m)** | **Warning** | **Pareto-$k$ bad** |
| ***Single-Focus Models*** *($H_1$: Blind Discovery)* | | | | | | | | | | |
| Gaussian | −346.73 | 8.19 | 1 | 0.00 | 1 | 0.00 | – | 41.61 | False | 0 |
| Exponential | −350.91 | 8.56 | 2 | 2.92 | 2 | 2.92 | True | 34.37 | False | 0 |
| Power Law | −353.82 | 8.98 | 3 | 3.78 | 3 | 3.78 | True | **32.65** | False | 0 |
| ***Two-Foci Models*** *($H_2$: Parsimony Stress Test)* | | | | | | | | | | |
| Gaussian* | −354.92 | 6.69 | 4 | 3.59 | 1 | 0.00 | – | 1.93; 0.12 | True | 1 |
| Exponential | −356.38 | 6.79 | 5 | 4.61 | 2 | 3.06 | True | 1.77; 0.09 | False | 0 |
| Power Law | −360.48 | 7.39 | 6 | 5.53 | 3 | 4.62 | True | **1.42; 0.06** | False | 0 |
| *Null Model* | −403.86 | 5.59 | 7 | 8.26 | – | – | – | – | False | 0 |

* **Warning:** Model generated a Pareto-$k$ warning ($k > 0.7$), indicating potential unreliability due to influential observations.

‡ **Predictive Equivalence:** Defines where the difference in ELPD is less than twice the standard error of the difference ($|\Delta\text{ELPD}| < 2 \times \text{SE}$).

**Two-Foci Localization:** Values represent the Euclidean distance to the Broad Street pump (529396.54, 181025.06) and the Little Marlborough Street pump (529192.54, 181079.39), respectively.

Table S12: **Sensitivity of forensic localization to spatial prior precision.** Single-Focus models were evaluated across varying spatial prior widths ($\sigma \in [5, 250]$) to quantify the interaction between prior knowledge and data-driven inference. *Abbreviations:* $p_{loo}$, effective number of parameters; $SE_{\Delta}$, standard error of the difference in ELPD; Pred. Eq., Predictive Equivalent (statistically indistinguishable from the top-ranked model where $|\Delta ELPD| < 2 \times SE$); Pareto-$k$ bad, number of observations with Pareto-$k > 0.7$.

| $\sigma$ (m) | Model Kernel | Rank (LOO) | ELPD | $p_{loo}$ | SE | ΔELPD | Warning | Pareto-$k$ bad | Pred. Eq. | $f_{x\_est}$ (m) | $f_{y\_est}$ (m) | Localization Error (m) |
|---|---|---|---|---|---|---|---|---|---|---|---|---|
| | Gaussian | 1 | -351.6 | 7.1 | 0.0 | 0.0 | False | 0 | - | 529404.54 | 181030.39 | 9.61 |
| 5 | Exponential | 2 | -353.9 | 7.3 | 2.9 | -2.4 | False | 0 | True | 529403.35 | 181029.29 | 8.02 |
| | Power Law | 3 | -356.2 | 7.4 | 3.8 | -4.6 | False | 0 | True | 529403.07 | 181028.78 | **7.51** |
| | Gaussian | 1 | -347.9 | 7.4 | 0.0 | 0.0 | False | 0 | - | 529415.32 | 181038.11 | 22.87 |
| 10 | Exponential | 2 | -351.6 | 7.8 | 2.9 | -3.7 | False | 0 | True | 529413.26 | 181034.74 | 19.32 |
| | Power Law | 3 | -354.4 | 8.2 | 3.8 | -6.5 | False | 0 | True | 529412.68 | 181033.55 | **18.23** |
| | Gaussian | 1 | -346.6 | 8.0 | 0.0 | 0.0 | False | 0 | - | 529426.14 | 181046.58 | 36.59 |
| 25 | Exponential | 2 | -350.8 | 8.4 | 2.9 | -4.2 | False | 0 | True | 529423.37 | 181039.66 | 30.54 |
| | Power Law | 3 | -353.6 | 8.7 | 3.8 | -7.1 | False | 0 | True | 529422.72 | 181037.71 | **29.07** |
| | Gaussian | 1 | -346.6 | 8.1 | 0.0 | 0.0 | False | 0 | - | 529428.98 | 181048.91 | 40.26 |
| 50 | Exponential | 2 | -350.9 | 8.6 | 2.9 | -4.3 | False | 0 | True | 529425.91 | 181040.70 | 33.28 |
| | Power Law | 3 | -353.7 | 8.9 | 3.8 | -7.1 | False | 0 | True | 529425.24 | 181038.64 | **31.75** |
| | Gaussian | 1 | -346.7 | 8.1 | 0.0 | 0.0 | False | 0 | - | 529429.92 | 181049.51 | 41.37 |
| 100 | Exponential | 2 | -351.0 | 8.6 | 2.9 | -4.3 | False | 0 | True | 529426.70 | 181041.01 | 34.11 |
| | Power Law | 3 | -353.8 | 9.0 | 3.8 | -7.1 | True | 1 | True | 529426.01 | 181038.84 | **32.53** |
| | Gaussian | 1 | -346.7 | 8.2 | 0.0 | 0.0 | False | 0 | - | 529430.07 | 181049.83 | 41.69 |
| 250 | Exponential | 2 | -351.0 | 8.7 | 2.9 | -4.3 | False | 0 | True | 529426.96 | 181041.15 | 34.41 |
| | Power Law | 3 | -353.8 | 8.9 | 3.8 | -7.1 | False | 0 | True | 529426.18 | 181038.96 | **32.74** |

**Note:** Localization error is the Euclidean distance between the estimated focus and the true source, which refers to the Broad Street Pump coordinates ($x = 529396.54, y = 181025.06$).

Table S13: **Predictive performance metrics for spatial models of the 1854 Soho cholera epidemic.** Comparison of candidate models (Null, Single-Focus, Two-Foci) based on overall goodness-of-fit ($R^2$, RMSE) and capability to recover the proportions of zero- and one-inflation. *Abbreviations:* RMSE, Root Mean Square Error; MAE, Mean Absolute Error; Continuous RMSE, RMSE of the continuous Beta component.

| | **Overall Fit** | | | **Zeros** ($P(y=0)$) | | **Ones** ($P(y=1)$) | | **Intensity** |
|---|---|---|---|---|---|---|---|---|
| **Model Specification** | $R^2$ | **RMSE** | **MAE** | **Obs** | **Pred** | **Obs** | **Pred** | **Cont. RMSE** |
| *Null Model* | $< 0.001$ | 0.0589 | 0.0345 | 0.801 | 0.800 | 0.0005 | 0.0010 | 0.1136 |
| ***Single-Focus Models*** | | | | | | | | |
| Exponential | 0.093 | 0.0561 | 0.0279 | 0.801 | 0.822 | 0.0005 | 0.0009 | 0.1107 |
| Gaussian | 0.089 | 0.0562 | 0.0273 | 0.801 | 0.826 | 0.0005 | 0.0009 | 0.1109 |
| Power Law | 0.093 | 0.0561 | 0.0283 | 0.801 | 0.820 | 0.0005 | 0.0009 | 0.1107 |
| ***Two-Foci Models*** | | | | | | | | |
| Exponential | 0.093 | 0.0561 | 0.0282 | 0.801 | 0.821 | 0.0005 | 0.0009 | 0.1108 |
| Gaussian | 0.087 | 0.0563 | 0.0279 | 0.801 | 0.823 | 0.0005 | 0.0009 | 0.1110 |
| Power Law | 0.086 | 0.0563 | 0.0292 | 0.801 | 0.816 | 0.0005 | 0.0010 | 0.1110 |

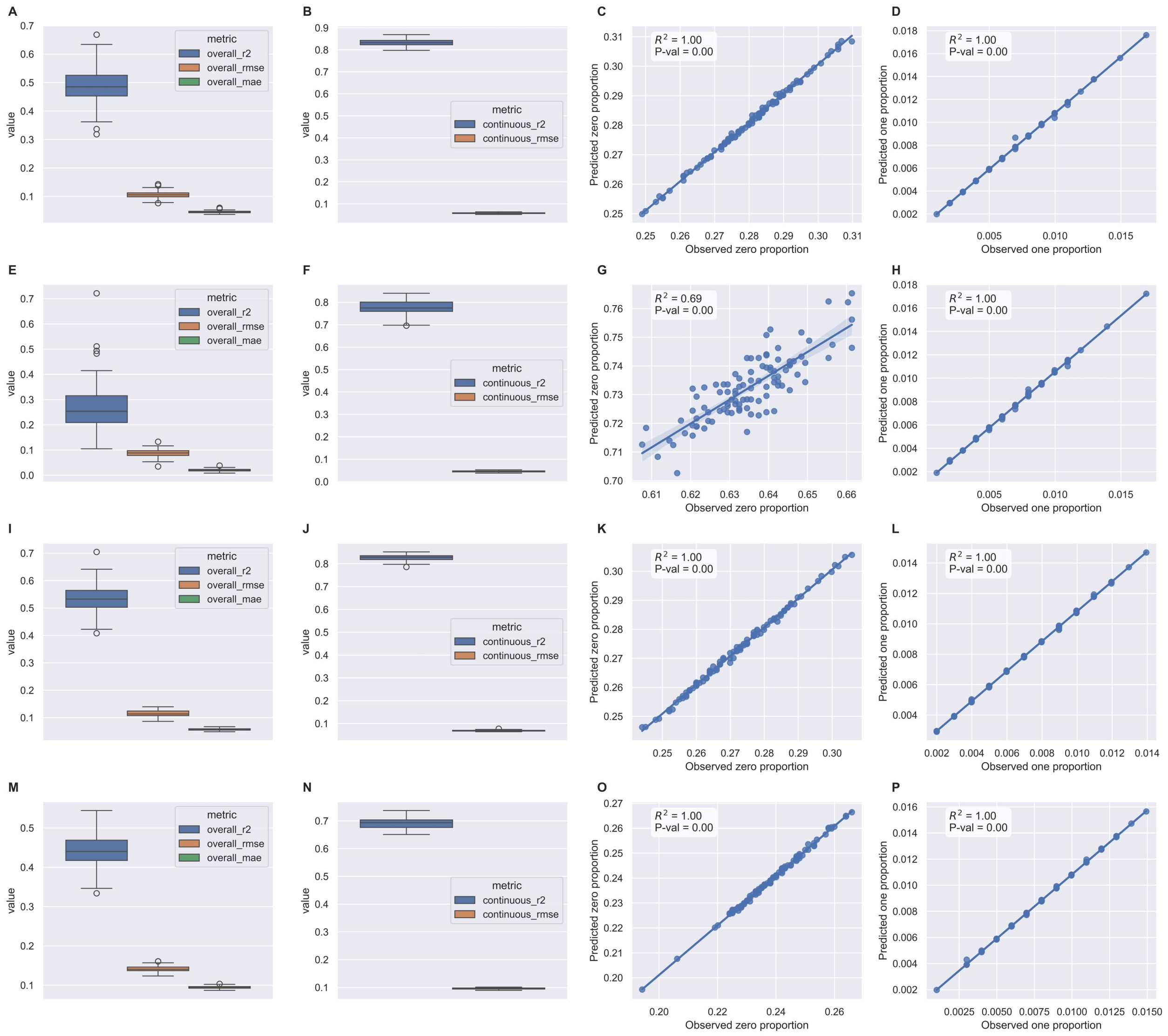


Figure S1: **Posterior predictive checks (PPC) and predictive performance across four epidemiological scenarios.** Rows correspond to the scenarios defined in Table S1: (A-D) Base, (E-H) Narrow & Steep, (I-L) Wide & Steep, and (M-P) Wide & Shallow. Metrics include aggregate $R^2$, RMSE, and MAE (A, E, I, M), alongside specific $R^2$ and RMSE for the continuous Beta component (B, F, J, N); Scatter plots evaluate model calibration by comparing observed versus predicted proportions of zero-inflation (C, G, K, O) and one-inflation (D, H, L, P). The **Narrow & Steep** scenario exhibited the lowest aggregate $R^2$ ($0.27 \pm 0.09$, panel E) and the highest spatial sparsity, with zero proportion exceeding 60%. The reduced predictive fit in this scenario ($R^2 = 0.69$, panel G) indicates that extreme localization increases uncertainty at the infection cluster boundaries. In contrast, other scenarios maintained higher aggregate $R^2$ (0.44 to 0.53, panels A, I, and M) and near-perfect zero-prediction ($R^2 = 1.00$) within a 0.2 - 0.3 proportion range (C, K, O).

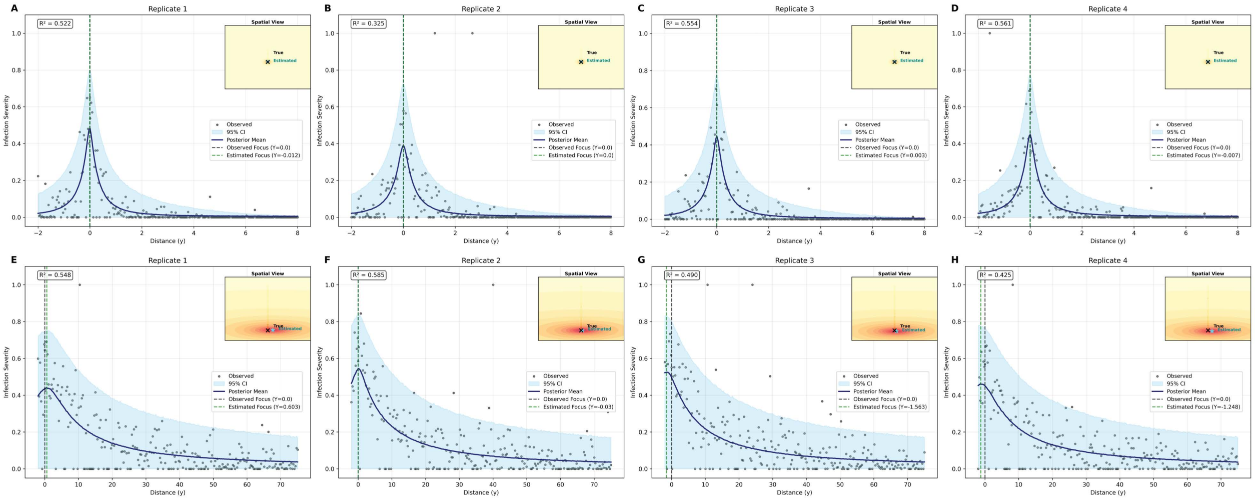


Figure S2: **Spatial validation of posterior predictive distribution for Narrow & Steep ($y = 8$) and Wide & shallow ($y = 75$) scenarios at optimized sampling scales.** Panels (A-D) and (E-H) display the observed infection frequency distributions alongside the posterior predictive mean (navy line) and 95% credible interval (shaded sky-blue region) for the Narrow & Steep and Wide & Shallow scenarios, respectively. The vertical dashed lines indicate the estimated infection focus at $y = 0$. Corresponding 2D inserts visualize the framework's capacity for joint estimation, successfully localizing unknown infection sources (stars) relative to the generative true foci (crosses) using weakly informative priors.

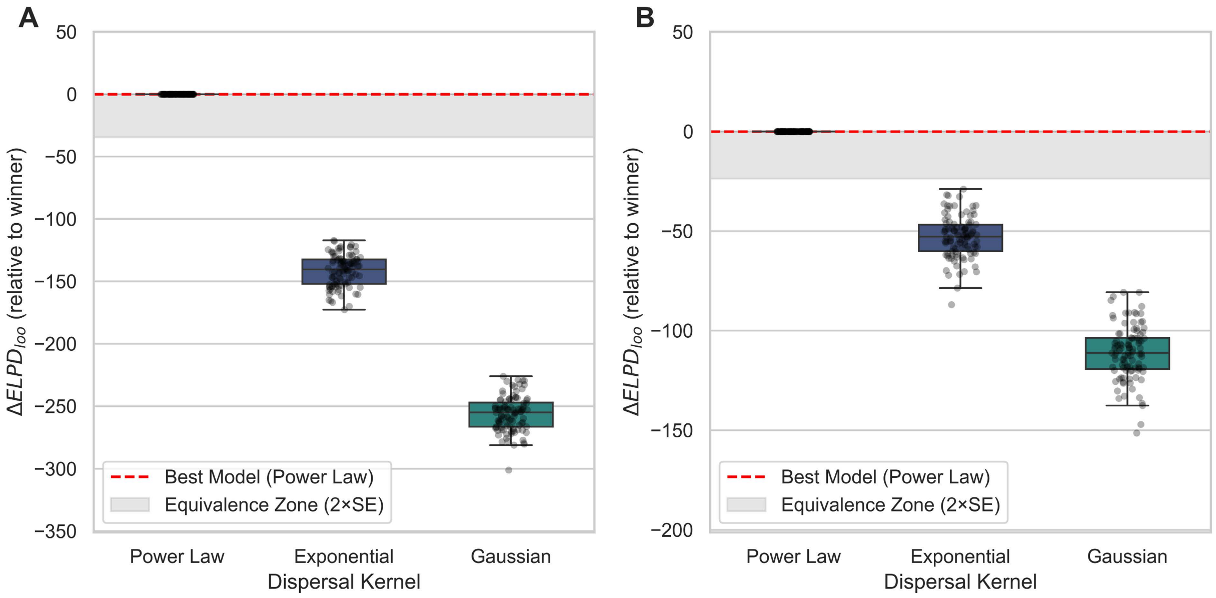


Figure S3: **Model selection sensitivity and structural identifiability for Narrow & Steep ($y = 8$) and Wide & shallow ($y = 75$) scenarios at optimized sampling scales.** Panels show the distribution of $\Delta ELPD_{loo}$ relative to the best performing model for (A) the Narrow & Steep scenario at the $y = 8$ scale and (B) the Wide & Shallow scenario at $y = 75$ scale. In both optimized configurations, the framework correctly identified the generative Power Law kernel in 100% simulations.

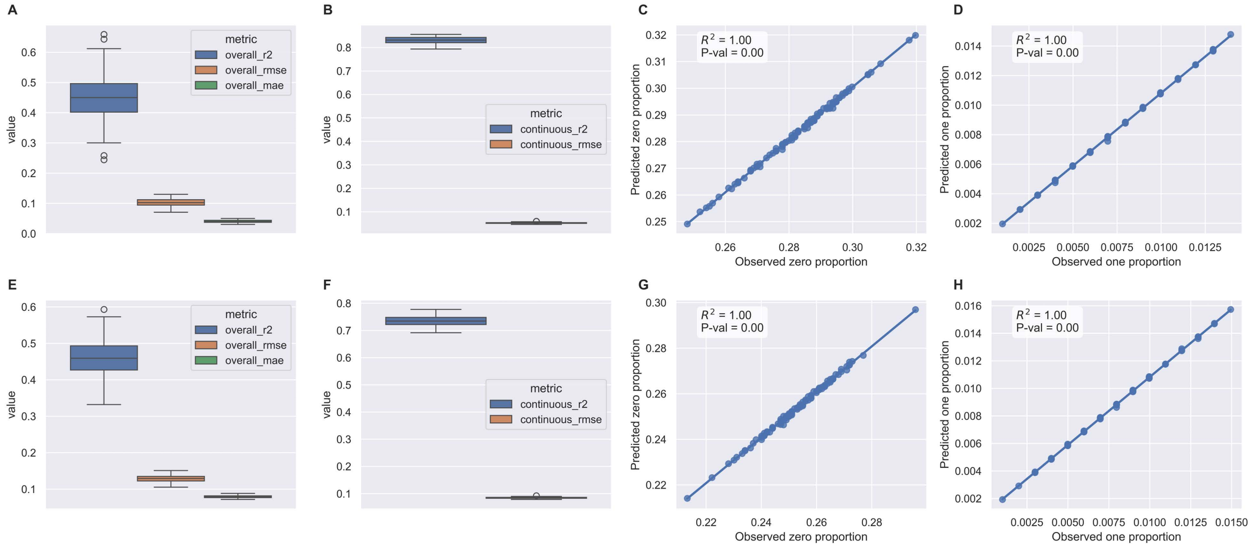


Figure S4: **Posterior predictive checks (PPC) and predictive performance for Narrow & Steep ($y = 8$) and Wide & shallow ($y = 75$) scenarios at optimized sampling scales.** Rows 1 and 2 correspond to the Narrow & Steep and Wide & shallow scenarios, respectively. Metrics evaluated across 100 simulations include aggregated $R^2$, RMSE, and MAE (A, E), alongside specific $R^2$ and RMSE for the continuous Beta component (B, F); Scatter plots demonstrate near-perfect model calibration ($R^2 = 1.00$) for both zero-inflation (C, G) and one-inflation (D, H) proportions.

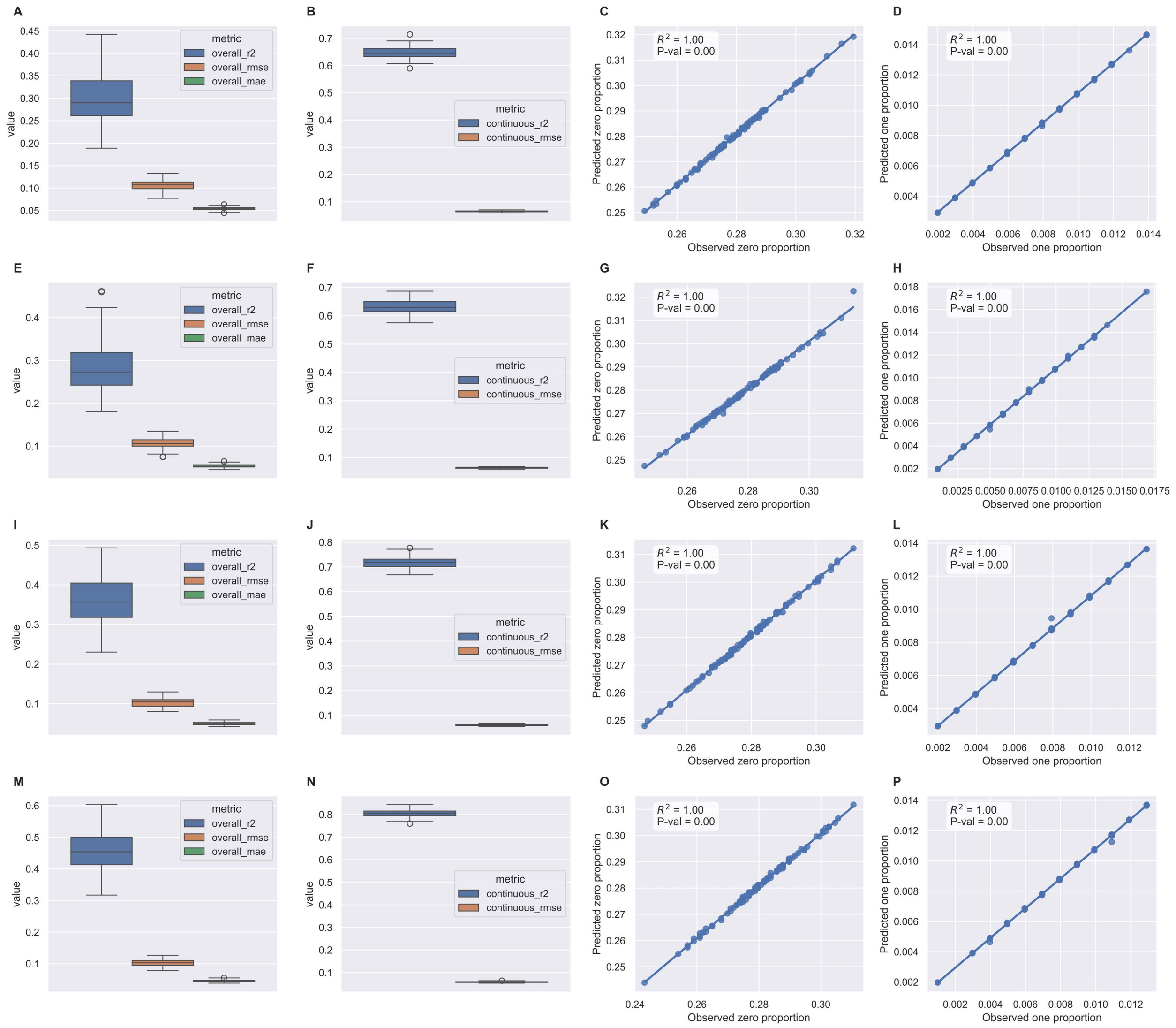


Figure S5: **Posterior predictive checks (PPC) and predictive performance of Hi-BASIL across diverse foci mixture weights.** Rows correspond to foci mixture weight configurations: (A-D) (0.5, 0.5), (E-H) (0.6, 0.4), (I-L) (0.8, 0.2), and (M-P) (0.95, 0.05). Metrics include aggregate $R^2$, RMSE, and MAE (A, E, I, M), alongside specific $R^2$ and RMSE for the continuous Beta component (B, F, J, N); Scatter plots evaluate model calibration by comparing observed versus predicted proportions of zero-inflation (C, G, K, O) and one-inflation (D, H, L, P), both predictive accuracy remain high and stable across all configurations ($R^2 = 1.0$).

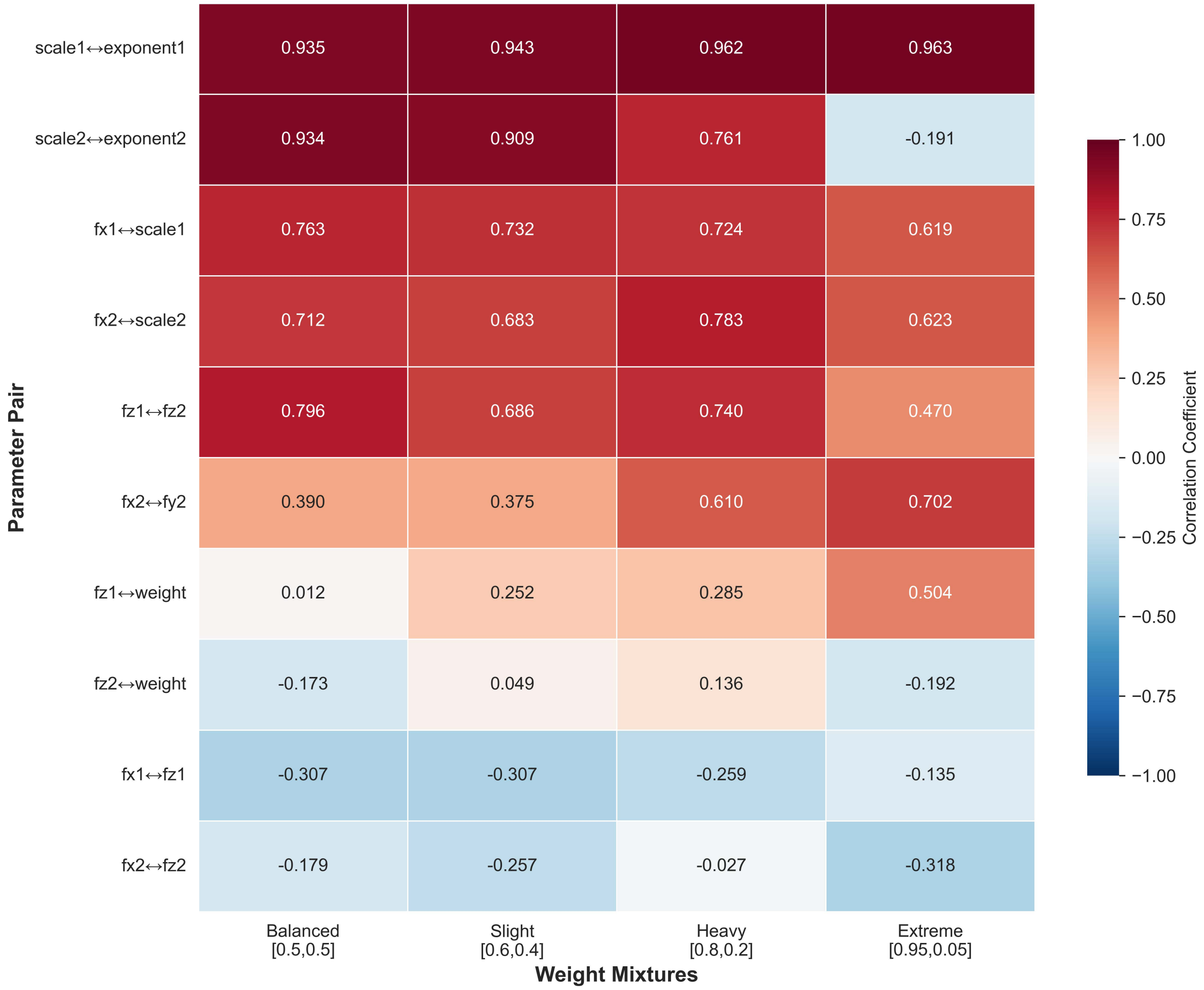


Figure S6: **Heatmap of Pearson correlation coefficients between parameter estimation biases across weight mixture scenarios.** The plot illustrates the evolution of confounding between parameter pairs evolves across four weight mixtures ranging from balanced ($[0.5, 0.5]$) to extremely skewed ($[0.95, 0.05]$). Primary scale-exponent correlation (row 1) remained remarkably stable ($r > 0.9$) across all scenarios, indicating persistent structural confounding. In contrast, secondary location uncertainty ($fx2 \leftrightarrow fy2$, row 6) and primary intensity-weight confounding ($fz1 \leftrightarrow weight$, row 7) increase substantially with focus imbalance. Distinct threshold effects emerge in the extreme $[0.95, 0.05]$ mixture, where secondary dispersal parameters lost identifiability ($scale2 \leftrightarrow exponent2$ dropping to $-0.191$), suggesting a breakdown in the model's ability to resolve the minor component.

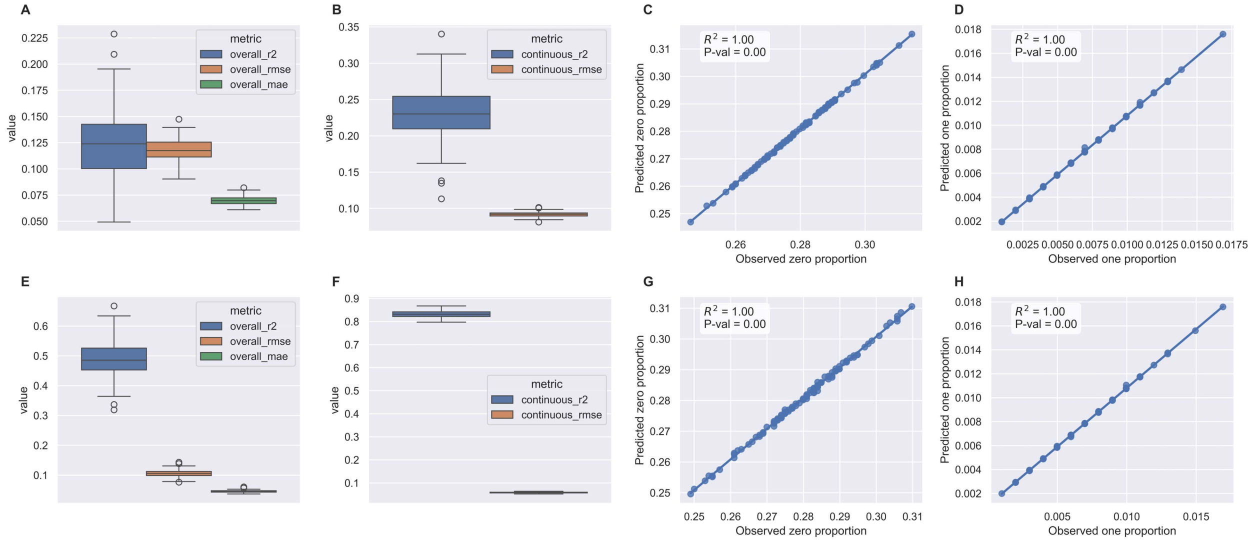


Figure S7: **Posterior predictive checks (PPC) and predictive performance of HiBASIL under model specification.** Rows correspond to two model misspecification scenarios: (A-D) Single-focus model applied to Two-foci data (under-parameterization), and (E-H) Two-foci model applied to Single-focus data (over-parameterization). Predictive performance is assessed via aggregate $R^2$, RMSE, and MAE (A, E), as well as specific $R^2$ and RMSE for the continuous Beta component (B, F). Scatter plots evaluate model calibration by comparing observed versus predicted proportions of zero-inflation (C,G) and one-inflation (D, H).

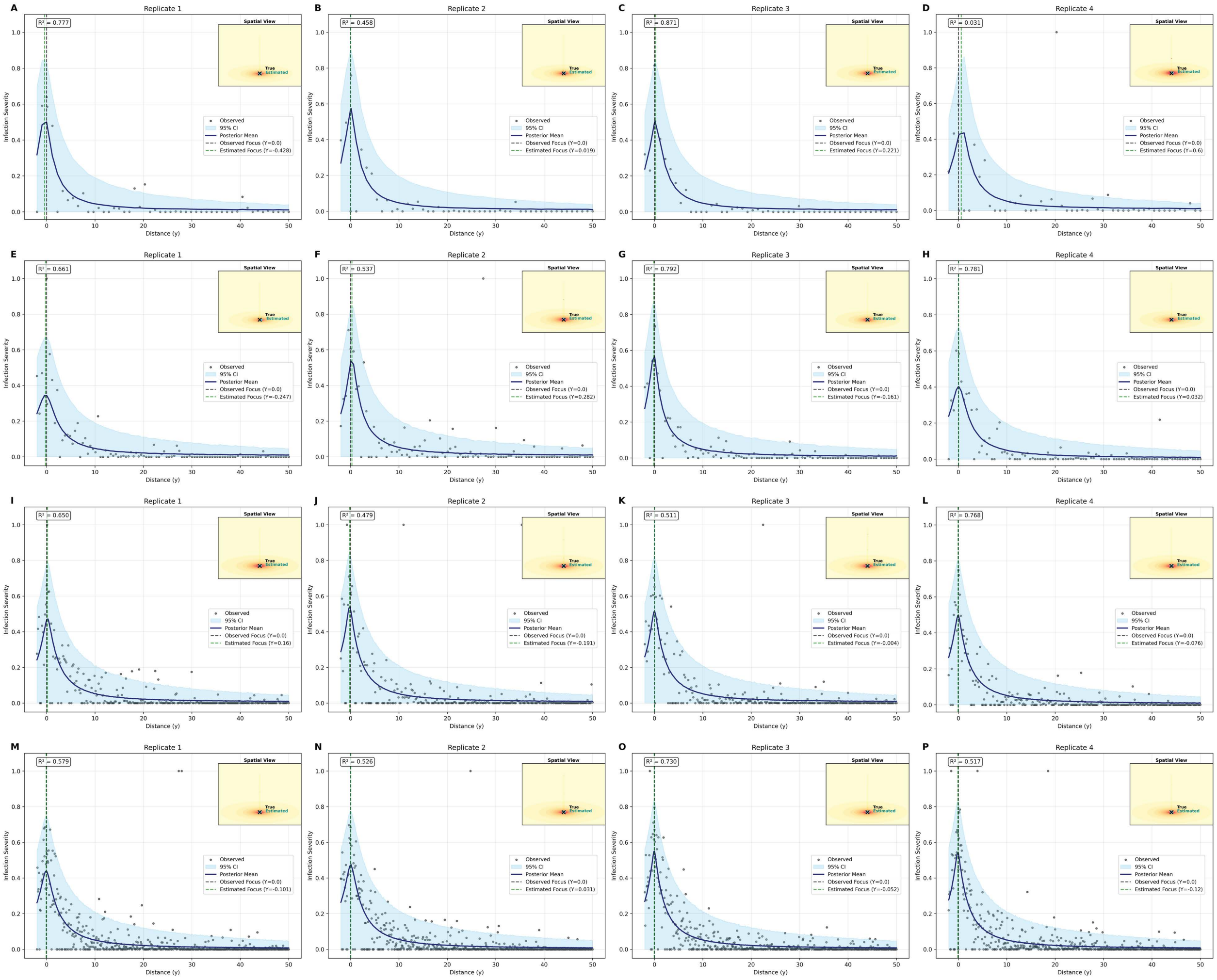


Figure S8: **Spatial validation of posterior predictive distribution of HiBASIL for single-focus scenarios across varying sample sizes.** Main panels illustrate 1D spatial transects at $x = 0$, comparing observed severity (black points) against the posterior predictive mean (navy line) and 95% credible interval (shaded sky-blue region). Simulation scenarios are organized by row:(A-D) $N = 50$, (E-H) $N = 100$, (I-L) $N = 250$, and (M-P) $N = 500$. Vertical dashed lines denote the estimated source locations at $y = 0$. The 2D inserts visualize the framework's capacity for joint parameter estimation and spatial source localization, specifically highlighting the estimation unknown infection sources (stars) relative to the true foci (crosses) under weakly informative priors.

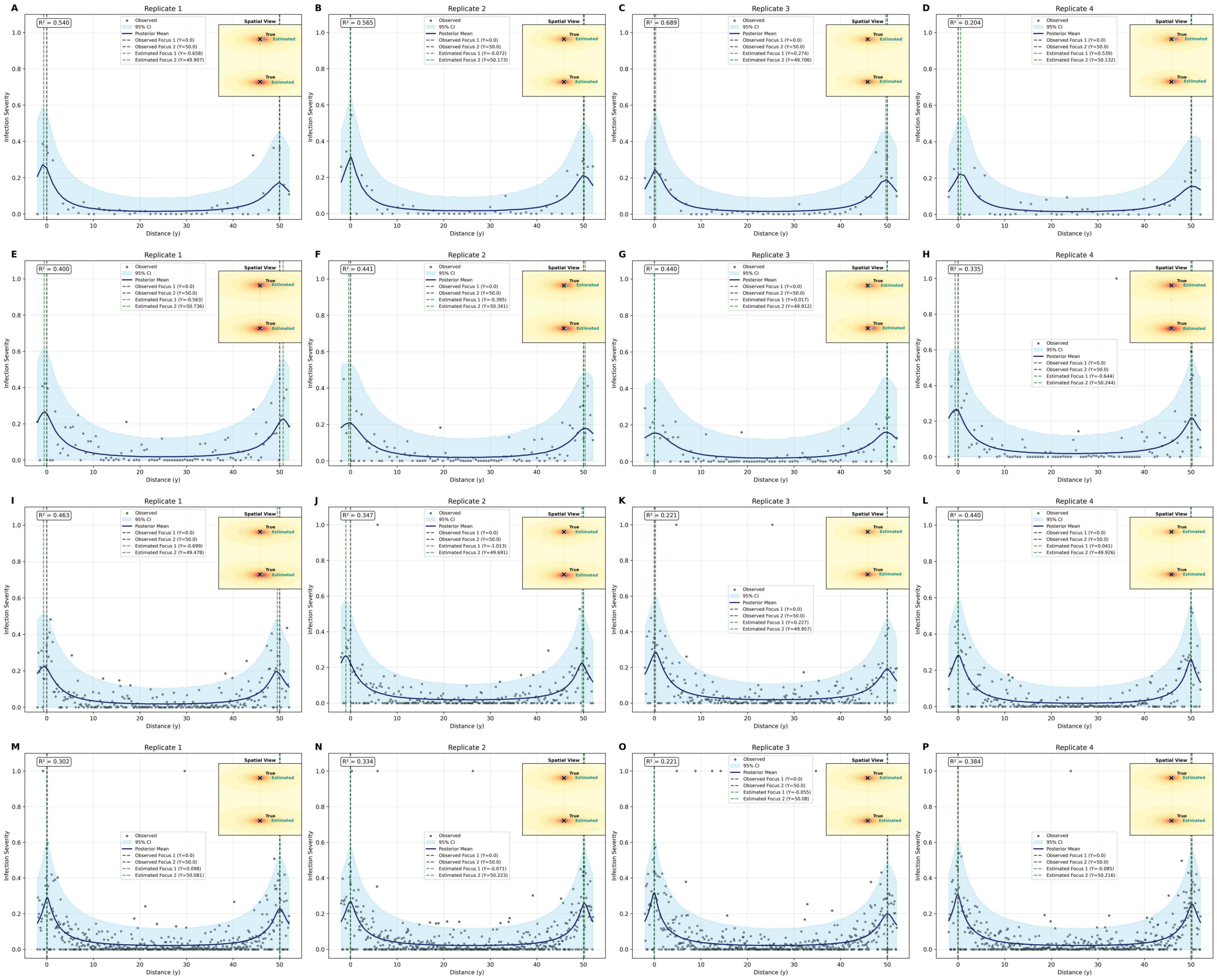


Figure S9: **Spatial validation of posterior predictive distribution for two-foci scenarios across varying sample sizes.** Main panels present 1D spatial transects at $x = 0$ for a complex two-foci system, comparing observed severity (black points) against the posterior predictive mean (navy line) and 95% credible interval (shaded sky-blue region). Scenarios are presented in increasing order of data density:(A-D) $N = 50$, (E-H) $N = 100$, (I-L) $N = 250$, and (M-P) $N = 500$. Vertical dashed lines indicate estimated source locations at $y = 0$ and $y = 50$. Corresponding 2D insets demonstrate the performance of the Bayesian framework in identifying multiple sources of inoculum simultaneously, especially estimate unknown infection sources (stars) relative to the true foci (crosses) using weakly informative priors.

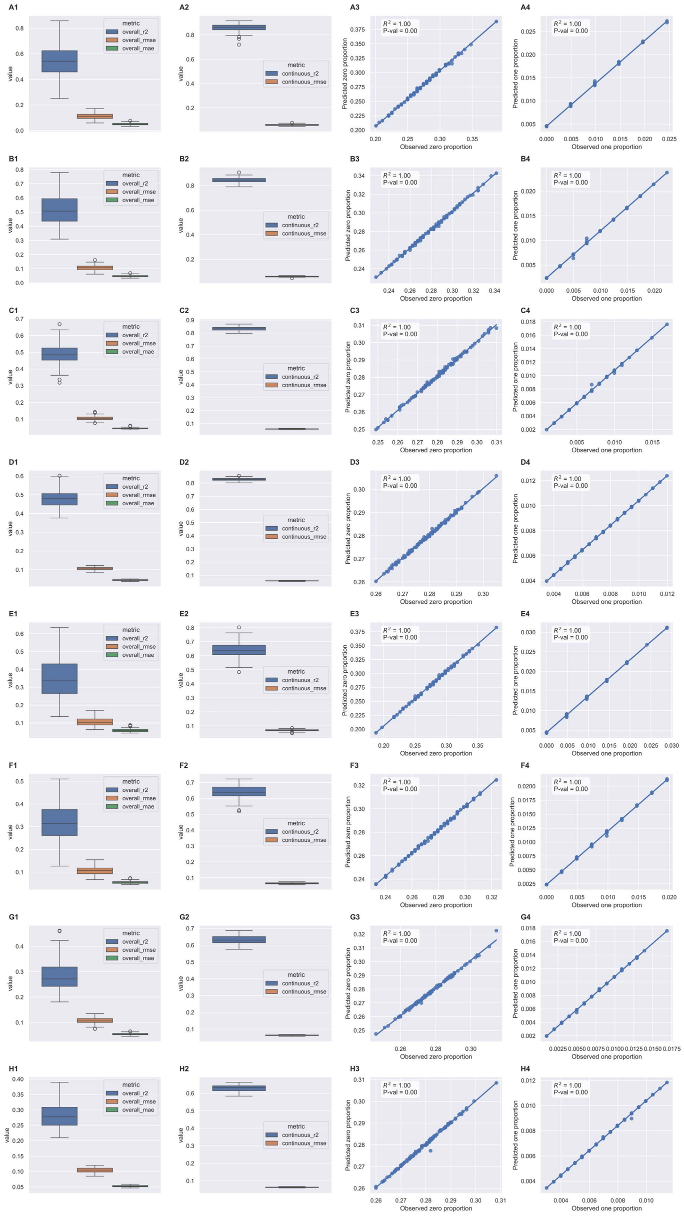


Figure S10: **Posterior predictive checks (PPC) and predictive performance of HiBASIL across gradients of spatial complexity and sample sizes.** Rows correspond to performance for specific simulation scenarios: Single-focus model with sample sizes $N = 50$ (A1-4), $N = 100$(B1-4), $N = 250$ (C1-4), and $N = 500$ (D1-4); and Two-foci models with sample sizes $N = 50$, $N = 100$ (F1-4), $N = 250$ (G1-4), and $N = 500$ (H1-4). Predictive performance is assessed via global $R^2$, RMSE, and MAE (A1-H1), while panels (A2-H2) show specific $R^2$ and RMSE for the continuous Beta component. Scatter plots evaluate model calibration by comparing observed versus predicted proportions of zero-inflation (A3-H3) and one-inflation (A4-H4). Calibration accuracy remain high and stable across all scenarios ($R^2 = 1.0$).

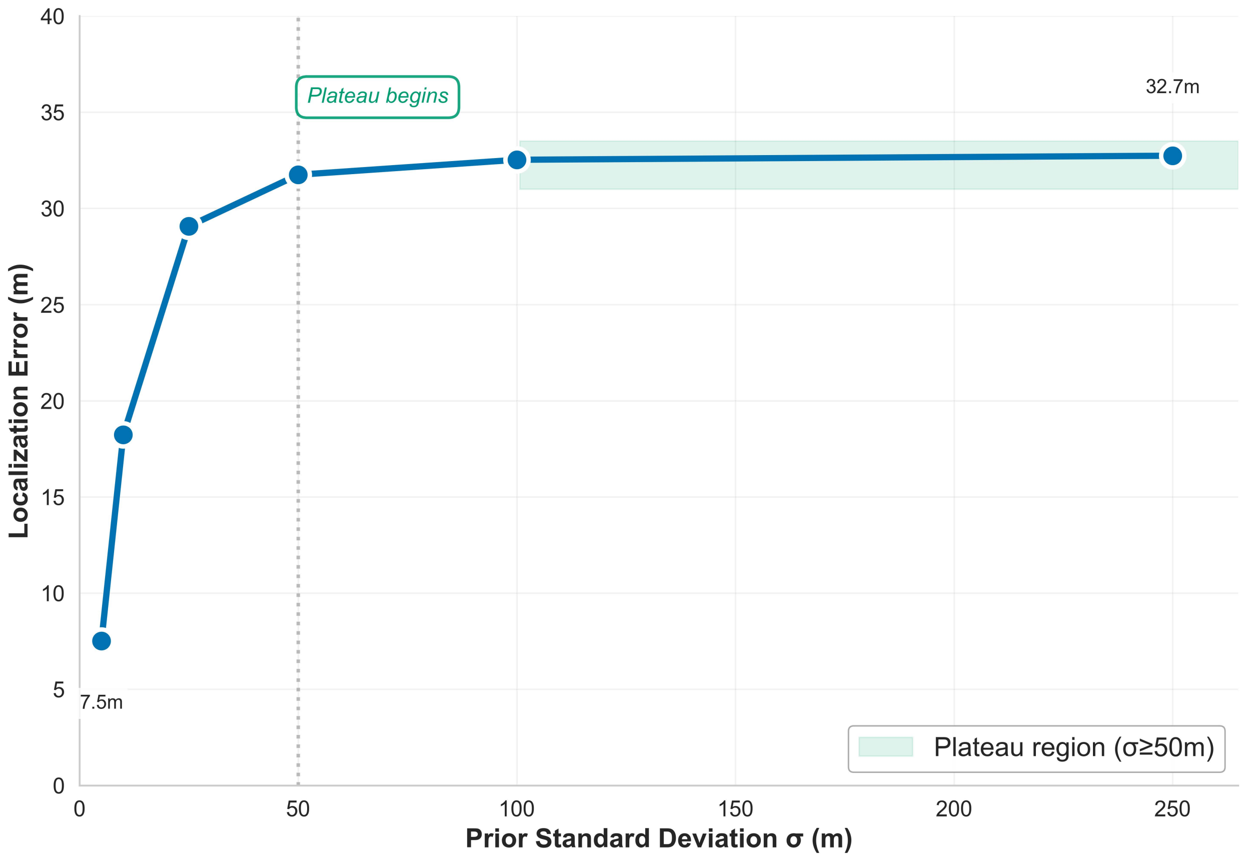


Figure S11: **Robustness of source estimation to prior specification.** The inverse L-shaped sensitivity curve demonstrates that the Single-Focus Power Law model is robust to initialization uncertainty. While pinpoint accuracy ($< 10m$ requires tightly constrained priors ($\sigma < 25m$), the model achieves a stable data-driven baseline of $\approx 32.7m$ for all diffuse priors ($\sigma \geq 50m$).

### S1 Dataset S1: Null Model Validation

**Model Selection Results for Null and Likelihood-decoupled Dispersal Models on Non-Spatial Data.** This dataset contains the leave-one-out cross-validation (LOO-CV) results for 100 simulated null datasets (no spatial structure) comparing the null model ($M_{\text{null}}$) against three likelihood-decoupled dispersal models ($M_{\text{exp}}$, $M_{\text{gau}}$, and $M_{\text{pow}}$). These models fit dispersal kernels independently without mechanistic coupling to source locations, serving as a baseline to demonstrate HiBASIL's integrated approach. Results indicate that $M_{\text{null}}$ is correctly identified when no spatial structure exists (specificity) (File: `./supplements/non_spatial_null_model_likelihood_decoupled_dispersal_models.csv`)

### S2 Dataset S2: Under-Parameterization

**Posterior Bimodality Analysis (Under-parameterized Scenario).** This CSV file contains the results of the *find_peaks* algorithm applied to the 100 simulations (400 replicates) of the Single-focus model applied to Two-foci data. It includes peak locations, distances between peaks, and modality classifications. (File: `./supplements/misspecification_scenario1_powerlaw.csv`)

### S3 Dataset S3: Over-parameterization

**Posterior Unimodality Analysis (Over-parameterized Scenario).** This CSV file contains the peak-finding results for Two-foci model applied to Single-focus data. It details the convergence of two-foci paramters onto a single ground-truth location. (File: `./supplements/misspecifica`

### S4 Supplementary Note 1: Model Specificity and Prior Independence

This note provide the full validation for HiBASIL's specificity and its behavior under null spatial conditions. The objective was to confirm that the model selection process effectively penalizes unnecessary complexity and that the dispersal parameters do not drift toward spurious spatial patterns when the underlying data is spatially random.

##### Methods for Control Analysis

The control phase evaluates the specificity and structural integrity of HiBASIL. Specifically, we aimed to verify that the model does not identify spurious (false positive) spatial patterns and to confirm that parameter estimation is correctly driven by the likelihood, allowing the model to simplify to a null state when spatial complexity is absent. To isolate the framework's sensitivity to spatial structure, we fixed the mean severity ($\mu$) to its known generative value. This component-wise validation strategy eliminates parameter confounding, ensuring that the model selection process is evaluated solely on its ability to distinguish between spatial dispersal kernels and non-spatial noise.

1. **Non-spatial Null Model ($M_{null}$):** This model assumes that disease severity and occurrence are uniform across the field and independent of distance. In this configuration, the mean severity ($\mu$) is fixed as a deterministic constant equal to the generative mean ($f_{z1}$) used in the simulation. The model estimates three global parameters as constant intercepts: the baseline zero-inflation ($\pi_0$), the one-inflation ($q$), and the precision ($\varphi$). This model serves as the baseline for assessing specificity.

2. **Likelihood-Decoupled Dispersal Models ($M_{exp}$, $M_{gau}$, $M_{pow}$)** To assess inference integrity, these models incorporate spatial kernels that are mathematically defined but strategically decoupled from the likelihood. Consistent with $M_{null}$, the mean severity ($\mu$) remains fixed to the true simulated constant $f_{z1}$. These models include kernel-

specific parameters: the **scale** ($a$) for the Exponential and Gaussian kernels, and both the **scale** ($a$) and **exponent** ($b$) for the Power Law kernel. Because these parameters are decoupled from the likelihood, the MCMC sampler should recover their respective prior distributions. This also allow us to verify that the inclusion of inactive spatial kernels does not bias the estimation of the ZOIB parameters (zero-inflation ($\pi_0$), one-inflation ($q$), and precision ($\varphi$)).

**Control Validation Procedure** To simulate a scenario with no underlying dispersal gradient (i.e., spatially random infection), we generated 100 independent null disease datasets using the simulation framework described for the non-spatial control generation. Each dataset was fitted to the four competing models ($M_{null}$, $M_{exp}$, $M_{gau}$, and $M_{pow}$) to assess the following:

- **Model Selection and Specificity:** We conducted a formal model comparison using LOO-CV, as detailed in subsection 2.3, to assess whether the framework correctly identifies the absence of spatial structure. We tested the hypothesis that the additional parameters in the dispersal models ($M_{exp}$, $M_{gau}$, and $M_{pow}$) would incur a complexity penalty without offering predictive improvement over the null ($M_{null}$). This hypothesis supported if $M_{null}$ were consistently identified as the most parsimonious model, or if the dispersal models provide no significant predictive gain (i.e., $|\Delta ELPD_{loo}| \leq 2 \times SE$). Furthermore, we performed Posterior Predictive Checks (PPC) on $M_{null}$ to evaluate wether the global parameters ($\pi_0$, $q$, $\varphi$) effectively captured the empirical distribution. This validates the framework's specificity and prevents the misidentification of stochastic noise as spatial structure.

- **Estimation Accuracy and Robustness:** To verify that the inclusion of inactive spatial kernels did not degrade the framework's core inference accuracy, we evaluated the estimation of the ZOIB parameters ($\pi_0$, $q$, $\varphi$). Performance was quantified using Mean Bias and 95% credible interval coverage to ensure the recovery of generative values. We monitored the Potential Scale Reduction Factor ($\hat{R}$) and Bulk-ESS to confirm that the sampler maintained high efficiency and convergence even while navigating the redundant parameter space created by the decoupled kernels.

- **Prior-to-Posterior Shift and Integrity:** To quantify the relative influence of priors on the decoupled parameters, we calculated the shift and distribution overlap between prior and posterior distributions. The mean shift was defined as the absolute difference between the prior and posterior means ($\Delta\mu = |\mu_{\text{prior}} - \mu_{\text{posterior}}|$). Distributional overlap was measured as the percentage of area shared by the respective probability density functions. A simulation was defined as successful if the posterior mean for dispersal parameters (scale and exponent) exhibited a relative shift of $< 10\%$ from the prior mean and maintained an overlap of $> 80\%$, indicating that these parameters remained prior-driven in the absence of a likelihood signal.

#### Control Validation Results

To ensure a strictly controlled comparison, we standardized the parameter space across all candidate models by fixing the mean severity to the generative constant. By eliminating mean-level estimation uncertainty as a potential confounding factor, the cross-validation process was able to focus exclusively on the structural differences between the non-spatial null and the spatial kernels. Under this unified framework, the effective number of parameters ($p_{loo}$) for the null model converged to values nearly identical to those of the spatial alternatives (2.870, 2.865, 2.865, 2.882 for $M_{null}$, $M_{exp}$, $M_{gau}$, and $M_{pow}$, respectively). This confirms that subsequent model selection reflects differences in spatial likelihood performance rather than discrepancies in parameter count or model complexity.

Preliminary validation via PPC for the non-spatial null model ($M_{null}$) showed that the framework correctly identifies the absence of spatial structure in the control datasets (Figure S12). The overall $R^2$ is tightly clustered at zero ($-0.00033 \pm 0.00055$), and the continuous $R^2$ is appropriately negative (Figure S12A&B). These metrics indicate that the null model does not overfit stochastic noise or attribute spurious explanatory power to random infection

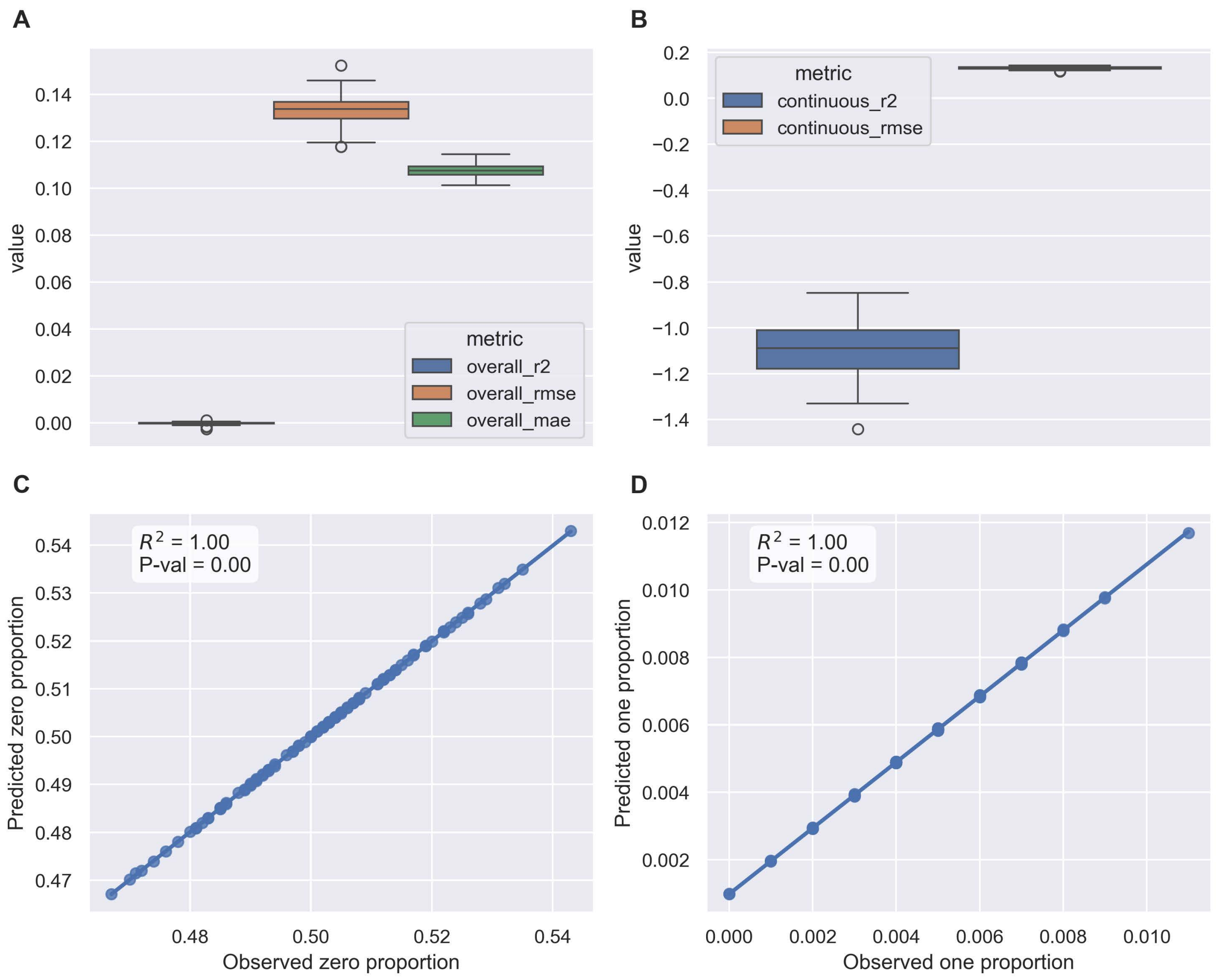


Figure S12: **Posterior Predictive Check (PPC) for the Non-Spatial Null Model ($M_{null}$).** (A) Distribution of aggregate $R^2$, RMSE and MAE across 100 null simulations. (B) Performance metrics for the continuous (Beta) component of the ZOIB model. (C) Correspondence between observed and predicted proportions of zero-severity. (D) Correspondence between observed and predicted proportions of one-severity. The 1:1 identity line represents perfect model calibration, confirming accurate estimation of global parameters.

patterns. Furthermore, the model demonstrates high calibration across the ZOIB components: the observed versus predicted proportions for both zero- and one-severity values align precisely with the 1:1 identity line ($R^2 = 1.0$) (Figure S12C&D). This perfect recovery of discrete components indicates that the global parameters ($\pi_0$, $q$, $\varphi$) are accurately estimated, providing a robust baseline for model comparison.

Moreover, the framework demonstrated high specificity through its consistent identification of model parsimony. Across 100 independent null simulations, LOO-CV revealed that the non-spatial model ($M_{null}$) was identified as either the top-ranked model or predictively equivalent to the winner in 80% of the trials. The exponential kernel ($M_{exp}$) met this threshold in 91% of the trials; notably, the Gaussian kernel $M_{gau}$ performed identically to $M_{exp}$ due to their shared priors and parameterization in this validation setup; meanwhile, the power law kernel $M_{pow}$ reached predictive equivalence in 59% of trials. While the decoupled spatial models ($M_{exp}$ and $M_{gau}$) achieved marginally higher ELPD point estimates than $M_{null}$ in 55 trials due to stochastic noise, the mean difference ($\Delta ELPD$) between $M_{null}$ and $M_{exp}$ remained negligible at -0.0047 (SE = 0.008). Because the magnitude of the difference is significantly less than the two standard errors ($|\Delta ELPD_{loo}| < 2 \times SE$), the models are considered predictively indistinguishable. This demonstrates that despite the identical effective complexity ($p_{loo}$) across candidates, the framework successfully prioritized the simpler null hypothesis in the absence of a spatial signal, effectively preventing the misidentification of spurious spatial patterns (see Supplementary File dataset S1).

The framework's integrity was further validated by confirming that in the absence of a spatial signal, the posterior estimates for dispersal parameters do not deviate significantly from their priors. Across the 100 null simulations, the posterior distributions for all kernel types (Exponential, Gaussian, and Power Law) exhibited high overlap with the priors and negligible mean shifts, achieving a 100% success rate (Table S14). For the Power Law kernel, which possesses the highest parameter complexity, the posterior means for the scale ($\mu = 15.96 \pm 0.08$) and exponent ($\mu = 2.00 \pm 0.01$) showed negligible shift from their prior centers (Table S14, Figure S13). This high degree of distribution overlap (94.8%, 94.7%) indicates that the likelihood for those parameters is non-informative under null conditions. Consequently, the posterior is effectively regularized by the prior, preventing the framework from converging on spurious spatial patterns (Figure S13).

Table S14: **Summary of Prior-Posterior Overlap for Dispersal Parameters.** Values represent the mean ± standard deviation across 100 null simulations. Success rate indicates that every simulation met the criteria of <10% relative shift and >80% distribution overlap, confirming that the model did not infer spurious spatial information from random noise.

| Model | Parameter | Prior Mean | Posterior Mean (avg) | Mean Shift | Overlap | Success Rate |
|---|---|---|---|---|---|---|
| Exponential | Scale (a) | 15.58 | 15.92 ± 0.07 | -0.02 ± 0.36 | 94.7% ± 0.7% | 100.00% |
| Gaussian | Scale (a) | 15.58 | 15.92 ± 0.07 | -0.02 ± 0.36 | 94.7% ± 0.7% | 100.00% |
| Power Law | Scale (a) | 15.58 | 15.96 ± 0.08 | 0.02 ± 0.36 | 94.8% ± 0.6% | 100.00% |
| Power Law | Exponent (b) | 2.03 | 2.00 ± 0.01 | 0.01 ± 0.03 | 94.7% ± 0.7% | 100.00% |

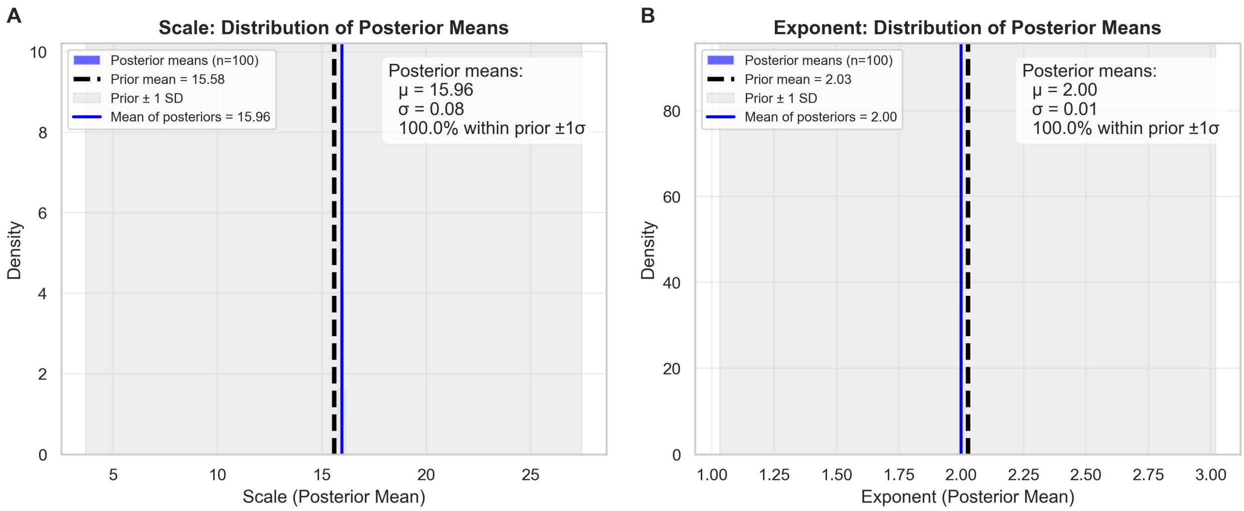


Figure S13: **Distribution of posterior means for Power Law dispersal kernel parameters across 100 null simulations.** (A) Scale parameter ($\mu_{prior} = 15.58$): The aggregate posterior mean ($\mu_{post} = 15.96 \pm 0.08$) shows a negligible shift from the prior, with 100% of trials falling within one standard deviation of the prior mean. (B) Exponent parameter ($\mu_{prior} = 2.03$): The exponent posterior similarly remains centered at $2.00 \pm 0.01$.

### S5 Supplementary Note 2: Field Experiment Protocols and Goodness of Fit for the ZOIB Distribution

#### Experiment Design and Site Description

The field experiment was conducted at the Williamsdale Farm Extension and Research Center of North Carolina State University (34 °45'46"N, 78 °5'59"W). Zucchini squash (*Cucurbita pepo* L. cv. 'Spineless Perfection F1'; Harris Seeds, Rochester, NY, USA) served as the study host plant. Seeds were direct-sown on May 18, 2023. Treatments were laid out a Randomized Complete Block Design (RCBD) with four replicates. Each block comprised five treatment subplots: (i) single-focus, inoculated once; (ii) single-focus, inoculated twice; (iii) two-foci, inoculated once; (iv) two-foci, inoculated twice; and (v) untreated control (Figure S14). Subplot measured 26 m $\times$ 10 m, separated by a 20 m inter-subplot buffer within blocks and a 4.9 m gap between blocks. Each subplot contained five rows with 2-ft (0.61 m) intra-row plant spacing and 10-ft (3.05 m) inter-row spacing. Artificial inoculation foci locations were designed by red and black circles as illustrated in Figure S14.

#### Disease Assessment

Approximately one month after planting, plants were sampled at 1.83 m intervals along the disease assessment survey row. For each sampled plant, five leaves were randomly selected and marked with colorful ribbons to ensure consistent disease assessment throughout the experiment. Inoculation was performed by spraying 15 mL of fresh *P. cubensis* sporangia suspension (isolate D3, lineage I; $> 1.0 \times 10^4$ sporangia / mL) to the designated focus plant. Following inoculation, each inoculated plant was covered with a black plastic bag overnight to maintain high humidity necessary for successful infection. For treatments requiring a second inoculation, six laboratory-infected squash plants were deployed around each primary focus to serve as additional inoculation sources.

Disease severity was evaluated as the percentage of leaf area displaying chlorotic and

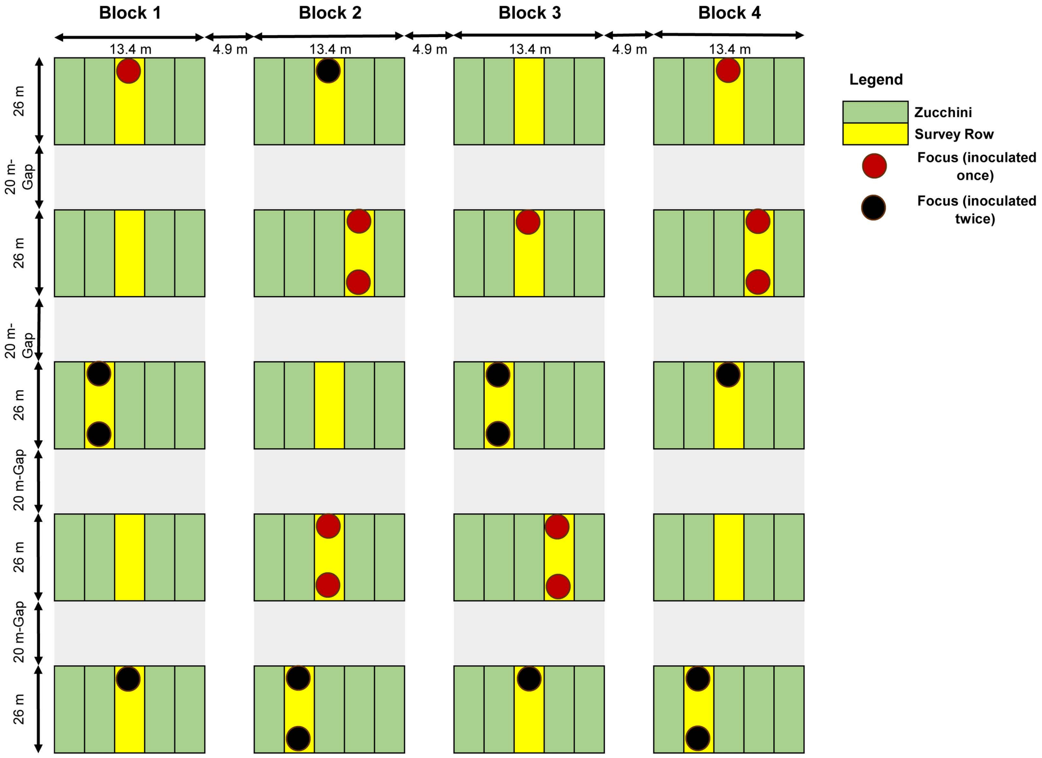


Figure S14: **Experimental layout and treatment allocation in 2023.** The field experiment was established using a randomized complete block design with four replicates. Treatments included: untreated control; single-focus (inoculated once); single-focus (inoculated twice); two-foci (inoculated once), and two-foci (inoculated twice). Each subplot consisted of five rows of zucchini (*Cucurbita pepo*). Focus symbols indicate inoculation methods: red circles represent a single sporangial suspension spray, while black circles denote an initial spray followed by the introduction of sporulating plants. The surveyed rows are highlighted in yellow. Subplot dimensions include the plastic mulch bed, whereas inter-block gaps represent the distance between mulch beds.

necrotic symptoms. Disease assessment started 5 to 7 days post-inoculation and were repeated every three to four days, totaling six disease records over a 3-week period. Disease severity of each tagged leaf and its corresponding plant location were recorded. Zucchini fruits were harvested regularly throughout the study to prevent physiological leaf yellowing due to fruit maturation. Data collected during the third survey were utilized for subsequent distribution fitting and dispersal modeling.

#### Fitting ZOIB Distribution by Maximum Likelihood Estimation

The suitability of the ZOIB distribution was evaluated for each treatment using maximum likelihood estimation (MLE). Parameters ($p$, $q$, $\alpha$, $\beta$, and $\varphi$) were estimated by minimizing the negative log-likelihood using the Differential Evolution algorithm (Storn, 2015) via the scipy.optimize module (Virtanen et al., 2020). A Kolmogorov-Smirnov (KS) test (Massey, 1951) was employed to verify that the distribution sufficiently captured the zero- and one-inflation, variance, and skewness of the severity data. The significance of the KS test was determined via bootstrapping with 10,000 iterations.

#### Goodness of Fit Results for the ZOIB Distribution

The ZOIB distribution successfully characterized the disease severity for the majority of treatments ($P > 0.15$ in 4 out of 5 groups; Table S15; Figure S15). A marginal deviation was observed in the single-focus (inoculated once) treatment ($P = 0.03$; Table S15), likely due to the inherent stochasticity of disease establishment at lower inoculum pressures; however, visual inspection confirmed that the model accurately captured the zero-inflation and non-constant variance inherent of disease severity (Figure S15). Given the overall high goodness-of-fit, the ZOIB was considered statistically appropriate for the subsequent hierarchical Bayesian dispersal analysis.

Table S15: **Estimated parameters for ZOIB distribution for disease severity by treatment (2023).**

| **Treatment** | **Observed** $p$ | **Observed** $q$ | **Estimated** $\alpha$ | **Estimated** $\beta$ | **Estimated** $\varphi$ | **Estimated** $p$ | **Estimated** $q$ | **KS statistic** | **KS** $P$**-value** |
|---|---|---|---|---|---|---|---|---|---|
| Control | 0.20 | 0 | 1.10 | 43.05 | 44.15 | 0.20 | 0 | 0.19 | 1.00 |
| Single focus (inoculated once) | 0.16 | 0 | 0.50 | 6.04 | 6.53 | 0.16 | 0 | 0.21 | 0.03 |
| Single focus (inoculated twice) | 0.02 | 0 | 0.52 | 2.14 | 2.66 | 0.02 | 0 | 0.14 | 0.17 |
| Two foci (inoculated once) | 0.03 | 0 | 0.80 | 4.78 | 5.57 | 0.03 | 0 | 0.12 | 0.29 |
| Two foci (inoculated twice) | 0.00 | 0 | 0.70 | 3.01 | 3.70 | 0.00 | 0 | 0.13 | 0.21 |

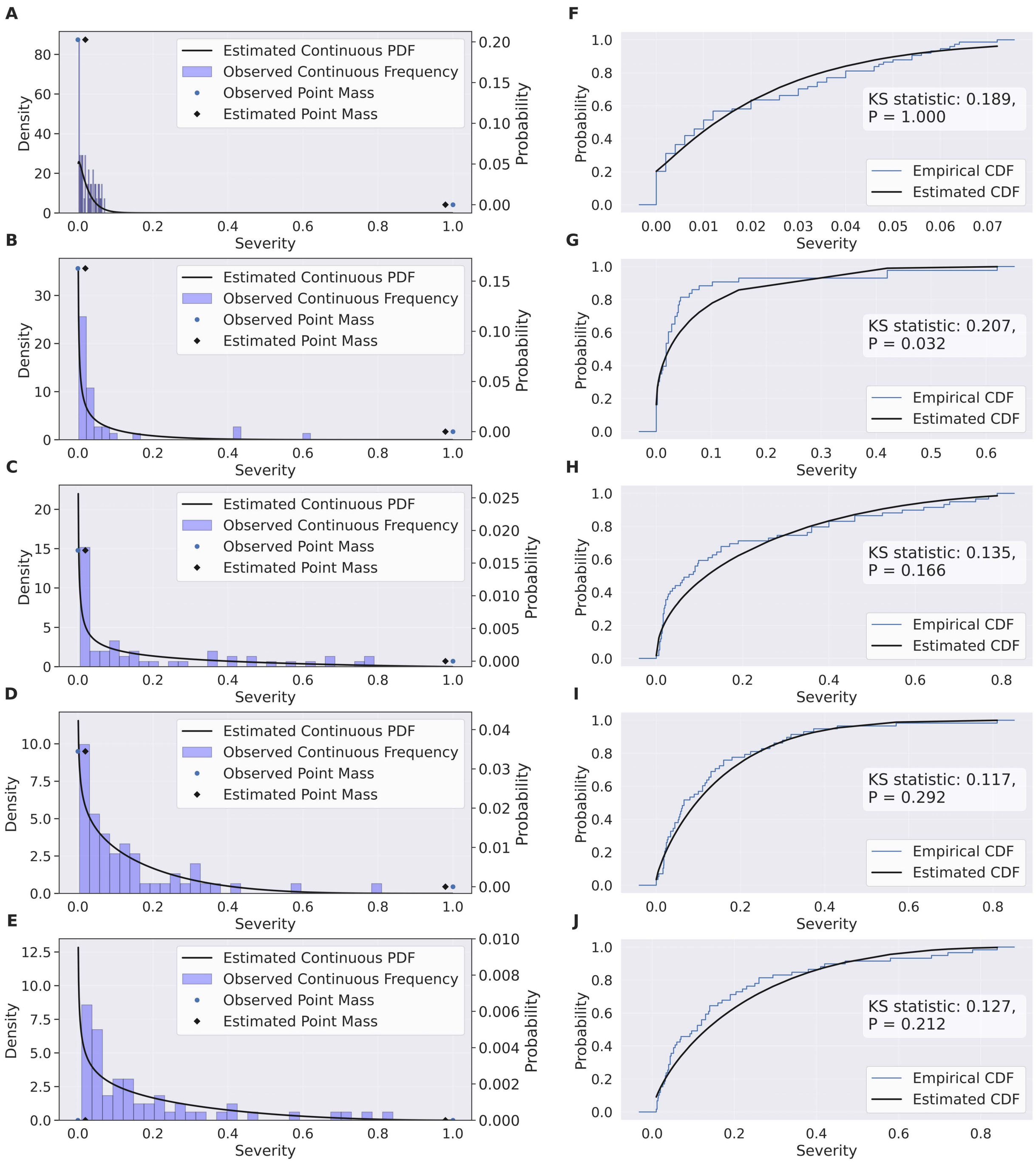


Figure S15: **Zero-one-inflated beta (ZOIB) model fits and Kolmogorov-Smirnov (KS) goodness-of-fit assessments for 2023 survey data.** Left panels (A-E) illustrate the observed frequency distributions with estimated probability density function and point masses. Right panels (F-J) show corresponding KS tests, comparing the empirical and estimated cumulative distribution functions. Rows represent experimental treatments: control (A,F); single-focus, inoculated once (B,G) or twice (C,H); two-foci, inoculated once (D,I), or twice (E,J).

### S6 Supplementary Note 3: Field Experiments Baseline Calibration and Model Sensitivity

#### Control Data (Specificity)

For the non-treatment control data, we evaluated the dispersal models against a baseline Null model, which assumed no spatial dependence in disease severity. Model selection via leave-One-Out Cross-Validation (LOO-CV) identified the Null model as the superior fit ($rank_{loo} = 1$), followed by the Two-foci Power Law and the Single-focus Power Law models. While the differences in expected log predictive density (ELPD) among these top candidate models were within two standard errors ($\Delta ELPD_{loo} < 2 \times SE$), the Null model exhibited the lowest effective number of parameters ($p_{loo} = 4.428$), and thus the most parsimonious model (Table S16).

We also assessed the raw predictive performance of each model (Table S16). As expected with increased complexity, the Two-foci Exponential model yielded the highest coefficients of determination (overall $R^2 = 0.533$; continuous $R^2 = 0.479$), followed closely by the Two-foci Power Law model (overall $R^2 = 0.521$; continuous $R^2 = 0.467$). In contrast, the Null model showed lower raw predictive metrics (overall $R^2 = 0.426$; continuous $R^2 = 0.354$). However, the RMSE for the overall model and the continuous Beta component differed by only 0.001 or 0.002 across all candidate models. Additionally, visual inspection of the posterior predictive distributions (Figure S16; Figure S17) revealed that the spatially explicit models produced flat gradients virtually indistinguishable from the Null model. This lack of distinct spatial structure indicates that the higher $R^2$ values in complex models likely reflect overfitting to noise rather than a biological signal. Consequently, based on LOO-CV and spatial interpretation, the Null model was selected as the appropriate baseline for the non-treatment control.

Table S16: Predictive Performance and Leave-One-Out Cross Validation Results for Non-treatment Control Data of Cucurbit Downy Mildew Field Experiments.

| **Metric** | **Null Model** | **Single-focus Model** | | | **Two-foci Model** | | |
|---|---|---|---|---|---|---|---|
| | | Exponential | Gaussian | Power Law | Exponential | Gaussian | Power Law |
| overall $R^2$ | 0.426 | 0.476 | 0.473 | 0.491 | **0.533** | 0.484 | 0.521 |
| overall RMSE | 0.016 | 0.015 | 0.015 | 0.015 | **0.014** | 0.015 | 0.015 |
| overall MAE | 0.013 | 0.013 | 0.013 | 0.013 | **0.012** | **0.012** | **0.012** |
| obs_zeros | 0.203 | 0.203 | 0.203 | 0.203 | 0.203 | 0.203 | 0.203 |
| pred_zeros | 0.238 | 0.239 | 0.238 | 0.238 | 0.238 | 0.238 | 0.238 |
| obs_ones | 0.000 | 0.000 | 0.000 | 0.000 | 0.000 | 0.000 | 0.000 |
| pred_ones | 0.007 | 0.007 | 0.007 | 0.007 | 0.007 | 0.007 | 0.007 |
| continuous $R^2$ | 0.354 | 0.405 | 0.401 | 0.429 | **0.479** | 0.416 | 0.467 |
| continuous RMSE | 0.017 | 0.016 | 0.016 | 0.016 | **0.015** | 0.016 | **0.015** |
| n_obs | 74 | 74 | 74 | 74 | 74 | 74 | 74 |
| elpd_loo | **148.010** | 144.623 | 146.593 | 147.146 | 146.165 | 146.473 | 147.726 |
| p_loo | **4.428** | 6.010 | 5.356 | 5.290 | 6.114 | 5.696 | 5.395 |
| rank_loo | **1** | 7 | 4 | 3 | 6 | 5 | 2 |
| se_d_loo | 0.000 | 1.127 | 0.786 | 0.665 | 0.937 | 0.800 | 0.644 |
| warning | FALSE | FALSE | FALSE | FALSE | FALSE | FALSE | FALSE |
| pareto_k_bad | 0 | 0 | 0 | 0 | 0 | 0 | 0 |

*Note:* Models were applied to non-treatment control data ($n = 74$). Bold values indicate the best performance in each category. The metric Pareto_k_bad counts observations with a Pareto shape parameter $k > 0.7$, which indicates highly influential data points or potential outliers.

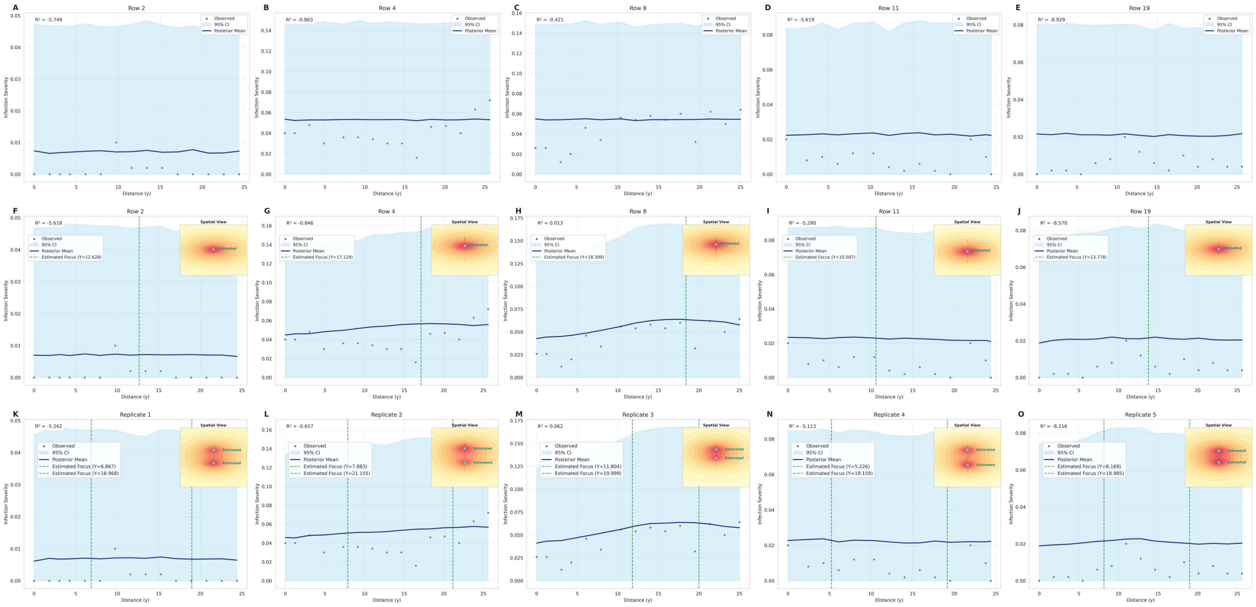


Figure S16: **Spatial validation of posterior predictive distribution for non-treatment control data.** Primary panels display 1D spatial transects at $x = 0$, comparing observed severity (black points) against the posterior predictive mean (navy line) and 95% credible interval (shaded sky-blue region). Rows correspond to the three top-ranked models:(A-E) the Null model, (F-J) the Single-focus Power Law model, and (K-O) the Two-foci Power Law model. Vertical dashed lines mark the estimated locations of infection sources along the transect. Corresponding 2D insets visualize the inferred spatial field and source locations (stars) as estimated within the Bayesian framework.

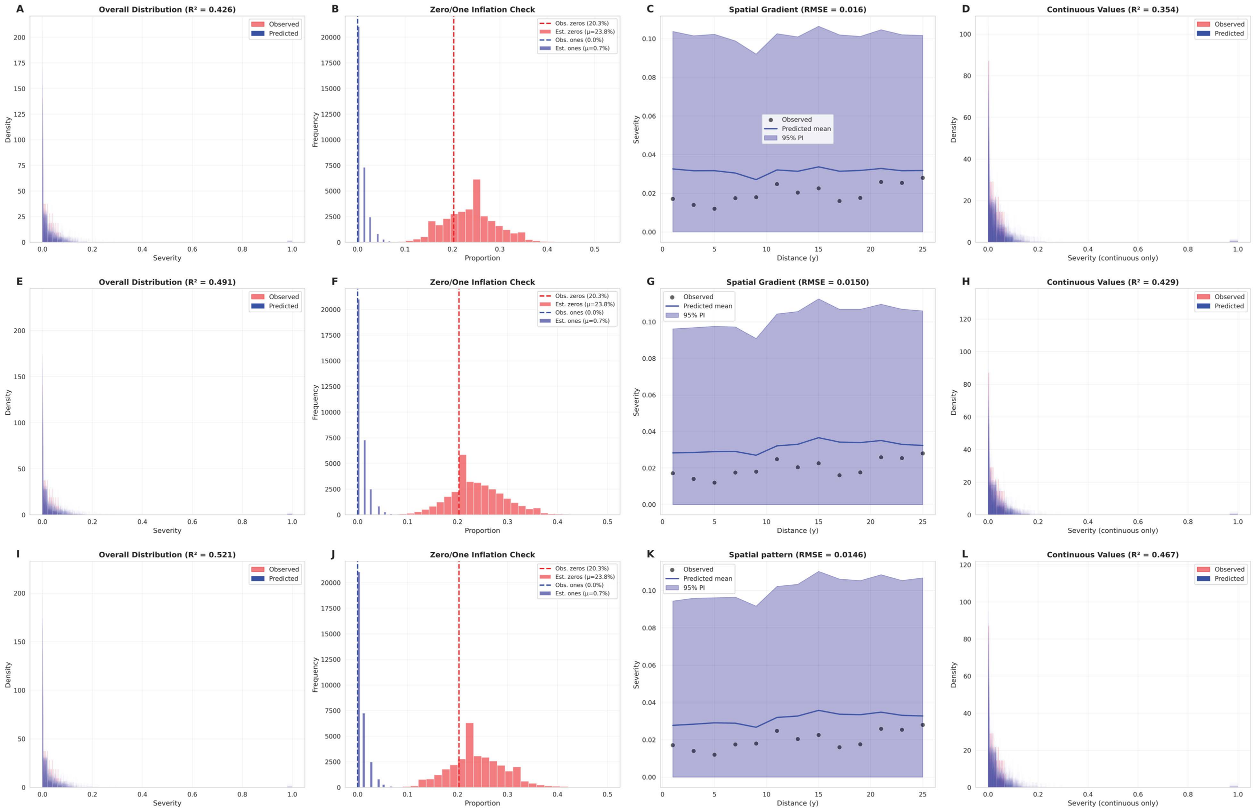


Figure S17: **Posterior predictive checks (PPC) and predictive performance for non-treatment control data.** Rows correspond to the three evaluated models: (A-D) the Null model, (E-H) the Single-focus Power Law model, and (I-L) the two-foci Power Law model. Columns display: Overall distribution of observed versus predicted severity with associated $R^2$ (A, E, I); Zero- and One-inflation checks comparing observed proportions (dashed lines) against model estimates (histograms) (B, F, J); Spatial gradients comparing observed binned-mean (2-meter bins) severity (points) with the posterior predicted mean (line) (C, G, K); and predictive distributions and $R^2$ for the continuous Beta component specifically (D, H, L).

#### Kernel Validation With Known Focus Location - Single Focus

**Model Selection and Predictive Baseline** To establish a baseline for the dispersal kernels without the confounding spatial uncertainty of source inference, we first fitted single-focus models (Exponential, Gaussian, and Power Law kernels) with fixed, known focus coordinates. Across both inoculated-once and inoculated-twice treatments, the Power Law model consistently exhibited the superior fit, characterized by the highest expected log predictive density (ELPD) and the most parsimonious effective number of parameters ($p_{loo}$) (Table S17). Consistent with the unknown location scenarios, the Power Law model decisively outperformed the Exponential and Gaussian kernels. In the inoculated-once treatment, the Power Law kernel reduced the overall RMSE by approximately 40-50% compared to alternative kernels (RMSE of 0.049 vs. 0.085 for Exponential and 0.112 for Gaussian). As expected, predictive accuracy was marginally higher in scenario of known focus location. These fixed-focus-location Power Law models yielded overall $R^2$ values of 0.844 and 0.875 for the inoculated-once and inoculated-twice treatments, respectively (compared to 0.65 and 0.83 in the joint estimation scenarios) (Table S17; Figure S19A – H). These baseline results confirmed that the Power Law is the biologically appropriate structural kernel for *P. cubensis* dispersal in our experimental system before introducing the spatial inference.

**Parameter Sensitivity and Estimation Trade-offs** Comparing parameter estimates between the known and unknown focus location scenarios reveals specific behavioral trade-offs within the hierarchical framework. When the focus location was jointly estimated, the inferred focus intensities ($f_{z1}$) were generally estimated slightly higher than in the known-location scenario (Table S18; Table 8). This suggests a compensatory trade-off during MCMC sampling: to account for slight spatial uncertainty while searching for the exact source coordinates, the model marginally inflates the focal intensity to ensure the likelihood adequately covers the observed spatial disease peak.

Similarly, dispersal scale parameters exhibited sensitivity to spatial uncertainty. In the inoculated-twice treatment, the known focus location models produced row-specific scales ranging from 1.42 to 11.18. Whereas the unknown focus location models showed expanded variance, ranging from 3.88 to 19.20. This indicates that the scale parameter sensitivity to the environmental heterogeneity when exact spatial origins are relaxed. Despite these minor shifts in latent parameters, visual inspection of the fitted spatial patterns revealed no significant difference in the final predictive surface; both known and unknown scenarios recovered the dispersal kernel with high fidelity (Figure S18; Figure 12F–L).

**Structural Stability of the ZOIB Framework** Crucially, the estimated probabilities for structural zeros (healthy plants, $\pi 0$) and structural ones (fully infected plants, $q$) remained remarkably consistent across known and unknown focus location scenarios (Table S18; Table 8). The stability of these mixture components, regardless of spatial coordinates uncertainty, demonstrates the robustness of the HiBASIL framework in partitioning healthy plants and fully infected plants from the continuous disease gradient.

Table S17: Predictive Performance and LOO-CV Results of Single-focus Models for Cucurbit Downy Mildew Field Experiments under **Known Focus Location** Scenarios.

| | **Single-focus (inoc. once)** | | | **Single-focus (inoc. twice)** | | |
|---|---|---|---|---|---|---|
| **Metric** | Exponential | Gaussian | Power Law | Exponential | Gaussian | Power Law |
| overall $R^2$ | 0.526 | 0.182 | **0.844** | 0.781 | 0.718 | **0.875** |
| overall RMSE | 0.085 | 0.112 | **0.049** | 0.108 | 0.122 | **0.082** |
| overall MAE | 0.067 | 0.084 | **0.042** | 0.084 | 0.096 | **0.061** |
| obs_zeros | 0.163 | 0.163 | 0.163 | 0.017 | 0.017 | 0.017 |
| pred_zeros | 0.180 | 0.180 | 0.181 | 0.028 | 0.028 | 0.028 |
| obs_ones | 0.000 | 0.000 | 0.000 | 0.000 | 0.000 | 0.000 |
| pred_ones | 0.018 | 0.018 | 0.017 | 0.014 | 0.014 | 0.014 |
| continuous $R^2$ | 0.517 | 0.164 | **0.851** | 0.782 | 0.718 | **0.875** |
| continuous RMSE | 0.092 | 0.121 | **0.051** | 0.108 | 0.123 | **0.082** |
| n_obs | 43 | 43 | 43 | 59 | 59 | 59 |
| elpd_loo | 42.786 | 38.175 | **53.244** | 67.731 | 64.674 | **77.890** |
| p_loo | 3.288 | 3.610 | **2.249** | 6.259 | 5.200 | **4.886** |
| rank_loo | 2 | 3 | **1** | 2 | 3 | **1** |
| se_d_loo | 1.460 | 2.784 | 0.000 | 3.559 | 5.099 | 0.000 |
| warning | False | False | False | True | False | True |
| Pareto_k_bad | 0 | 0 | 0 | 1 | 0 | 2 |

Table S18: Parameter Estimation Results of Single-focus Models for Cucurbit Downy Mildew Field Experiments under **Known Focus Location** (Fixed Coordinates).

| **Parameter** | **Inoculated once** | **Inoculated twice** |
|---|---|---|
| *Focus Location* | | |
| Focus $x$ ($f_{x1}$) | Fixed | Fixed |
| Focus $y$ ($f_{y1}$) | Fixed | Fixed |
| *Focus Intensity ($f_{z1}$)* | | |
| **Global Mean** $f_{z1}$ | $0.49 \pm 0.12$ | $0.60 \pm 0.12$ |
| *Row-Specific Intensity:* | | |
| $f_{z1}$ - Row 0 | $0.41 \pm 0.11$ | $0.77 \pm 0.09$ |
| $f_{z1}$ - Row 1 | $0.46 \pm 0.10$ | $0.49 \pm 0.14$ |
| $f_{z1}$ - Row 2 | $0.58 \pm 0.11$ | $0.65 \pm 0.11$ |
| $f_{z1}$ - Row 3 | — | $0.51 \pm 0.14$ |
| *Dispersal Parameters* | | |
| Exponent (Decay) | $1.30 \pm 0.28$ | 0.90 (0.29) |
| Scale (Global) | $2.39 \pm 1.10$ | — $a$ |
| Global Mean Scale ($scale_{\mu}$) | — | $1.51 \pm 0.81$ |
| Scale Variance ($scale_{\sigma}$) | — | $1.38 \pm 0.5$ |
| Scale - Row 0 | — | $11.18 \pm 7.17$ |
| Scale - Row 1 | — | $1.42 \pm 3.90$ |
| Scale - Row 2 | — | $3.18 \pm 3.30$ |
| Scale - Row 3 | — | $5.70 \pm 281.15$ |
| *ZOIB Mixture Components* | | |
| Zero-Inflation ($\pi_0$) | $0.20 \pm 0.06$ | $0.04 \pm 0.02$ |
| One-Inflation ($q$) | $0.02 \pm 0.02$ | $0.02 \pm 0.01$ |
| Precision ($\varphi$) | $8.47 \pm 2.00$ | $9.78 \pm 1.92$ |

*Note: a:* For the inoculated-twice treatment, scale was estimated via a population mean with row-specific offsets.

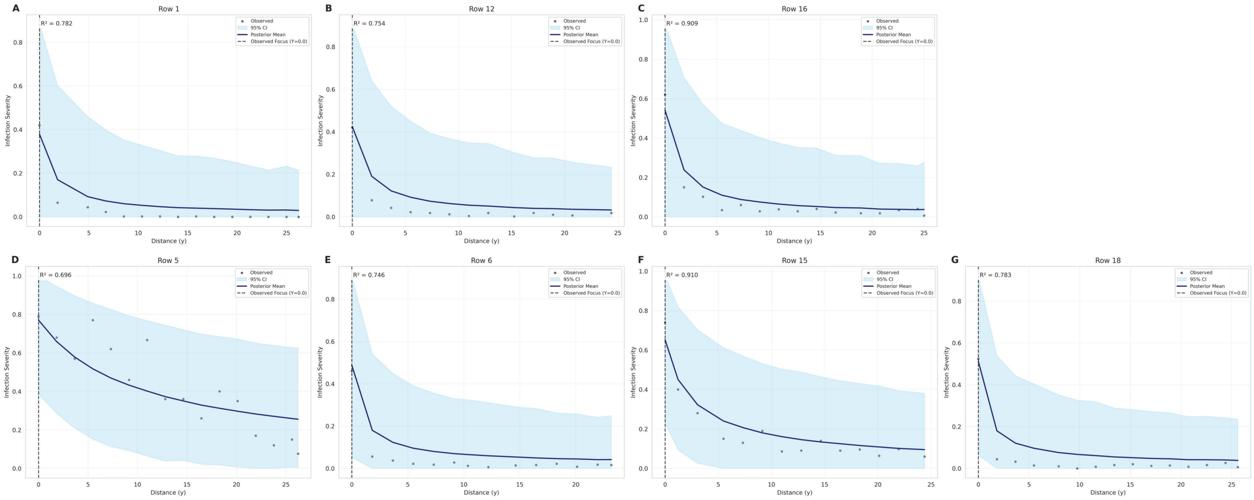


Figure S18: **Spatial validation of posterior predictive distribution for single-focus treatment data under known focus location scenarios.** Panels distinguish between single-inoculation (A-C) and the double-inoculation (D-G) replicates. Primary panels display 1D spatial transects at $x = 0$, comparing observed severity (black points) against the posterior predictive mean (navy line) and 95% credible interval (shaded sky-blue region).

#### Kernel Validation When Known Location - Two Foci Baseline

**Model Selection and Predictive Baselines** To establish a performance baselines for multi-source scenarios, we first evaluated the two-foci models using known focus coordinates. We fitted Exponential, Gaussian, and Power Law kernels to both inoculated-once and inoculated-twice experiments. The Power Law model consistently outperformed alternative kernels, achieving the highest expected log predictive density (Table S19). In these baseline scenarios, the inoculated-once and inoculated-twice treatments yielded overall $R^2$ values of 0.72 and 0.67, respectively. Models that jointly estimated the focus location exhibited only moderately reduction in explanatory capacity, with overall $R^2$ values decreasing to 0.60 and 0.52, respectively (Table S10). Given the low prevalence of structural zeros in the two-foci data, the predictive performance ($R^2$) of the continuous Beta component was nearly identical to the overall model (Table S19).

**Inference Stability and Multi-Source Synergies** A comparison of parameter estimates between known and unknown focus location scenarios reveals a high degree of structural stability (Table S20; Table 9). For the inoculated-once treatment, the inferred global

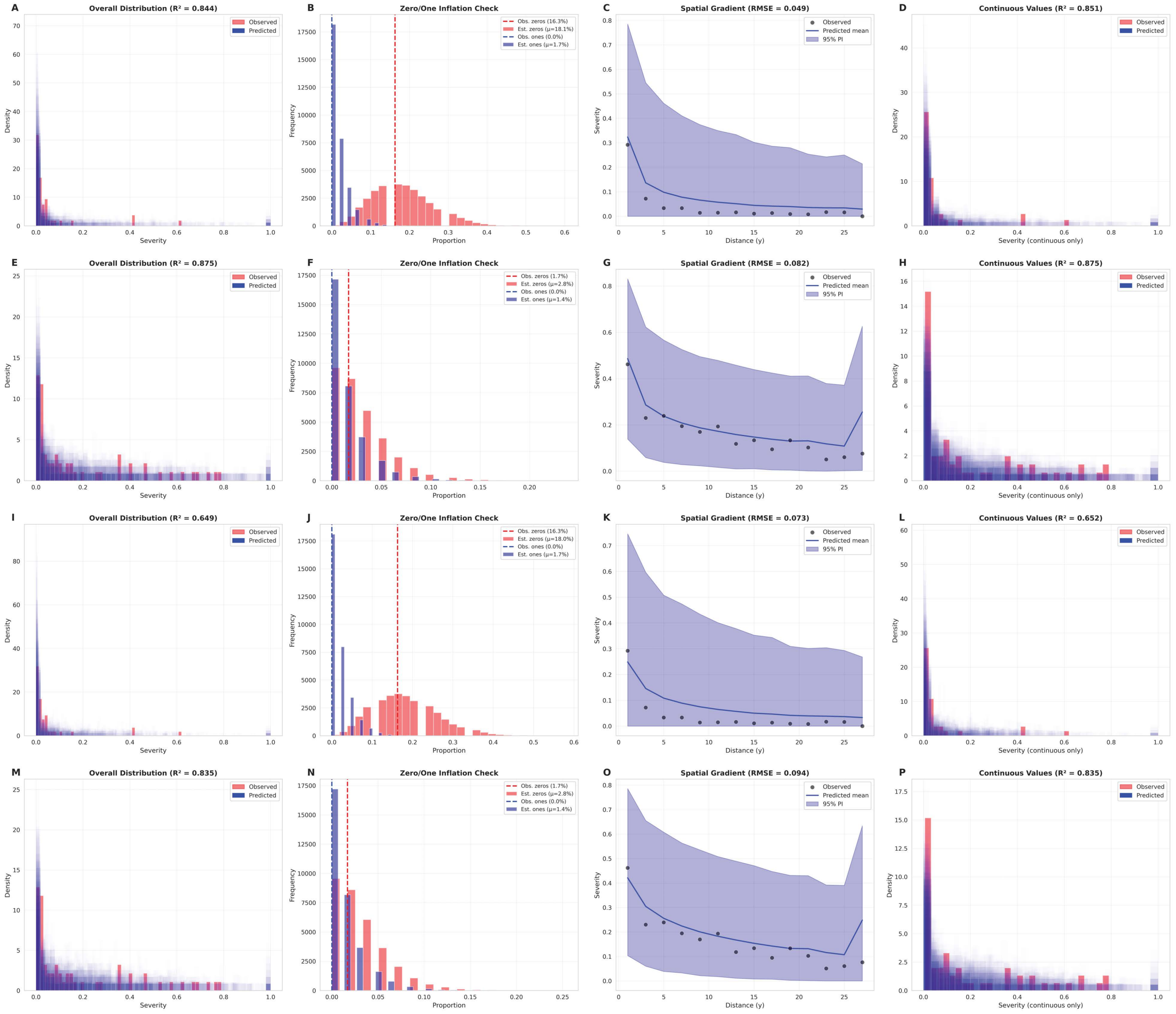


Figure S19: **Posterior predictive checks (PPC) and predictive performance for single-focus treatment data.** The analysis evaluates model performance across known and unknown focus location scenarios for single- and double-inoculation experiments. Rows represent: (A-D) Single-inoculation (known focus location); (E-H) double-inoculation (known focus location); (I-L) Single-inoculation (unknown focus location); and (M-P) double-inoculation (unknown focus location). Columns display: Overall distribution of observed versus predicted severity with associated $R^2$ (A, E, I, M); Zero- and One-inflation checks comparing observed proportions (dashed lines) against model estimates (histograms) (B, F, J, N); Spatial gradients comparing observed binned-mean severity (points; 2-m bins) with the posterior predicted mean (solid line) and 95% credible intervals (shaded region) (C, G, K, O); and predictive distributions and $R^2$ for the continuous Beta component specifically (D, H, L, P).

mean intensities were remarkably similar between known ($f_{z1} = 0.71$ and $f_{z2} = 0.72$) and unknown ($f_{z1} = 0.75$ and $f_{z2} = 0.75$) scenarios. A similar consistency was observed in the inoculated-twice treatment (known:$f_{z1} = 0.77$ and $f_{z2} = 0.81$; unknown:$f_{z1} = 0.78$ and $f_{z2} = 0.81$).

The dispersal parameter also demonstrated high fidelity during joint inference (Table 9). For example, the decay exponent for Focus 1 in the inoculated-once treatment was estimated at 3.59 (known) and 3.37 (unknown), showing that the model can recover the specific physics of the dispersal gradient while searching for the spatial origin. Furthermore, the ZOIB mixture components (zero- and one-inflation) remained nearly identical across known and unknown focus scenarios (for example $\pi_0$ and $q$ were estimated as 0.05 and 0.02, respectively, in the inoculated-once treatment under both scenarios), underscoring the framework's robustness in partitioning healthy plants and fully infected plants from the infection gradient. Visual inspection of posterior predictive patterns confirms this high similarity, with both known and unknown scenarios recovering the observed spatial disease peaks with equal precision (Figure S21; Figure 12M–T; Figure S20). These results suggest that the HiBASIL framework is well-equipped for multi-source inference; rather than increasing the estimation burden, the presence of multiple foci provide a richer spatial signal that the framework successfully disentangle.

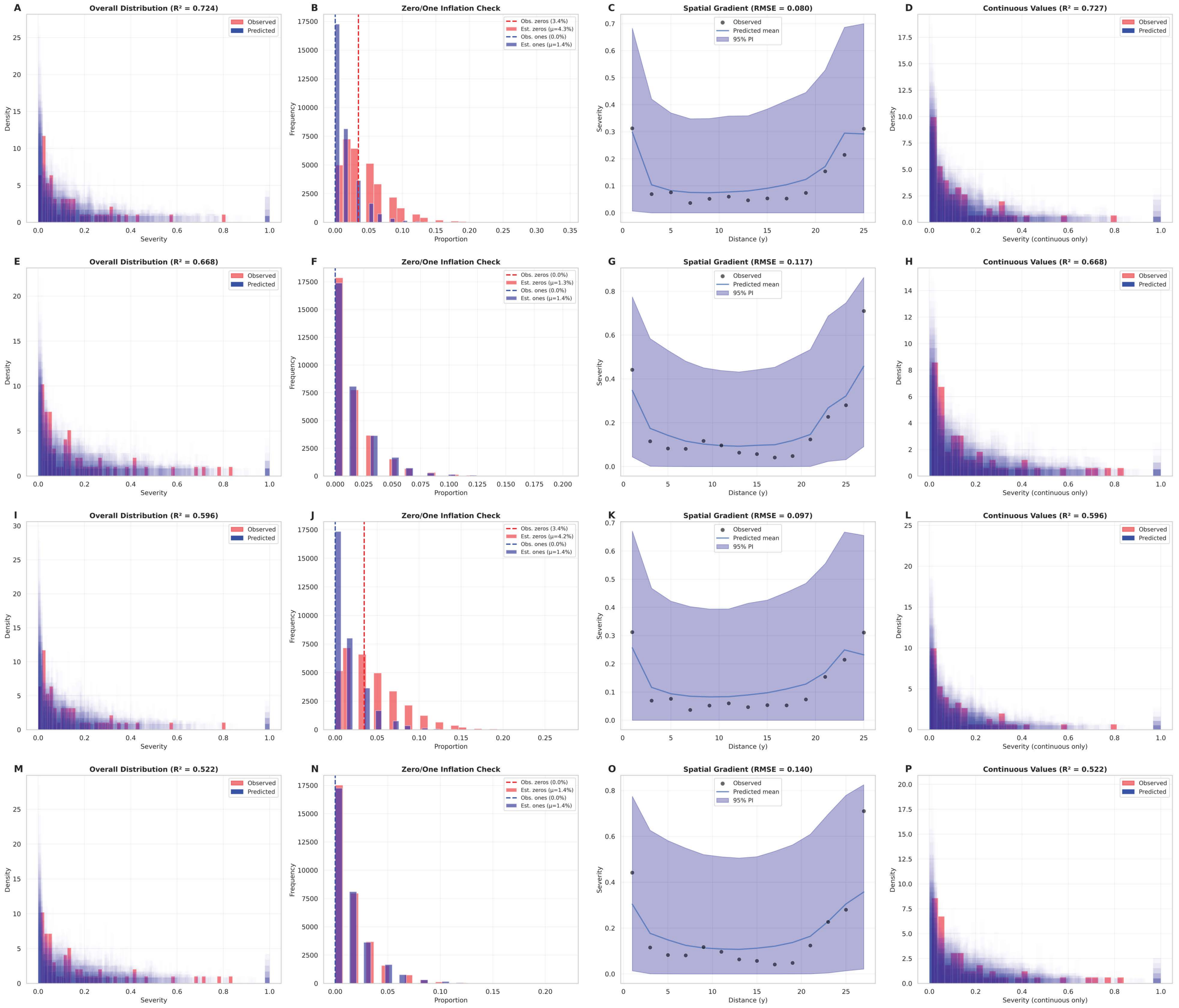


Figure S20: **Posterior predictive checks and predictive performance for two-foci treatment experiments.** Model performance is evaluated across known and unknown focus location scenarios for single- and double-inoculation experiments. Rows represent: (A-D) Single-inoculation with known focus location; (E-H) Double-inoculation with known focus location; (I-L) Single-inoculation with unknown focus location; and (M-P) Double-inoculation with unknown focus location. Columns display: Overall distribution of observed versus predicted severity with associated $R^2$ (A, E, I, M); Zero- and One-inflation checks comparing observed proportions (dashed lines) against model estimates (histograms) (B, F, J, N); Spatial gradients comparing observed binned-mean severity (points; 2-meter bins) with the posterior predicted mean (solid line) and 95% credible intervals (shaded region) (C, G, K, O); and Predictive distributions and $R^2$ for the continuous Beta component (D, H, L, P).

Table S19: Predictive Performance and LOO-CV Results for Two-foci Models for Cucurbit Downy Mildew Field Experiments under **Known Focus Location**.

| | Two-foci (inoculated once) | | | Two-foci (inoculated twice) | | |
|---|---|---|---|---|---|---|
| **Metric** | Exponential | Gaussian | Power Law | Exponential | Gaussian | Power Law |
| overall $R^2$ | 0.645 | 0.413 | **0.724** | 0.631 | 0.442 | **0.668** |
| overall RMSE | 0.090 | 0.116 | **0.080** | 0.123 | 0.151 | **0.117** |
| overall MAE | 0.071 | 0.086 | **0.064** | 0.087 | 0.118 | **0.083** |
| obs_zeros | 0.034 | 0.034 | 0.034 | 0.000 | 0.000 | 0.000 |
| pred_zeros | 0.043 | 0.043 | 0.043 | 0.014 | 0.014 | 0.013 |
| obs_ones | 0.000 | 0.000 | 0.000 | 0.000 | 0.000 | 0.000 |
| pred_ones | 0.014 | 0.014 | 0.014 | 0.014 | 0.014 | 0.014 |
| continuous $R^2$ | 0.645 | 0.415 | **0.727** | 0.631 | 0.442 | **0.668** |
| continuous RMSE | 0.091 | 0.117 | **0.080** | 0.123 | 0.151 | **0.117** |
| n_obs | 58 | 58 | 58 | 59 | 59 | 59 |
| elpd_loo | 59.438 | 49.659 | **63.311** | 57.085 | 47.074 | **58.847** |
| p_loo | 6.368 | 7.112 | **5.990** | **5.652** | 7.016 | 5.723 |
| rank_loo | 1 | 2 | **0** | 1 | 2 | **0** |
| se_d_loo | 1.841 | 3.323 | 0.000 | 1.830 | 2.893 | 0.000 |
| warning | False | False | False | False | False | False |
| pareto_k_bad | 0 | 0 | 0 | 0 | 0 | 0 |

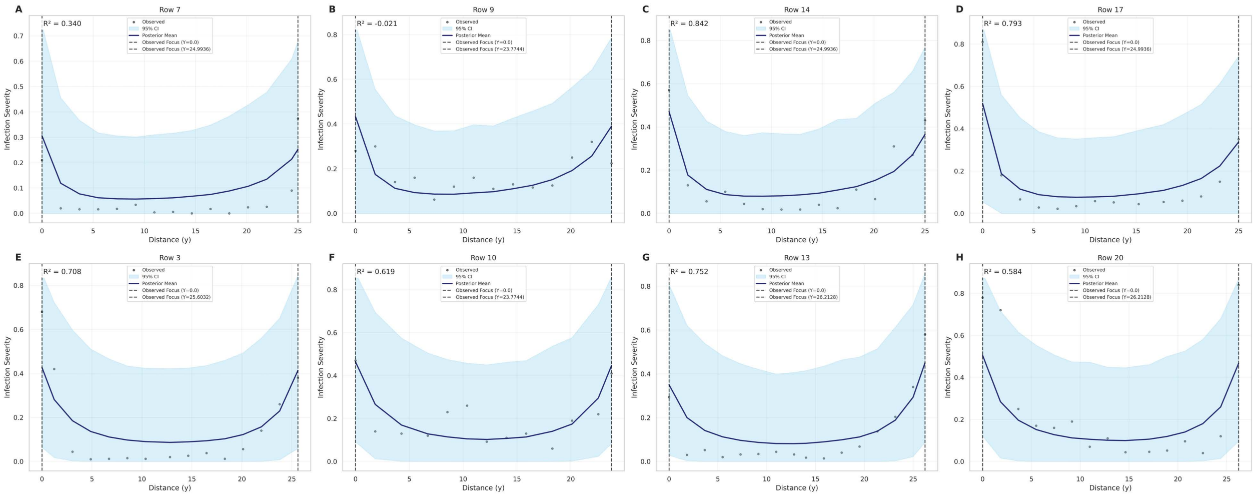


Figure S21: **Spatial validation of posterior predictive distribution for two-foci treatment data under known focus location scenarios.** Panels distinguish between single-inoculation (A-D) and double-inoculation (E-H) replicates. Each main plot display a one-dimensional spatial transects at $x = 0$, showing observed severity (black points) overlaid with the posterior predictive mean (navy line) and 95% credible interval (shaded sky-blue region).

Table S20: Parameter Estimation Results for Two-Foci Models for Cucurbit Downy Mildew Field Experiments under **Known Focus Location** (Fixed Coordinates).

| **Parameter** | **Inoculated once** | **Inoculated twice** |
|---|---|---|
| *Focus Locations* | | |
| Focus 1 & 2 Coordinates | Fixed | Fixed |
| Weight ($w$) | $0.55 \pm 0.07$ | $0.52 \pm 0.06$ |
| *Focus Intensity ($f_z$)* | | |
| **Global Mean** $f_{z1}$ | $0.71 \pm 0.14$ | $0.77 \pm 0.13$ |
| **Global Mean** $f_{z2}$ | $0.72 \pm 0.14$ | $0.81 \pm 0.12$ |
| *Row-Specific $f_{z1}$:* | | |
| $f_{z1}$ - Row 0 | $0.53 \pm 0.20$ | $0.79 \pm 0.16$ |
| $f_{z1}$ - Row 1 | $0.73 \pm 0.19$ | $0.86 \pm 0.13$ |
| $f_{z1}$ - Row 2 | $0.81 \pm 0.16$ | $0.64 \pm 0.19$ |
| $f_{z1}$ - Row 3 | $0.89 \pm 0.12$ | $0.94 \pm 0.08$ |
| *Row-Specific $f_{z2}$:* | | |
| $f_{z2}$ - Row 0 | $0.56 \pm 0.16$ | $0.82 \pm 0.16$ |
| $f_{z2}$ - Row 1 | $0.88 \pm 0.13$ | $0.87 \pm 0.13$ |
| $f_{z2}$ - Row 2 | $0.83 \pm 0.15$ | $0.90 \pm 0.11$ |
| $f_{z2}$ - Row 3 | $0.76 \pm 0.16$ | $0.91 \pm 0.10$ |
| *Dispersal Parameters* | | |
| Exponent 1 | $3.59 \pm 1.29$ | $2.11 \pm 0.92$ |
| Exponent 2 | $1.46 \pm 0.47$ | $2.17 \pm 0.92$ |
| Scale 1 | $4.50 \pm 1.41$ | $5.16 \pm 1.58$ |
| Scale 2 | $5.32 \pm 1.65$ | $5.09 \pm 1.59$ |
| *ZOIB Mixture Components* | | |
| Zero-Inflation ($\pi_0$) | $0.05 \pm 0.03$ | $0.02 \pm 0.02$ |
| One-Inflation ($q$) | $0.02 \pm 0.02$ | $0.01 \pm 0.01$ |
| Precision ($\varphi$) | $9.27 \pm 1.75$ | $7.09 \pm 1.26$ |
